# Genome-wide alterations of uracil distribution patterns in human DNA upon chemotherapeutic treatments

**DOI:** 10.1101/2020.03.04.976977

**Authors:** Hajnalka L. Pálinkás, Angéla Békési, Gergely Róna, Lőrinc Pongor, Gergely Tihanyi, Eszter Holub, Ádám Póti, Carolina Gemma, Simak Ali, Michael J. Morten, Eli Rothenberg, Michele Pagano, Dávid Szüts, Balázs Győrffy, Beáta G. Vértessy

## Abstract

Numerous anti-cancer drugs perturb thymidylate biosynthesis and lead to genomic uracil incorporation contributing to their antiproliferative effect. Still, it is not yet characterized if uracil incorporations have any positional preference. Here, we aimed to uncover genome-wide alterations in uracil pattern upon drug-treatment in human cancer cell-line HCT116. We developed a straightforward U-DNA sequencing method (U-DNA-Seq) that was combined with *in situ* super-resolution imaging. Using a novel robust analysis pipeline, we found broad regions with elevated probability of uracil occurrence both in treated and non-treated cells. Correlation with chromatin markers and other genomic features shows that non-treated cells possess uracil in the late replicating constitutive heterochromatic regions, while drug treatment induced a shift of incorporated uracil towards more active/functional segments. Data were corroborated by colocalization studies via dSTORM microscopy. This approach can also be applied to study the dynamic *spatio-temporal* nature of genomic uracil.

## INTRODUCTION

The thymine analogue uracil is one of the most frequent non-canonical bases in DNA appearing either by thymine replacing misincorporation or as a product of spontaneous or enzymatic cytosine deamination reaction (Krokan, Drabløs, & Slupphaug, 2002). Consequently, uracil in DNA is usually recognized as an error that is efficiently repaired by the multistep base excision repair (BER) pathway initiated by uracil-DNA glycosylases (UDGs) (Krokan & Bjørås, 2013; Wallace, 2014). In other respects, uracil in DNA is known to be involved in several physiological processes (e.g. antibody maturation (Liu & Schatz, 2009; Maul & Gearhart, 2010, 2014; Xu, Zan, Pone, Mai, & Casali, 2012), antiviral response (Burns, Leonard, & Harris, 2015; Stenglein, Burns, Li, Lengyel, & Harris, 2010), insect development (Horváth, Békési, Muha, Erdélyi, & Vértessy, 2013; Muha et al., 2012)), however, the exact mechanism and regulation of uracil-DNA metabolism including the roles of UDGs need to be elucidated. There are four known members of the UDG family in humans: (i) the most active uracil-DNA glycosylase encoded by the *ung* gene (UNG1 mitochondrial and UNG2 nuclear isoform), (ii) the single-strand selective monofunctional uracil-DNA glycosylase 1 (SMUG1), (iii) thymine DNA glycosylase (TDG specialized for repair of T:G and U:G) and (iv) methyl CpG binding domain protein (MBD4 repairs U:G) (Visnes et al., 2009). UNG2 removes most of the genomic uracil from both single- and double-stranded DNA regardless of the uracil originating from mutagenic cytosine deamination or thymine replacing misincorporation (Kavli et al., 2002).

Thymine replacing uracil misincorporation is normally prevented by the tight regulation of the cellular dUTP/dTTP ratio maintained by two enzymes, the dUTPase and the thymidylate synthase. The dUTPase enzyme (Vértessy & Tóth, 2009) removes dUTP from the cellular pool. Lack or inhibition of dUTPase leads to increased dUTP levels and under such conditions, DNA polymerases readily incorporate uracil opposite to adenine. Similarly, several anticancer drugs (such as 5-fluorouracil (5-FU), 5-fluoro-2’-deoxyuridine (5FdUR), capecitabine, methotrexate, raltitrexed (RTX), pemetrexed) target the *de novo* thymidylate synthesis pathway via thymidylate synthase inhibition to perturb the tightly regulated dUTP/dTTP ratio, eventually triggering thymineless cell death (Blackledge, 1998; Wilson, Danenberg, Johnston, Lenz, & Ladner, 2014). Although the exact molecular mechanism is not yet fully understood, massive uracil misincorporation, hyperactivity of the repair process and/or stalling of the replication fork are all suggested to be involved in the process (Khodursky, Guzmán, & Hanawalt, 2015; Ostrer, Hamann, & Khodursky, 2015). UNG has been suggested to play a key role in this mechanism, as being responsible for the initiating step in uracil removal that may lead to futile cycles if the cellular dUTP/dTTP ratio is elevated. A quantitative insight into the magnitude and the pattern of uracil incorporation into genomic DNA as induced by these chemotherapeutic treatments is expected to contribute to a better understanding of the cell death mechanism induced by the respective drugs.

Direct observation of the uracil moieties incorporated upon drug treatments have been hampered by the efficient and fast action of UNG. To overcome this problem, we wished to counteract the action of UNG in human cells *via* introduction of the UNG inhibitor, UGI (Luo, Walla, & Wyatt, 2008), into the cellular milieu. By this approach we aimed to reveal the nascent pattern of uracil moieties in DNA induced by perturbation of thymidylate metabolism both using genome-wide uracil-specific sequencing and *in situ* cellular imaging of uracils within human genomic DNA. Previously, we designed a uracil-DNA (U-DNA) sensor tailored from an inactive mutant of human UNG2 that was successfully applied in semi-quantitative dot blot analysis and direct immunocytochemistry (Róna et al., 2016). Some additional approaches have also been published to detect uracil-DNA within its genomic context such as i) techniques focusing on specific, well-defined regions of the genome (qPCR (Horváth & Vértessy, 2010) and 3D-PCR (Suspène, Henry, Guillot, Wain-Hobson, & Vartanian, 2005)), ii) techniques applicable to smaller sized genomes only (Excision-seq (Bryan, Ransom, Adane, York, & Hesselberth, 2014) and UPD-seq (Sakhtemani et al., 2019)), and iii) techniques requiring labour-intensive isolation and multistep processing of genomic DNA samples (dU-seq (Shu et al., 2018)).

Here, we employ the U-DNA sensor in a DNA-IP-seq-like approach (termed as U-DNA-Seq) and develop a robust bioinformatic pipeline specifically designed for reliable interpretation of next generation sequencing data for genome-wide distribution of uracil. We selected two drugs, RTX (raltitrexed, or tomudex) and 5FdUR that perturb thymidylate biosynthesis with different modes of action and analysed their effects on genomic uracil distribution. These two drugs are frequently applied in treatment of colon cancers, therefore we chose a human colon carcinoma cell line, HCT116 as a well-established and relevant cellular model. We show that drug treatment led to increased probability of uracil incorporation into more active chromatin regions in HCT116 cells expressing the UNG inhibitor protein UGI. In contrast, uracil was rather restricted to constitutive heterochromatic regions both in wild type cells and in non-treated UGI-expressing cells. Moreover, we further developed the U-DNA sensor-based staining method (Róna et al., 2016) that now uniquely allows *in situ* microscopic visualization of uracil in human genomic DNA. Confocal and super-resolution microscopy images and colocalization measurements strengthened the sequencing-based distribution patterns.

## MATERIALS AND METHODS

### Plasmid constructs and cloning of the FLAG-ΔUNG-SNAP construct

The pLGC-hUGI/EGFP plasmid was kindly provided by Michael D. Wyatt (South Carolina College of Pharmacy, University of South Carolina, US). Generation of catalytically inactive U-DNA sensor proteins (1xFLAG-ΔUNG, 3xFLAG-ΔUNG, FLAG-ΔUNG-DsRed) was described previously (Róna et al., 2016). pSNAPf (New England Biolabs (NEB), Ipswich, Massachusetts (MA), US) was PCR amplified with primers SNAP-Fw (5’ – TAA TGG TAC CGC GGG CCC GGG ATC CAC CGG TCG CCA CCA TGG ACA AAG ACT GCG AAA TG – 3’) and SNAP-Rev (5’ – ATA TCT CGA GGC CTG CAG GAC CCA GCC CAG G – 3’). The resulting fragments were digested by KpnI and XhoI and ligated into the KpnI/XhoI sites of the plasmid construct FLAG-ΔUNG-DsRed (in a pET-20b vector) yielding the FLAG-ΔUNG-SNAP construct. Scheme of the used constructs is shown in Supplementary Figure S8A. Primers used in this study were synthesized by Sigma-Aldrich (St. Louis, Missouri, US) and all constructs were verified by sequencing at Microsynth Seqlab GmbH (Göttingen, Germany). All UNG constructs were expressed in the *Escherichia coli* BL21(DE3) *ung-151* strain and purified using Ni-NTA affinity resin (Qiagen, Hilden Germany) as described previously (Róna et al., 2016).

### DNA isolation and purification

pEGFP-N1 plasmid (Clontech, Mountain View, California, US) was transformed into XL1-Blue [*dut+, ung+*] (Stratagene, San Diego, California (CA), US) or CJ236 [*dut-, ung-*] (NEB) *E. coli* competent cells. Cell cultures were grown for 16 h in Luria broth (LB) media supplemented with 50 μg/ml kanamycin at 37°C. Plasmids used in this study were purified using PureYield™ Plasmid Midiprep Kit (Promega, Madison, Wisconsin, US) according to the instructions of the manufacturer. XL1-Blue and CJ236 *E. coli* strains were propagated in LB media at 37°C and were harvested at log phase. Genomic DNA of bacterial samples as well as eukaryote cells was purified using the Quick-DNA™ Miniprep Plus Kit (Zymo Research, Irvine, California, US) using the recommendations of the manufacturer.

### Cell culture, transient transfection and treatment of cells

The 293T cell line was a generous gift of Yvonne Jones (Cancer Research UK, Oxford, UK). The HCT116 and the K562 cell lines were purchased from the European Collection of Cell Cultures (ECACC, Salisbury, UK). 293T cells were grown in Dulbecco’s modified Eagle’s medium (Gibco, Life Technologies, Carlsbad, CA, US), while HCT116 and K562 cells were maintained in McCoy’s 5A medium (Gibco) and RPMI 1640 (GlutaMAX™ Supplement, HEPES) Medium (Gibco), respectively. Media was supplemented with 50 μg/ml Penicillin-Streptomycin (Gibco) and 10% fetal bovine serum (Gibco). Cells were cultured at 37°C in a humidified incubator with 5% CO_2_ atmosphere. HCT116 cells were transfected with FuGENE HD (Promega) according to the manufacturer’s recommendation. For immunocytochemistry, HCT116 cells were transfected with normal pEGFP-N1 (purified from XL1-Blue [*dut+, ung+*] *E. coli* cells) or uracil-rich pEGFP-N1 (purified from CJ236 [*dut−, ung−*] *E. coli* cells) vector as described previously (Róna et al., 2016). Forty hours after transfection with UGI expressing vectors, transiently transfected cells were grown for an additional 48 h either in the absence or presence of 20 μM 5FdUR (Sigma) before collecting them for genomic DNA purification.

### Generation of UGI-expressing stable cell line

Retroviral packaging and stable cell line generation was done as described in (Rona et al., 2018). Briefly, 293T cells (1.5 × 10^6^ cells in T25 tissue culture flasks) were transfected with 1.5 μg pLGC-hUGI/EGFP, 0.5 μg pCMV-VSV-G envelope and 0.5 μg pGP packaging plasmids using Lipofectamine 3000 transfection reagent (Invitrogen, Carlsbad, CA, US) according to the manufacturer’s recommendation. The supernatant, containing lentiviral particles was collected and filtered through a 0.45 μm filter (Merck Millipore, Burlington, MA, US) 36 h after the transfection. Successfully transduced HCT116 were collected by FACS sorting for GFP-positive cells using a BD FACSAria III Cell sorter (BD Biosciences, San Jose, CA, US). UGI-expressing cells were treated with 20 μM 5FdUR or 100 nM RTX (Sigma) for 48 h before fixation for immunocytochemistry or collecting them for genomic DNA purification described above.

### Dot blot measurements and analysis for quantification of U-DNA

Detection of the genomic uracil content by dot blot measurements were carried out using 3xFLAG-ΔUNG construct, as described earlier (Róna et al., 2016). Dot blot assay was used for measuring genomic uracil levels of non-treated and drug (5FdUR or RTX) treated HCT116 cells expressing UGI (Supplementary Figure S1B), or to confirm the successful enrichment of uracil containing DNA (Figure 1B) and also to compare uracil recognition specificity of the FLAG-ΔUNG-DsRed and FLAG-ΔUNG-SNAP constructs (Supplementary Figure S8B). Densitometry was done using ImageJ (Fiji) software (National Institutes of Health, US). Analysis of the data and the calculation of the number of deoxyuridine nucleotides in the unknown genomic DNA was described before (Molnár, Marton, Izrael, Pálinkás, & Vértessy, 2018; Róna et al., 2016). Briefly, the number of uracil/million bases in the unknown samples were determined by interpolating their normalized intensities to the calibration curve of the standard. Statistical analysis of dot blot (Supplementary Figure S1C) was carried out by Microsoft Excel using the non-parametric two-sided Mann-Whitney U test. Differences were considered statistically significant at p < 0.005. Data presented are representative of six independent datasets (n = 6).

**Figure 1.**
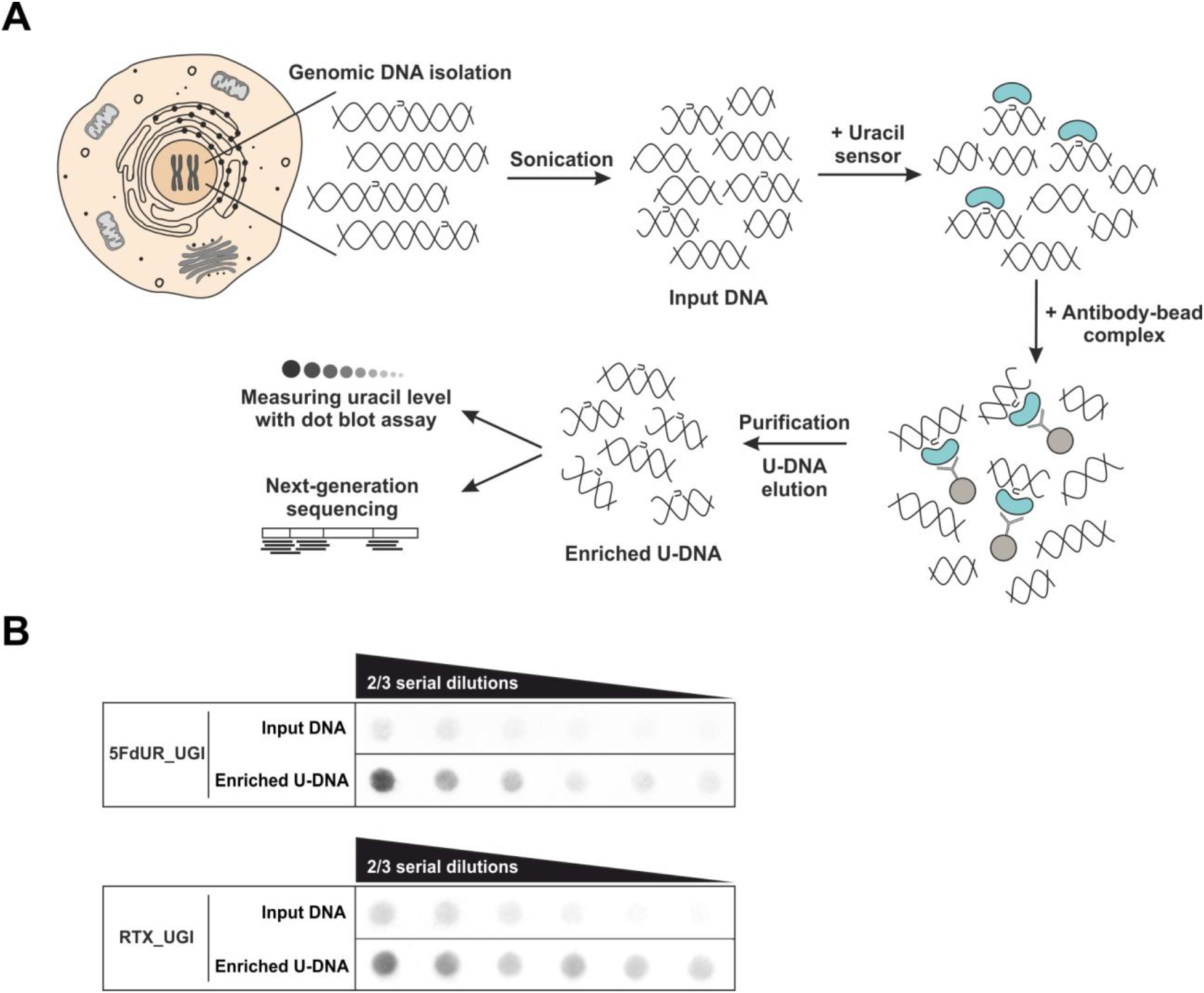
U-DNA-Seq provides genome-wide mapping of uracil-DNA distribution. **(A)** Schematic image of the novel U-DNA immunoprecipitation and sequencing method (U-DNA-Seq). After sonication, enrichment of the fragmented U-DNA was carried out by the 1xFLAG-ΔUNG sensor construct followed by pull-down with anti-FLAG agarose beads. U-DNA enrichment compared to input DNA was confirmed by dot blot assay before samples were subjected to NGS. **(B)** Immunoprecipitation led to elevated uracil levels in enriched U-DNA samples compared to input DNA in case of both 5FdUR (5FdUR_UGI) and RTX (RTX_UGI) treated samples. In case of the given treatment, the same amount of DNA was loaded from input and enriched U-DNA samples providing correct visual comparison of the dots. Two-third serial dilutions were applied.

### DNA immunoprecipitation

After 48h treatment, the surface attached cells were harvested. Genomic DNA was purified by Quick-DNA− Miniprep Plus Kit (Zymo Research) and eluted in nuclease-free water. 12 μg of genomic DNA was sonicated into fragments ranging between 100 and 500 basepairs (bp) (checked by agarose gel electrophoresis) with a BioRuptor (Diagenode, Liège, Belgium). 25% of the samples was saved as input, and the remaining DNA was resuspended in the following IP buffer: 30 mM Tris-HCl, pH = 7.4, 140 mM NaCl, 0.01% Tween-20, 1 mM ethylenediaminetetraacetic acid (EDTA), 15 mM β-mercaptoethanol, 1 mM phenylmethylsulfonyl fluoride, 5 mM benzamidine. Immunoprecipitations were carried out with 15 μg of 1xFLAG-ΔUNG construct for 2.5 h at room temperature with constant rotation. Anti-FLAG M2 agarose beads (Sigma) were equilibrated in IP buffer and then added to the IP mixture for 16 h at 4°C with constant rotation. Beads were washed three times for 10 min in IP buffer and resuspended in elution buffer containing 1% sodium dodecyl sulphate (SDS), 0.1 M NaHCO_3_. Elution of uracil sensor protein binding U-DNA was done by vortexing for 5 min with an additional incubation for 20 min with constant rotation. After centrifugation (13000 rpm for 3 min), supernatant was transferred to clean tubes. This procedure was repeated with the same amount of elution buffer and protein/DNA eluted complexes were combined in the same tube. Samples were incubated with 150 μg/ml RNAse A (Epicentre, Paris, France) for 30 min, followed by the addition of 500 μg/ml Proteinase K (Sigma) for 1 h at 37°C for removal of RNA and proteins. Immunoprecipitated DNA was purified with NucleoSpin Gel and PCR Clean-up Kit (MACHEREY-NAGEL GmbH & Co. KG, Düren, Germany) according to the manufacturer’s instructions. Densitometry analysis of agarose gel was done using ImageJ (Fiji) software for concentration calculation of fragmented DNA. Enrichment of uracil in the DNA samples was examined by dot blot assay. DNA libraries were created from the samples and then subjected to next-generation sequencing (NGS). Scheme of U-DNA-Seq is shown in Figure 1A.

### High-throughput DNA sequencing and data analysis

Sequencing of input and enriched U-DNA samples were done on two independent biological replicates at BGI (China) generating 100 bp paired-end reads (PE) on a HiSeq 4000 instrument or at Novogene (China) using the Novaseq 6000 platform resulting in 150 bp PE reads. Analysis pipeline is summarized in Figure 2, and details including the applied command lines and scripts are found in the Supplementary Material. Sequencing reads were aligned to the GRCh38 human reference genome (version GRCh38.d1.vd1) (Jensen, Ferretti, Grossman, & Staudt, 2017) using BWA (version 0.7.17) (Li & Durbin, 2010). Aligned reads were converted to BAM format and sorted using samtools (version 1.9) (Li et al., 2009). Duplicate reads were marked using Picard Tools (version 1.95). As a part of pre-processing, blacklisting and filtering of ambiguously mapped reads were also performed (cf. Supplementary Material and Supplementary Figure S3). For data processing, to derive uracil distribution signal, first, normalized coverage signals were calculated and smoothened using bamCoverage from the deepTools package (Ramírez et al., 2016), which resulted in genome-scaled coverage tracks in bigWig format. Then, log2 ratio of the coverage tracks (enriched / input) were calculated with bigWigCompare. These bigwig files were compared using the multiBigWigSummary, Pearson correlations were calculated using the plotCorrelation tools also from the deepTools package (Figure 3B). From the log2 ratio tracks, interval (bed) files were derived using reasonable thresholds (for details see the Supplementary Material and Supplementary Table S3A). Log2 ratio signal distribution (Figure 3C) was calculated using R. Peaks of coverage were also called using the MACS2 with broad option (version 2.1.2), a standard tool in chromatin marker ChIP-seq data analysis (Zhang et al., 2008). Results of peak calling and the regions derived from the log2 ratio tracks were compared (Supplementary Table S3). Hereafter, the two terms ‘peak’ and ‘region’ will be consequently applied for the results of the two approaches, respectively. Colocalization analysis of identified uracil enriched regions with other ChIP-seq and DNA accessibility data was performed on a dataset containing HCT116 specific or other relevant data only (for details see Supplementary Material) using GIGGLE search tool (Layer et al., 2018). To plot results of GIGGLE search, OriginPro 8.6 was used (Figure 4A). Measuring overlaps with other genomic features (Figure 4B) was done using bedtools annotate tool (Quinlan & Hall, 2010) as it is described in Supplementary Material. Correlation analysis between uracil enrichment and replication timing (Figure 4C and Supplementary Figure S11C) was done using R as it is described in Supplementary Material. Sequencing data were visualized (Figure 3A, Supplementary Figures S3, S5, S6, S10A, S11A) using the IGV browser (Thorvaldsdóttir, Robinson, & Mesirov, 2013).

**Figure 2.**
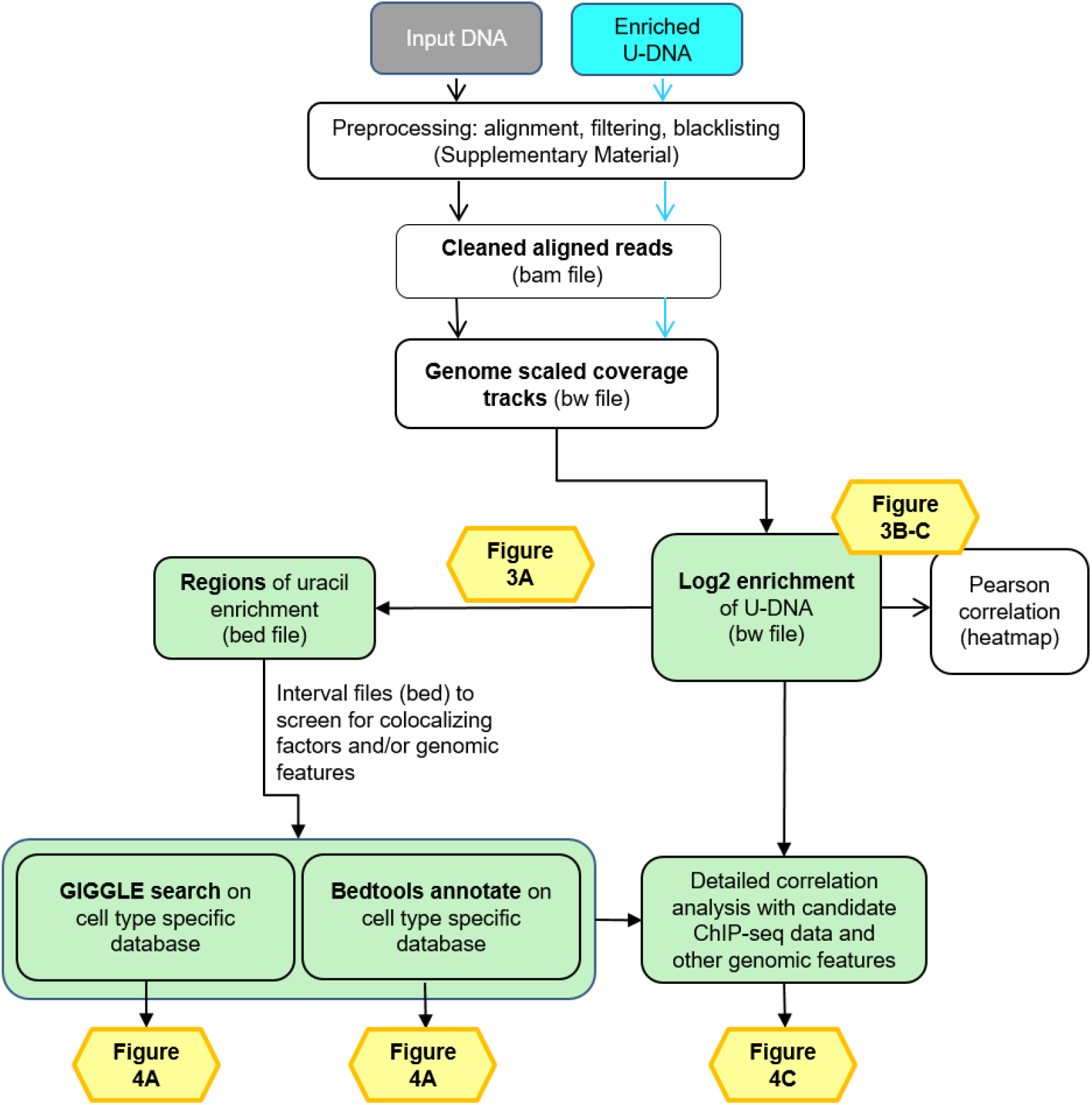
Data analysis pipeline. Both input and enriched U-DNA samples were pre-processed the same way: initial trimming and alignment were followed by filtering for uniquely mapped reads and blacklisting of regions suffering from alignment artefacts, resulting in cleaned aligned reads in the format of bam files. The key steps of our proposed data processing are 1) calculation of genome scaled coverage tracks (bigwig/bw files), 2) calculation of log2 (enriched coverage / input coverage) ratio tracks (bigwig/bw files), 3) extraction of interval (bed) files of uracil enriched regions from the corresponding log2 ratio tracks. To correlate the uracil enrichment profiles with other published data, first quick screens using interval files were done, and then detailed correlation analysis with a promising candidate of colocalizing genomic features was performed using coverage track files. GIGGLE search (Layer et al., 2018) and bedtools annotate (Quinlan & Hall, 2010) were used for scoring the similarities between query uracil-DNA and the database interval files. Figures corresponding to the different analysis steps are also indicated. A more detailed pipeline is shown in Supplementary Figure S2, and the full methodology is described in the Supplementary Material.

**Figure 3.**
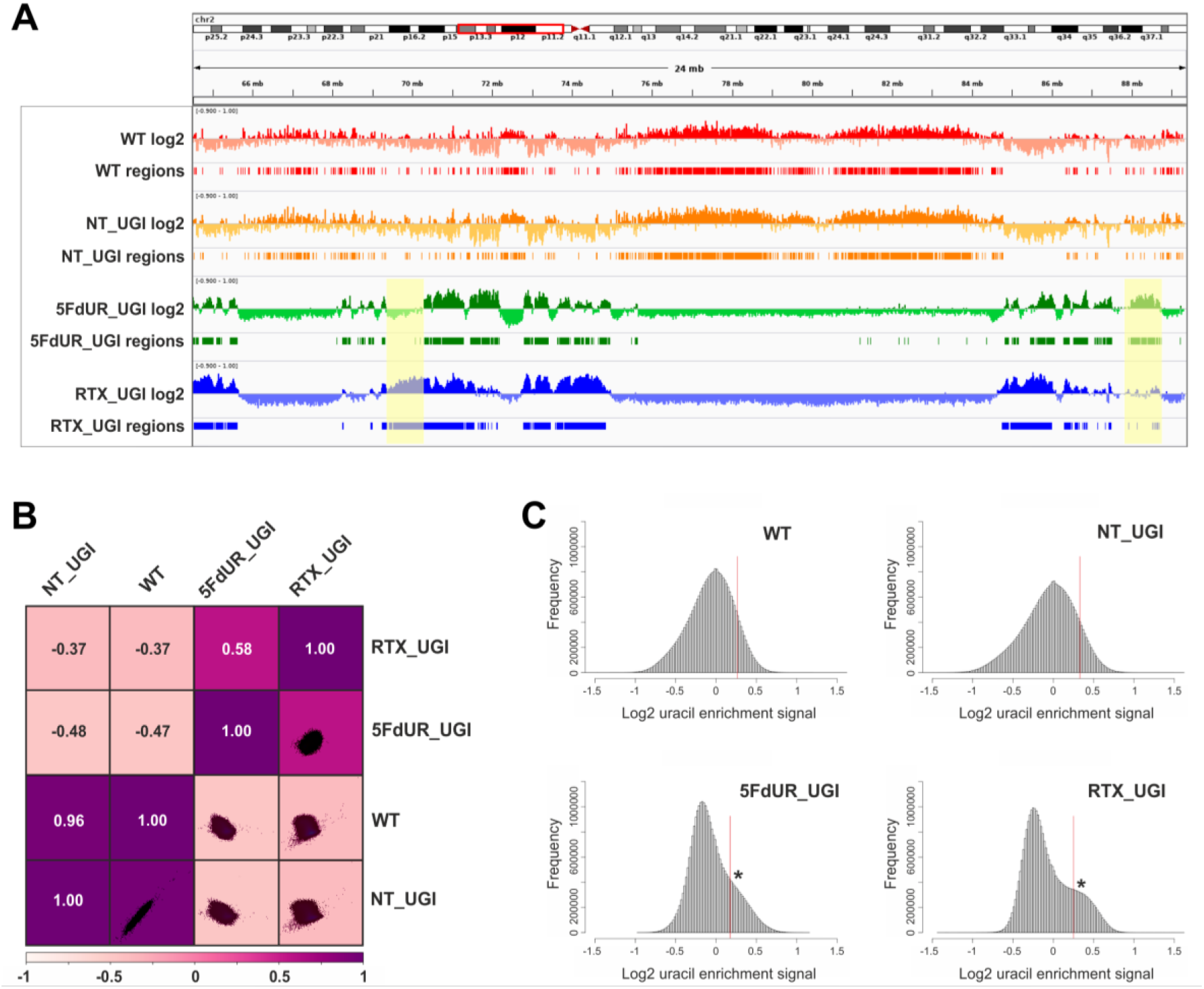
Comparison of processed U-DNA-Seq data among samples. **(A)** Representative IGV view on the log2 ratio and the derived regions of uracil enrichment (two replicates for each sample were merged). Log2 ratio signal tracks of enriched versus input coverage (log2, upper track) and derived regions of uracil enrichment (regions, bottom track) for non-treated: wild type (WT, red) and UGI-expressing (NT_UGI, orange); and for treated: with 5FdUR (5FdUR_UGI, green) or raltitrexed (RTX_UGI, blue) HCT116 samples are shown in genomic segment (chr2:64,500,000-89,500,001). Differences between treated and non-treated samples are clearly visible. Furthermore, 5FdUR and RTX treatments caused similar but not identical uracil enrichment profile (differences are highlighted with yellow background). **(B)** Comparison of log2 uracil enrichment profiles among samples was performed using multiBigWigSummary (deepTools) and Pearson correlation were plotted using plotCorrelation (deepTools). A heatmap combined with scatterplots is shown for the four samples. **(C)** Histograms of log2 ratio profiles were calculated and plotted using R. A sub-population of data bins with elevated log2 uracil enrichment signal is clearly visible (indicated with asterisk) in case of drug-treated samples, where high uracil incorporation was detected (cf. Supplementary Figure S1B-C). Thresholds applied in determination of uracil enriched regions are indicated with red line (cf. Supplementary Table S3A).

**Figure 4.**
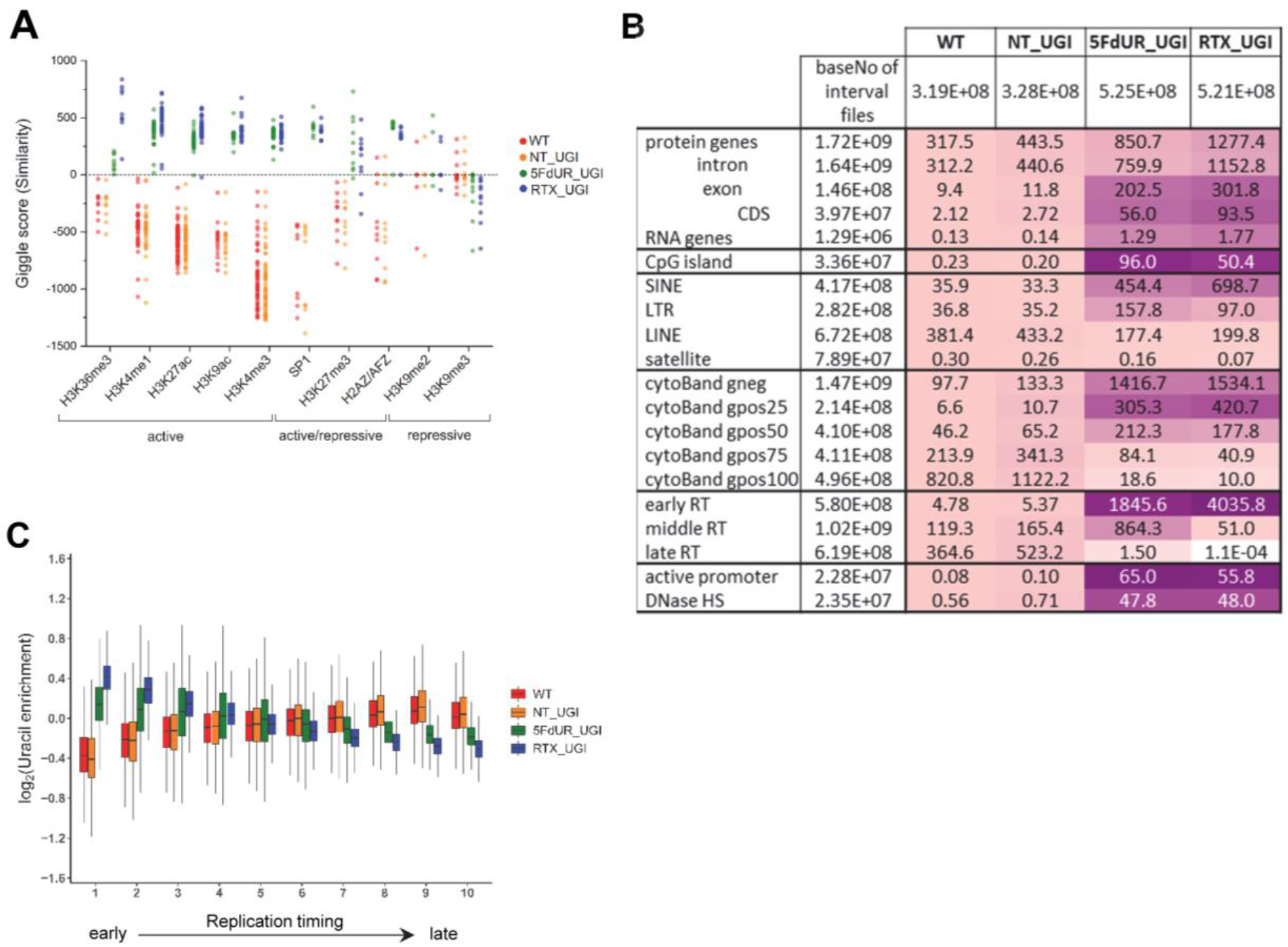
Characterization of U-DNA enrichment patterns. **(A)** Top hs from GIGGLE search on HCT116 specific dataset. GIGGLE search was performed with interval (bed) files of uracil enriched regions on a set of HCT116 related ChIP-seq and DIP-seq experiment data (for details see the Supplementary Material). Factors corresponding to the top 10 hits for each sample were selected. GIGGLE scores between all four samples and all experiments corresponding to these factors were plotted excluding CNOT3, H2B, H3K27me1/2 where data were not informative (data are found in Supplementary Material Appendix 2). Histone marks and the only transcription factor, SP1 are categorized depending on their occurrence in transcriptionally active or repressive regions. Notably, some of them have plastic behaviour allowing either transcriptionally active or repressive function. U-DNA-Seq samples are as follows: non-treated wild type (WT, red), non-treated UGI-expressing (NT_UGI, orange), 5FdUR treated UGI-expressing (5FdUR_UGI, green), and RTX treated UGI-expressing (RTX_UGI, blue) HCT116 cells. **(B)** Correlation with genomic features. Interval (bed) files of genomic features were obtained from UCSC, Ensembl, and ReplicationDomain databases (for details see the Supplementary Material), and correlation with interval files of uracil regions were analysed using bedtools annotate software. Numbers of overlapping basepairs were summarized for each pair of interval files, and scores were calculated according the formula: (baseNo_overlap/baseNo_sample_file) * (baseNo_overlap/baseNo_feature_file) * 10000. Heatmaps were created based on fold increase of the scores compared to the corresponding WT scores. Sizes of interval files in number of basepairs are also given in the second column and the second line. Upon drug treatments, a clear shift from non-coding / heterochromatic / late replicated segments towards more active / coding / euchromatic / early replicated segments can be seen. CDS, coding sequence; SINE, short interspersed element; LTR, long terminal repeat; LINE, long interspersed element; cytoBand, cytogenic chromosome band negatively (gneg) or positively (gpos) stained by Giemsa; RT, replication timing; DNaseHS, DNase hypersensitive site. **(C)** Correlation analysis with replication timing. Replication timing data (bigWig files with 5000 bp binsize) specific for HCT116 were downloaded from ReplicationDomain database. Data bins were distributed to 10 equal size groups according to replication timing from early to late. Then log2 uracil enrichment signals for these data bin groups were plotted for each sample using R.

### Immunofluorescent staining of uracil residues

Detection of uracil residues was done in extrachromosomal plasmids after transfection (Supplementary Figure S8C) or in genomic DNA of HCT116 cells (Figure 5–7). Staining of extrachromosomal DNA was done as described previously (Róna et al., 2016) with minor modifications for comparison of FLAG-ΔUNG-DsRed or FLAG-ΔUNG-SNAP sensor constructs. Briefly, uracil residues were visualized by applying 1.5 μg/ml of the FLAG-ΔUNG-DsRed or the FLAG-ΔUNG-SNAP, and then primary (anti-FLAG M2 antibody (1:10000, Sigma) and secondary antibodies (Alexa 488 (1:1000, Molecular Probes, Eugene, Oregon, US)). For immunofluorescent staining of genomic uracil residues, control or HCT116 cells stably expressing UGI were seeded onto 24-well plates containing cover glasses or onto μ-Slides (or their glass bottom derivative) (ibidi GmbH, Germany) suitable for use in STED and single molecule applications, and treated as indicated. In case of dSTORM imaging, coverslips were coated with poly-D-lysine (Merck Millipore) before seeding the cells. Sub-confluent cultures of cells were fixed using 4% paraformaldehyde (PFA, pH = 7.4 in phosphate-buffered saline (PBS)) or Carnoy’s fixative (ethanol: acetic acid: chloroform = 6:3:1) for 15 min. In case of dSTORM imaging, cells were pre-extracted with ice-cold CSK buffer (10 mM PIPES, pH = 6.8, 100 mM NaCl, 300 mM sucrose, 1 mM EGTA, 3 mM MgCl_2_, 0.25% Triton X-100) containing protease and phosphatase inhibitor tablets (Roche, Basel, Switzerland) for 5 min before PFA fixation. After washing or rehydration steps (1:1 ethanol:PBS, 3:7 ethanol:PBS, PBS), epitope unmasking was done by applying 2 M HCl, 0.5% Triton X-100 for 30 min. DNA denaturation with HCl was required in order to increase DNA accessibility for efficient staining and to eliminate any potential interaction between the overexpressed UGI and the applied UNG sensor construct. After neutralization with 0.1 M Na_2_B_4_O_7_ (pH = 8.5) for 5 min followed by PBS washes, cells were incubated in blocking solution I (TBS-T (50 mM Tris-HCl, pH = 7.4, 2.7 mM KCl, 137 mM NaCl, 0.05% Triton X-100) containing 5% non-fat dried milk) for 15 min, followed by incubation in blocking buffer I supplemented with 200 μg/ml salmon sperm DNA (Invitrogen) for an additional 45 min. Uracil residues were visualized by applying 4 μg/ml of the FLAG-ΔUNG-SNAP construct for 1 h in blocking buffer I with 200 μg/ml salmon sperm DNA at room temperature. After several washing steps with TBS-T containing 200 μg/ml salmon sperm DNA, primary, then secondary antibodies were operated in blocking buffer II (5% fetal goat serum (FGS), 3% fetal bovine serum albumin (BSA) and 0.05% Triton X-100 in PBS). Anti-FLAG M2 antibody (1:10000, Sigma), then Alexa 488 conjugated secondary antibody (1:1000, Molecular Probes) was applied for 1 h in blocking buffer II, enabling visualization of FLAG epitope. SNAP-tag substrates were also used to label SNAP-tag fusion proteins when FLAG-ΔUNG-SNAP was applied as the uracil sensor protein. Cells were labelled with 2.5 μM (0.5 μM for dSTORM imaging) SNAP-Surface Alexa Fluor 546 or 647 (indicated as SNAP546 and SNAP647 in this study) (NEB) for 20 min and optionally counterstained with 1 μg/ml DAPI (4’,6-diamidino-2-phenylindole, Sigma) nucleic acid stain, followed by several PBS washing steps before embedding in FluorSave− Reagent (Calbiochem, Merck Millipore). For labelling of histone markers, anti-H3K36me3 (1:8000, CST (Danvers, MA, US), cat.no.: 4909T) or anti-H3K27me3 (1:6000, CST, cat.no.: 9733T) primary antibodies were used, then visualized by Alexa 568 conjugated secondary antibody (1:10000, Molecular Probes) in dSTORM or Alexa 555 conjugated secondary antibody (1:2000, Molecular Probes) in confocal imaging.

**Figure 5.**
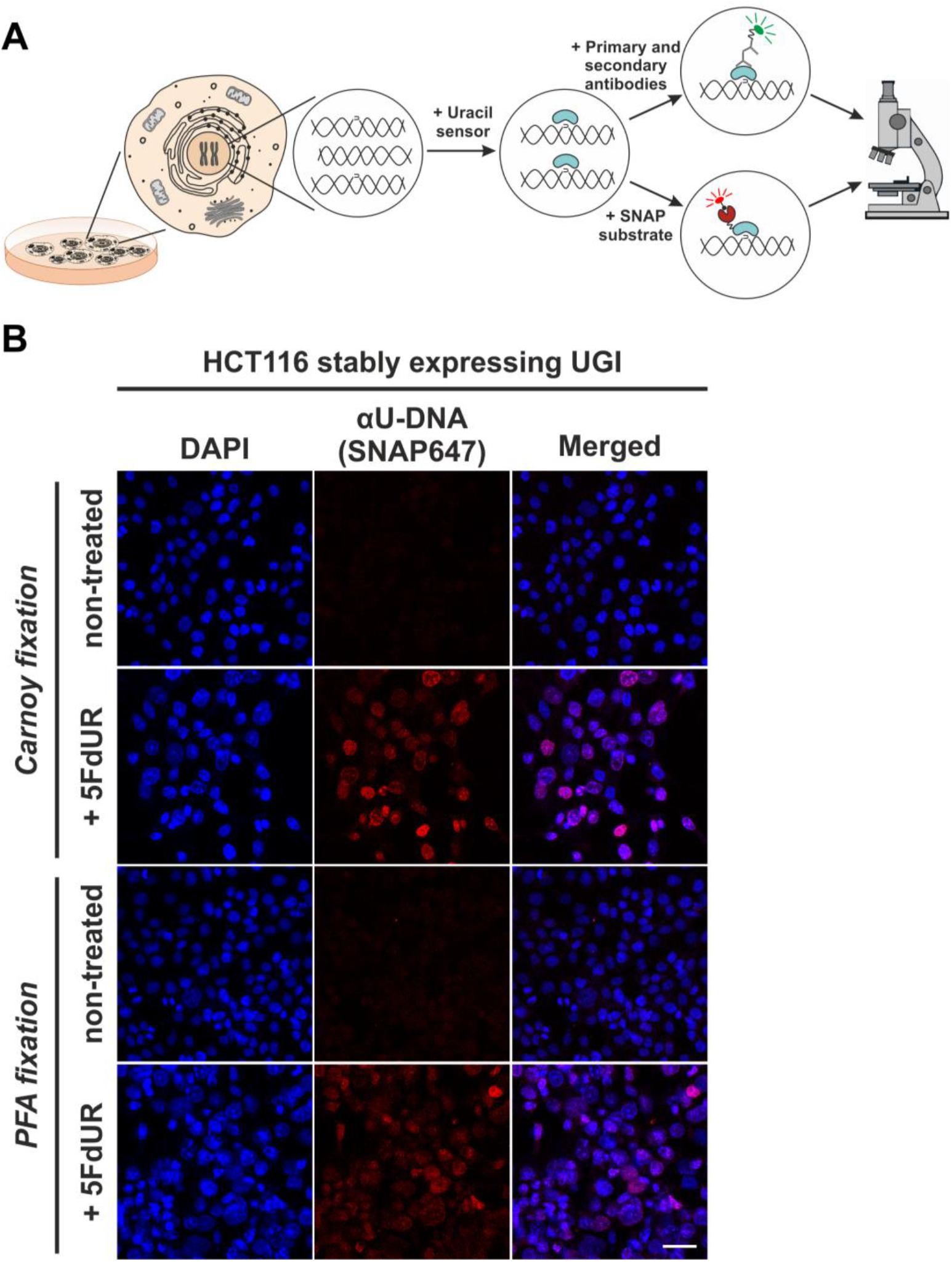
*In situ* detection of the cellular endogenous U-DNA content. **(A)** Scheme represents that genomic uracil residues can be visualized *in situ* using our further developed U-DNA sensor construct via immunocytochemistry (through FLAG-tag) or directly via SNAP-tag chemistry. **(B)** HCT116 cells expressing UGI and treated with 5FdUR show efficient staining with the uracil sensor compared to non-treated cells. Uracil residues are labelled by our FLAG-ΔUNG-SNAP sensor protein visualized by the SNAP647 substrate. DAPI was used for DNA counterstaining. Our optimized staining method is capable of comparable, specific uracil detection in HCT116 cells even with PFA fixation compared to the Carnoy fixation applied previously (Róna et al., 2016). Scale bar represents 40 μm. Note that the nuclei of the treated cells (5FdUR_UGI) are enlarged as compared to the non-treated ones (NT_UGI) presumably due to cell cycle arrest (Huehls et al., 2016; Yan et al., 2016).

### Confocal and STED imaging and analysis

Confocal images were acquired on a Zeiss LSCM 710 microscope using a 20x (NA = 0.8) or a 63x (NA = 1.4) Plan Apo objective or a Leica TCS SP8 STED 3X microscope using a 100x (NA = 1.4) Plan Apo objective. STED images were acquired on the Leica TCS SP8 STED 3X microscope using 660 nm STED (1.5 W, continuous wave) laser for depletion (in combination with Alexa 546). The same image acquisition settings were applied on each sample for comparison. A moderate degree of deconvolution was applied to the recorded STED images using the Huygens STED Deconvolution Wizard (Huygens Software), based on theoretical point spread function (PSF) values. Fluorescence images were processed using ZEN and ImageJ (Fiji) software. 3D projection movies (Supplementary Movies) were constructed from Z-stack images captured by confocal or STED imaging.

### dSTORM imaging and image reconstruction

Super-resolution images were obtained and reconstructed as previously described (Rona et al., 2018). Briefly, dSTORM images were recorded using an in-house built imaging platform based around an inverted microscope. Two colour imaging was carried out sequentially on samples labelled with SNAP-Surface Alexa Fluor 647 and Alexa Fluor 568. The imaging buffer, consisting of 1 mg/ml glucose oxidase, 0.02 mg/ml catalase, 10% glucose, 100 mM mercaptoethylamine (MEA) in PBS, was mixed and added just before imaging. For display purposes, super-resolution images shown in the manuscript have been adjusted for brightness and smoothed; however, quantitative analysis were performed on images before being manually processed to avoid any user bias.

### Interaction factor

The interaction factor (IF) quantifies the colocalization of red and green foci within a cell nucleus by measuring the area of overlap between the two sets of foci (Bermudez-Hernandez et al., 2017; Whelan et al., 2018). The positions of the green foci are then randomized and the overlap between the two colours is measured again. This randomization is repeated 20 times and the interaction factor is the ratio between the experimental overlap area and the mean of the randomized overlap areas. If the red and green foci were completely independent of each other, the IF value would equal one. A value greater than one signifies a higher degree of colocalization compared to a random sample. Non-parametric Mann-Whitney U test was used to calculate statistics on the graphs. Differences of the IF values were considered statistically significant at p < 0.0001 as indicated in Figure 7C-D. Data are presented from two independent biological experiments.

## RESULTS

### Genome-wide mapping of uracil-DNA distribution patterns by U-DNA-Seq

We designed an adequate DNA immunoprecipitation method that can provide U-DNA specific genomic information by next-generation sequencing. This method, termed U-DNA-Seq is based on the rationale of the well-established DIP-Seq technology. Figure 1A presents the scheme of the protocol leading to an enriched U-DNA sample that was then subjected to NGS. Immunoprecipitation was carried out by applying the FLAG-tagged catalytically inactive ΔUNG sensor (described in (Róna et al., 2016)) to bind to uracil in purified and fragmented genomic DNA, followed by a pull-down with anti-FLAG agarose beads.

To allow better detection of nascent uracil, the UNG-inhibitor UGI (derived from *Bacillus subtilis* bacteriophage PBS2) was expressed in HCT116 cells to prevent the action of the major uracil-DNA glycosylase. Besides transient transfection, a stably UGI transfected HCT116 cell line was also established by retroviral transduction of human codon optimized UGI along with EGFP (Supplementary Figure S1A). We proceeded to treat the UGI expressing cells with either 5FdUR or RTX. Notably, this combination of UGI expression and drug treatment did not result in any observable cell death. As shown in Supplementary Figure S1B-C, UGI expression and drug (5FdUR or RTX) treatment led to significantly increased uracil content in genomic DNA. It is important to note that either UGI expression or treatments with drugs targeting *de novo* thymidylate biosynthesis pathways on their own do not lead to elevated U-DNA level (Luo, Walla, & Wyatt, 2008; Róna et al., 2016; Yan et al., 2016). Following U-DNA immunoprecipitation, successful enrichment of U-DNA could be confirmed by dot blot assay in the case of drug-treated cells (5FdUR_UGI or RTX_UGI, Figure 1B). Specificity of U-DNA immunoprecipitation is also underlined by the fact that pull down with empty anti-FLAG beads not containing the U-DNA sensor resulted in negligible amount of DNA (less than 5%). Then, enriched and input DNA samples both from treated (5FdUR_UGI and RTX_UGI) and non-treated (wild type (WT) and NT_UGI) samples were subjected to library preparation and NGS.

Sequencing data were analysed using the herein developed computational pipeline shown in Figure 2 (for more details see the Supplementary Figure S2 and Supplementary Table S1). When reads were aligned to the reference GRCh38 human genome, only uniquely mapped reads were kept and regions suffering from alignment artefacts were excluded from the analysis by blacklisting (Supplementary Figure S3). Statistics on pre-processing steps are shown in Supplementary Table S2. Correlation among the samples and replicates at the level of cleaned aligned reads (bam files) was checked by Pearson correlation analysis (for details see Supplementary Figure S4). Here, a clear difference was shown between the input and the enriched samples; input samples were more similar to each other regardless the applied treatment, while the drug-treated and non-treated enriched samples showed dramatic differences.

There are two principal approaches to extract the signals of uracil enrichment from the cleaned aligned reads: 1) computing genome scaled coverage and log2 ratio tracks, and 2) peak calling that is conventionally used for ChIP-seq data analysis. Log2 ratio tracks provide more detailed information on the uracil-DNA distribution patterns, however, it is not compatible with efficient screening on large dataset (Figure 2 and Supplementary Figure S2). Hence, we generated interval (bed) files from these log2 ratio tracks for each sample that contain simplified information on uracil enriched regions as described in the Supplementary Material. Then, we evaluated both the regions derived from the log2 ratio tracks, and the peak calling results (Supplementary Figure S5 and Supplementary Table S3). We found that the uracil enriched genomic regions are rather broad and much less intense than conventional peaks in ChIP-seq for transcription factors or even for histone modifications. This is somehow expected considering basically stochastic nature of uracil occurrence via both misincorporation and spontaneous cytosine deamination. In agreement with this, reliability and reproducibility of the peak calling approach (using MACS2 with “broad” option) was found to be clearly suboptimal for determination of uracil distribution patterns (Supplementary Figure S5 and Supplementary Table S3). Therefore, we decided to proceed with the coverage track approach rather than the peak calling. All of the main figures rely on analysis performed with either the log2 ratio tracks or the regions of uracil enrichment derived from the log2 ratio tracks.

The log2 ratio tracks were generated by comparing the genome scaled sequencing coverages of the enriched U-DNA and the input sample (for details see Supplementary Material). Figure 3A shows the uracil distribution pattern in a selected chromosomal segment where an uneven distribution with variably spaced broad regions is observed (the same data for all the chromosomes are shown in Supplementary Figure S6). A clear difference between non-treated (WT and NT_UGI) and treated (5FdUR_UGI and RTX_UGI) cells is already obvious from this view, and the correlations were also measured quantitatively on the whole log2 ratio tracks by Pearson correlation coefficients and related scatter plots (Figure 3B, for individual replicates see Supplementary Figure S7).

The uracil-enrichment coverage tracks in Figure 3A and the related correlations in Figure 3B already suggested altered distribution of uracil-containing regions in the drug-treated (5FdUR_UGI and RTX_UGI) as compared to the non-treated (WT and NT_UGI) samples. This difference was further underlined in a histogram representation of uracil enrichment signal (Figure 3C) where drug treatment led to a higher number of genomic segments (more data bins) with increased uracil level. It was of immediate interest to investigate whether the uracil distribution patterns, distinctly characteristic for the non-treated versus drug-treated samples might show any correlation to any previously determined genomic features. For this reason, we built a relevant database by collecting cell type specific ChIP-seq and DNA accessibility data (for details see Supplementary Material), since epigenetic modifications and regulation occur in diverse fashion in different cell types.

Interrogation of the constructed specialized database with respect to the uracil-DNA distribution patterns was performed using interval (bed) files of uracil enriched regions (derived from log2 ratio track) for each U-DNA-Seq sample. To screen for similarity between sample and database interval (bed) files, we applied the GIGGLE search tool (full data are presented in Supplementary Appendix 2). GIGGLE scores are capable to adequately represent the measure of colocalization independently from the size of the compared intervals (Layer et al., 2018). Each interval file in the database corresponded to one specific ChIP-seq experiment with a given factor (e.g. histone markers, transcription factors, etc.). GIGGLE scores were then calculated pairwise (each sample to each database interval files), and plotted for the top ten factors corresponding to the highest scores (Figure 4A). The similarity scores of the U-DNA-Seq data with regard to the different chromatin markers indicate that non-treated cells (WT and NT_UGI) may possess uracils preferentially in the constitutive heterochromatin (high scores with H3K9me2 and H3K9me3 (Hyun, Jeon, Park, & Kim, 2017; Saksouk, Simboeck, & Déjardin, 2015)). On the other hand, drug treatment of the cells either with 5FdUR (5FdUR_UGI) or RTX (RTX_UGI), induces uracil incorporation into more active genomic segments, correlated with high similarity scores to euchromatin histone marks (H3K36me3 (Becker et al., 2017; Hyun et al., 2017; Pfister et al., 2014), H3K4me1/3 (Hyun et al., 2017), H3K27ac (Creyghton et al., 2010), H3K9ac (Gates et al., 2017)), or factors associated to either activation or repression in a context dependent manner (SP1 (Doetzlhofer et al., 1999), H3K27me3 (Becker et al., 2017; Saksouk et al., 2015), H2AZ/AFZ (Giaimo, Ferrante, Herchenröther, Hake, & Borggrefe, 2019)) (Figure 4A).

Based on the detected correlation with hetero- and euchromatin in case of non-treated and drug-treated cells, respectively, we wished to determine whether it might be reflected in other, more generalized genomic features also. Therefore, we investigated colocalization of U-DNA enriched regions with several genomic features using bedtools annotate (Quinlan & Hall, 2010) to extract the number of overlapping bases. Scores measuring the colocalization are presented in Figure 4B for a systematic selection of the tested features. The results of the full analysis are provided in Supplementary Table S4. The data suggest that uracil incorporation in transcriptionally active (e.g. active promoters, DNase hypersensitive sites), and potentially active genomic segments (CpG islands, genes, especially exons and CDS regions) is increased upon drug treatment. The proposed uracil enrichment in transcriptionally active genomic regions is also in agreement with the colocalization with different repeat classes: the drug-treated samples show higher colocalization with short interspersed elements (SINE (Kramerov & Vassetzky, 2005)) and long terminal repeats (LTR (Kovalskaya, Buzdin, Gogvadze, Vinogradova, & Sverdlov, 2006)) which are known to be more frequently transcribed as compared to long interspersed element (LINE (Boissinot & Furano, 2005)) and Satellite segments (López-Flores & Garrido-Ramos, 2012).

The observed similarity between wild type uracil distribution and the patterns of histone markers associated with heterochromatin (Figure 4A) is further underlined by the positive correlation between U-DNA and cytogenic chromosome G-bands (Figure 4B). Dark G-bands stained strongly by Giemsa was shown to correlate with AT-rich, heterochromatic, late replicating genomic segments (Gilbert, 2002; Holmquist, Gray, Porter, & Jordan, 1982). In contrast, negative G-bands are correlated better to the drug-treated uracil-DNA distribution pattern, also in agreement with our results from the comparison to histone markers. Consistently, similar difference between patterns of U-DNA in non-treated versus drug-treated cells in early or late replicating genomic segments is also revealed. Late replicating regions are better correlated to the U-DNA distribution in non-treated cells, while the drug treatment induced U-DNA pattern is more similar to the early replicating segments (Figure 4B). It is widely accepted that replication timing strongly correlates with chromatin structure, namely the open euchromatin and the condensed heterochromatin replicates in early and late S-phase, respectively (Gilbert, 2002). The correlation between U-DNA enrichment and replication timing was further analysed using a better resolved time scale of replication (Figure 4C) which strengthened the initial observation. The correlations with G-banding and replication timing are also clearly visible on IGV views in Supplementary Figure S6. Furthermore, colocalization with AT-rich heterochromatin for non-treated and GC-rich euchromatin for drug-treated samples is also reflected by the base composition of uracil enriched regions (Supplementary Table S3A). The surprisingly high correlation between uracil enrichment in drug-treated cells and CpG islands (cf. Figure 4B) coincides with the elevated GC content of uracil enriched genomic regions in these samples.

### *In situ* detection of U-DNA using super-resolution microscopy

We aimed to correlate genome-wide uracil distribution patterns with *in situ* localization in the context of chromatin architecture. Therefore, we further develop the U-DNA sensor constructs (Róna et al., 2016) to allow *in situ* detection of genomic U-DNA in complex eukaryotic cells using microscopy. Figure 5A shows a schematic representation of the U-DNA staining procedure. The U-DNA sensor constructs were fused to different tags allowing antibody-based or direct detection via fluorescence microscopy. In order to achieve a versatile labelling technique to facilitate super-resolution imaging of U-DNA, we attached SNAP-tag to the C-terminal end of ΔUNG yielding FLAG-ΔUNG-SNAP, generating a novel sensor construct (Supplementary Figure S8A). The SNAP-tag offers a flexible biorthogonal chemical labelling strategy as it reacts specifically and covalently with benzylguanine derivatives, permitting the irreversible labelling of SNAP fusion proteins with a wide variety of synthetic probes (Keppler et al., 2003). In order to check whether the functionality of this new construct is still preserved, we performed dot blot and staining experiments. Results shown in Supplementary Figure S8B indicate that the FLAG-ΔUNG-SNAP construct is functional and shows similarly reliable U-DNA detection using dot blot approach, when compared to FLAG-ΔUNG-DsRed protein described previously (Róna et al., 2016). Supplementary Figure S8C shows that the new labelling construct, FLAG-ΔUNG-SNAP, also recognizes the presence of extrachromosomal uracil enriched plasmid aggregates in the cytoplasm. These results confirmed that the FLAG-ΔUNG-SNAP construct is capable of U-DNA detection in dot blot assays and suitable for *in situ* staining applications.

Our goal was to use this new sensor to detect *in situ* endogenous uracils in human cells in a setup that also allows colocalization with other chromatin factors. For visualization of our sensor, photostable SNAP-tag substrates (here SNAP647 or SNAP546) were used. Figure 5B shows that drug treatment and the inhibition of cellular UNG enzyme by UGI leads to significantly increased uracil content in genomic DNA that is readily observable on conventional confocal microscopic images. Figure 5B also demonstrates that our FLAG-ΔUNG-SNAP sensor can be applied for straightforward staining of genomic uracil after either Carnoy (as used previously (Róna et al., 2016)) or PFA fixation. Unlike Carnoy, PFA fixative is compatible with most antibody-based staining procedures, thus it is suitable for multi-colour imaging allowing colocalization studies. Next, we attempted to use super-resolution microscopy to have a better track of the uracil distribution pattern even in case of the low genomic uracil level found in the non-treated cells. Figure 6 compares confocal, STED and dSTORM microscopy techniques for U-DNA detection. The exquisite sensitivity of dSTORM is apparent from these experiments as it can detect the low level of genomic uracil in non-treated cells (cf. Figure 6B). Importantly, we observed different heterogeneous staining in the nucleus for uracil in non-treated and drug-treated cells. Furthermore, images of drug-treated cells show uracil staining with signal enrichment at the nuclear membrane and areas surrounding the nucleoli. Supplementary Movies SM1-SM4 (also Supplementary Figure S9) contribute to further visualization of uracil distribution captured by confocal and STED imaging.

**Figure 6.**
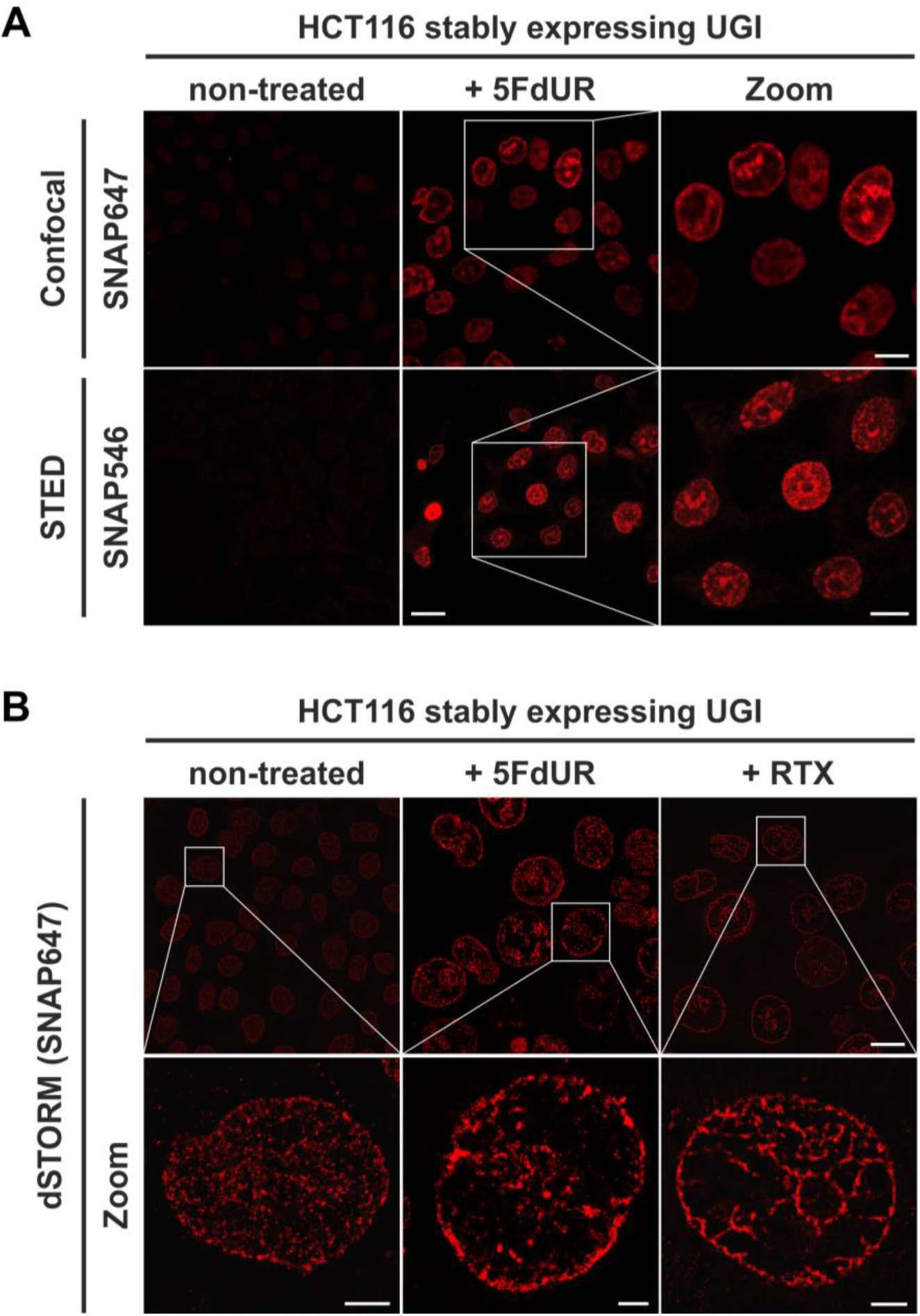
The FLAG-ΔUNG-SNAP sensor enables super-resolution detection of genomic uracil by STED and dSTORM microscopy. **(A)** U-DNA staining was performed on non-treated or 5FdUR treated HCT116 cells stably expressing UGI. Different SNAP-tag substrates, SNAP647 for confocal and SNAP546 for super-resolution imaging (STED) were used to label FLAG-ΔUNG-SNAP. Scale bar represents 20 μm for whole images and 10 μm for zoomed sections. **(B)** dSTORM imaging was also performed on non-treated or drug-treated (5FdUR or RTX) HCT116 cells stably expressing UGI to compare the sensitivity of these imaging techniques. U-DNA staining shows a characteristic distribution pattern in cells with elevated uracil levels as compared to non-treated cells. SNAP647 substrate was used to label FLAG-ΔUNG-SNAP. Scale bar represents 10 μm for whole images and 2 μm for zoomed sections.

Based on the genome-wide sequencing data analysis, we proceeded to select cognate chromatin markers for colocalization studies. As shown in Figure 4A, the highest similarity (GIGGLE) score corresponded to H3K36me3 and H3K27me3 for the RTX and 5FdUR treated samples, respectively. Also, both of these chromatin markers showed positive correlation with each drug-treated, while negative correlation with each non-treated sample. Using the herein demonstrated immunofluorescence protocol we obtained co-stained images of uracil and these histone markers by both confocal and dSTORM microscopies (Figure 7A-B). Validating U-DNA-Seq data, we found that U-DNA staining shows significant colocalization with staining for both chromatin markers; H3K36me3 and H3K27me3, based on cross-pair correlation analysis of the super-resolution images as shown in Figure 7C-D. The rate of colocalization, as determined by the interaction factor (IF) value (Bermudez-Hernandez et al., 2017; Whelan et al., 2018), was statistically significant between the uracil signal and both chromatin markers in each case of drug treatment, when compared to the non-treated sample as well as to a generated set of random distribution patterns of these chromatin markers.

**Figure 7.**
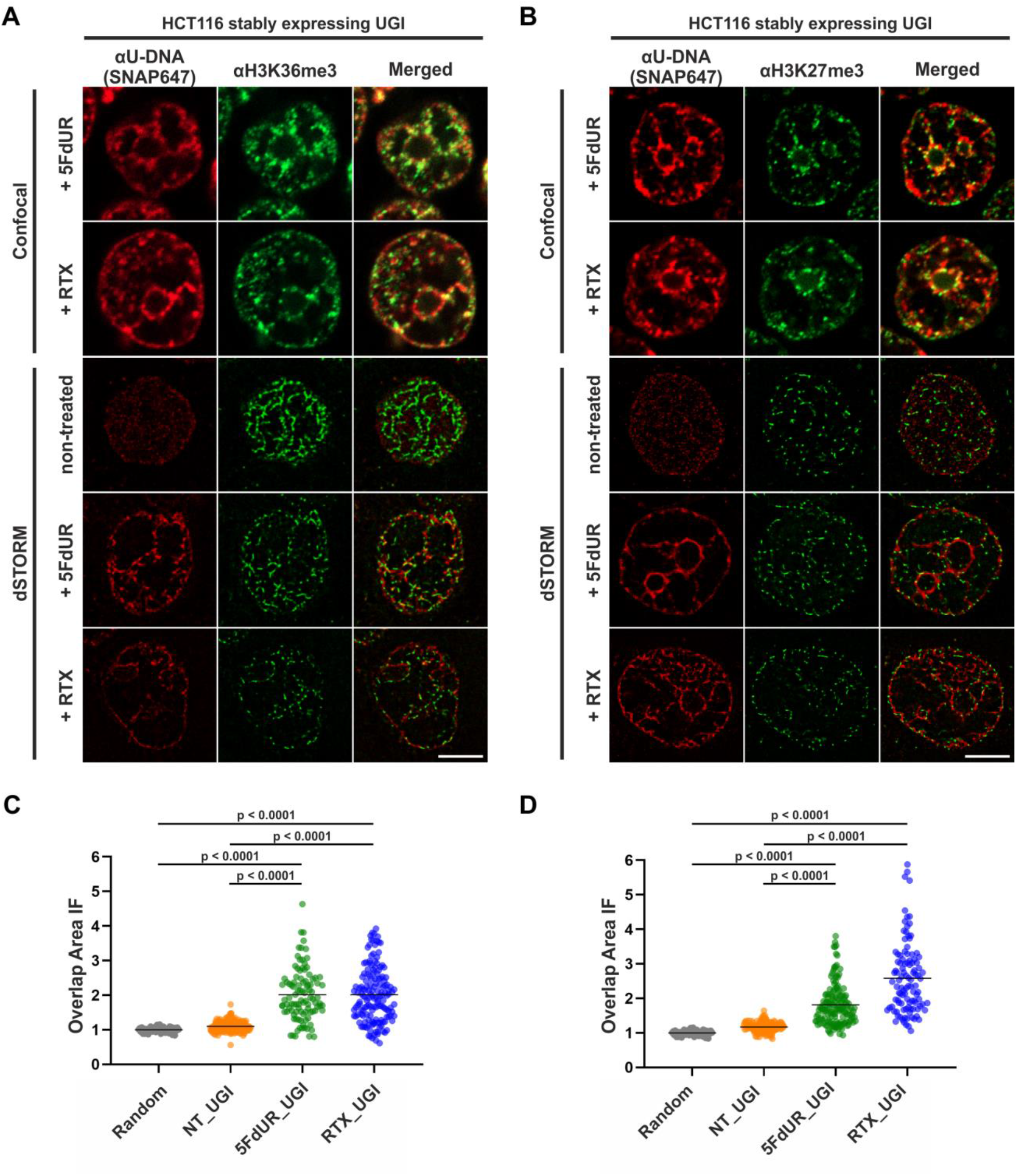
Genomic uracil moieties colocalize with H3K36me3 and H3K27me3 analysed by super-resolution microscopy. Confocal and dSTORM imaging were performed on non-treated, 5FdUR or RTX treated HCT116 cells stably expressing UGI to compare the localization of genomic uracil residues (red) to histone markers, H3K36me3 (green) **(A)** or H3K27me3 (green) **(B)**, selected based on the U-DNA-Seq results. Scale bar represents 5 μm. The graphs display the Cross-Pair-Correlation analysis between U-DNA and H3K36me3 **(C)** or H3K27me3 **(D)**, respectively. Overlap is defined as any amount of pixel overlap between segmented objects. Total number of analysed nuclei for H3K36me3 staining **(C)** were the following: NT_UGI (n=205), 5FdUR_UGI (n=101) and RTX_UGI (n=153) from 2 independent experiments. Total number of analysed nuclei for H3K27me3 staining **(D)** were the following: NT_UGI (n=154), 5FdUR_UGI (n=151) and RTX_UGI (n=107) from 2 independent experiments. Black line denotes the mean of each dataset. The colour code follows the one in Figure 3A.

## DISCUSSION

We present here the U-DNA-Seq method that can provide genome-wide uracil distribution data. Such information is highly beneficial as the global U-DNA quantification method published thus far (Galashevskaya et al., 2013) cannot address the genome-wide localization of uracils. U-DNA-Seq is a direct, feasible alternative to the recently published UPD-seq (Sakhtemani et al., 2019), Excision-seq (Bryan et al., 2014) or dU-seq (Shu et al., 2018) methods, all of which allows only indirect detection requiring one or more auxiliary chemical or enzymatic step(s). Only these three methods have the potential thus far to map genome-wide distribution of uracil within isolated genomic DNA based on NGS, and only dU-seq was used in the context of human genome. Each of these three methods has advantages and also limitations. UPD-Seq detects abasic (AP) sites that can be generated from numerous base alterations through the dedicated DNA glycosylase enzymes. dU-seq follows a complex workflow on isolated genomic DNA that results in replacement of deoxyuridines with biotinylated nucleotides that is pulled down using streptavidin beads and is subjected to sequencing (Shu et al., 2018). While dU-seq and also Excision-seq rely on multiple enzymatic reactions initiated by UNG, U-DNA-Seq is a direct and less labour-intensive method employing U-DNA specific binding of catalytically inactive UNG-derived sensor constructs. Both dU-seq and U-DNA-Seq involve enrichment of uracil containing DNA fragments that increase sensitivity, while Excision-seq does not apply enrichment and relies on differential ligation of fragments excised through base excision repair. Excision-seq and UPD-seq were reported as adequate methods for efficient detection of elevated uracil levels in smaller genome sizes as in mutant *Escherichia coli* (*E. coli*) and yeast strains. The Excision-seq experiment suggested correlation between uracil accumulation and replication timing (Bryan et al., 2014). Nevertheless, the large mammalian genome size and the low frequency and/or nature of the distribution of uracils might result in some biases or underestimation using Excision-seq method.

Notably, dU-seq and Excision-seq are potentially capable to identify the exact position of individual uracils, providing single-base resolution. Considering the basically stochastic nature of uracil incorporation during DNA synthesis due to insensitivity of the polymerases, and spontaneous cytosine deamination, this aspect has lower impact. Due to the stochastic processes, the actual positions of uracil are expected to be variable in every single cell. Therefore, a statistical approach has higher descriptive value about the uracil distribution.

In addition, the usual analysis methods designed for ChIP-seq experiment were proved to be suboptimal. We therefore constructed a novel computational pipeline that allows reliable data analysis avoiding overinterpretation. Re-analysis of the earlier published dU-seq data with the herein developed pipeline (cf. Supplementary Table S5), showed very high correlation with our U-DNA-Seq data in case of comparable samples (non-treated K562 cells in both cases; and 5FdUR-treated UGI expressing HCT116 vs 5FdUR treated UNG^−/−^ HEK293T cells, cf. Supplementary Figure S10) confirming robustness and reliability of our method. However, our interpretation is markedly different regarding the preferential centromeric location of uracils that has been suggested by *Shu et al* (Shu et al., 2018). We argue that based on short-read sequencing data, centromeres cannot be assessed, thereby, the proposed centromeric uracil enrichment cannot be confirmed by dU-seq (see detailed argumentation in Supplementary Material, Supplementary Figure S11).

Using U-DNA-Seq, here we demonstrate that distribution of uracil-containing regions is altered in the drug-treated (5FdUR or RTX, in combination with UGI) as compared to the non-treated (wild type and UGI expressing) samples. The genomic uracil distribution patterns either in non-treated and in drug-treated cells are found to be non-random: broad regions of uracil enriched genomic segments were detected. Within the third part of our pipeline (cf. Figure 2, and Supplementary Figure 2), we also analysed the distribution pattern of these broad peaks comparing them to a set of relevant and cell type specific data of ChIP-seq experiments and other genomic features. In drug-treated cells, these broad segments showed highest correlation with ChIP-seq-based patterns published for predominantly euchromatin and facultative heterochromatin markers (Figure 4). Increasing evidence suggests that active and repressed chromatin states can be determined in a combinatorial fashion where simultaneous histone marks can efficiently shift gene expression from inactive to active states or vice versa (Gates et al., 2017; Hyun et al., 2017). Hence, it is of special interest to note that our colocalization data show similarity scores not just for one but for a variety of factors. Importantly, regarding these factors and additional features, our results are highly coherent. Namely, the outstanding correlation of uracil-DNA patterns in drug-treated samples with active promoters, CpG islands, early replicating segments and DNase hypersensitive sites, highly supports the above conclusion. Euchromatin was shown to imply early replicating genomic regions, whereas heterochromatin replicates in late S-phase (Black, Van Rechem, & Whetstine, 2012). Accordingly, we report that the drug treatment induced U-DNA pattern is more similar to the early replicating segments, whereas U-DNA distribution in non-treated (wild type and UGI-expressing) cells shows simultaneous association with both heterochromatin markers and late replicating regions (Figure 4B-C).

Consistently with our observations, *Weeks et al* very recently showed that treatment with the antifolate pemetrexed in UNG −/− human colon cancer cells led to preferential enrichment of double-strand breaks (DSBs) within highly accessible euchromatic regions, like transcription factor binding sites, origins of replication, DNase hypersensitivity regions and CpG islands (Weeks, Zentner, Scacheri, & Gerson, 2014). This study did not directly address the occurrence of uracil moieties but caught the process initiated by uracil incorporation at a later stage. Still, the distribution pattern of the resulting DSBs showed similarities to our U-DNA-Seq data.

Taken together, in the non-treated cells, where the level of genomic uracil is low, we show that it is preferentially located in the constitutive heterochromatin, which can be explained by the fact that heterochromatin is generally highly condensed and thus less accessible for DNA repair and replicative DNA synthesis. In contrast, in the open, more frequently transcribed euchromatin, DNA repair can efficiently correct uracils in the presence of a balanced dNTP pool. The low amount of genomic uracil in non-treated cells might remain from either cytosine deamination or thymine replacing misincorporation that escaped DNA repair. However, drug (5FdUR or RTX) treatment perturbs the cellular nucleotide pool, and consequently highly increase the rate of thymine replacing uracil misincorporation events overwriting the background uracil pattern of non-treated cells’ genome. Uracil appearance via thymine replacing misincorporation implies prior DNA synthesis involved in either replication, transcription-coupled DNA repair or epigenetic reprograming (e. g. erasing the methyl-cytosine epigenetic mark). Importantly, we found that uracil pattern showed the highest correlation with the features (early replication, active promoters and DNase hypersensitive sites, and CpG islands) linked exactly to these processes (cf. Figure 4B).

The antifolate or nucleotide-based thymidylate synthase inhibitors, such as 5-FU, RTX or 5FdUR are known to lead to cell-cycle arrest, as it is reflected in the detected uracil-DNA pattern that strongly correlates with the early replicating segments in case of both drug treatments. The two drugs caused similar, but not equivalent uracil-DNA pattern. On the one hand, the correlation with the H3K36me3 marker as well as with the early replicating segments are both markedly stronger with the RTX treated sample as compared to the 5FdUR treated sample (cf. Figure 4). On the other hand, the correlation of uracil accumulation with the H3K27me3 marker, and with the CpG islands is stronger in the 5FdUR treated sample. Such differences might correspond to drug-specific mechanism of action, involving alterations in signalling processes, transcription regulation and the timing of cell-cycle arrest (Van Triest, Pinedo, Giaccone, & Peters, 2000). Details of these mechanisms remain obscure in the literature. Still, it is well-known that both drugs inhibit thymidylate synthase thereby facilitating dUTP incorporation into DNA, while the nucleotide analogue 5FdUR also leads to direct incorporation of 5-fluorodeoxyuridine triphosphate (FdUTP) into the DNA (Longley, Harkin, & Johnston, 2003; Pettersen et al., 2011). Genomic uracil and fluorouracil (FU) might have different effects on transcription and epigenetic regulation processes that might also contribute to the observed differences of the two U-DNA patterns. It should be noted that our method detects both uracil and also FU within the DNA, since the UNG enzyme also binds to FU (Pettersen et al., 2011). Phenotypic differences in cell-cycle progression upon the two drug treatments were also reported. The 5FdUR treatment was shown to cause an S-phase arrest in the second cycle (Huehls et al., 2016; Yan et al., 2016), while the actual time point of cell-cycle arrest upon RTX treatment is still controversial (Blackledge, 1998; Ding et al., 2019; Zhao, Zhang, Sun, Zhan, & Zhao, 2016).

The new U-DNA-Seq method was shown to be reliable, robust and potent enough to gain systematic information on uracil-DNA metabolism upon drug treatments. Such information could essentially contribute to the future understanding of the mechanistic details either of cytotoxic effect induced by anti-cancer drugs, or other biological processes involving genomic uracil appearance. To this end, it is also of key importance to establish new visualization methods allowing colocalization measurements between U-DNA and other factors in highly complex eukaryotic cells.

Therefore, we further developed the U-DNA sensor to visualize genomic uracil *in situ* in human cells. The FLAG-ΔUNG-SNAP sensor construct and the optimized staining method presented here were successfully applied in confocal and super-resolution (STED or dSTORM) microscopies (see Figures 5–7). To our knowledge, there is no alternative technique published so far for *in situ* microscopic detection of mammalian genomic uracil. A recent paper was published reporting a similar approach, where uracil-DNA glycosylase UdgX was coupled to a fluorescent tag and applied for staining of uracils in *E. coli* DNA (Datta et al., 2019), however, in our previous study ΔUNG had already been proved to be potent for *in situ* uracil detection in the same organism (Róna et al., 2016). Still, the UdgX-based tool was not further extended for detection of uracils within the highly complex chromatin of human cells. Moreover, our detection method also allows simultaneous staining for other factors in colocalization experiments, potentially providing mechanistic insight of several important biological phenomena that involves uracil-DNA. In the present study, two histone markers were selected based on the U-DNA-Seq results for colocalization studies. Using dSTORM super-resolution microscopy we could confirm the statistically significant correlation of genomic uracil with the two selected histone markers (H3K36me3 and H3K27me3) in drug-treated (5FdUR or RTX), UGI-expressing cells (Figure 7). The H3K36me3 was shown to associate with actively transcribed genes (Becker et al., 2017; Hyun et al., 2017; Pfister et al., 2014), while H3K27me3 is the most cited marker for facultative heterochromatin (Becker et al., 2017; Saksouk et al., 2015). In summary, co-staining of genomic uracil in drug-treated cells and the selected histone markers via dSTORM reinforced the association between uracil occurrence and transcriptionally active regions.

It has been argued that uracil accumulation may play a more decisive role in genomic instability than the induced uracil-excision repair (Huehls et al., 2016; Yan et al., 2016). Uracil in DNA may therefore be used as a key marker for efficiency of chemotherapeutic drugs targeting thymidylate biosynthesis. Our presently developed techniques to follow the extent and pattern of uracilation induced by several chemotherapeutic drugs may provide key novel insights into the mechanism of drug action.

## DATA AVAILABILITY

Sequencing data have been deposited into the Gene Expression Omnibus (GEO) under accession number GSE126822.

## SUPPLEMENTARY DATA

Supplementary Data are also available.

## ACKNOWLEDGEMENTS

We gratefully acknowledged the kind help of György Török and László Homolya in acquiring fluorescent images via STED microscopy. We also wish to say sincere thanks to György Várady in FACS sorting experiments, and to Gábor Tusnády for providing access to computational capacity. We acknowledge the ENCODE Consortium (ENCODE Project Consortium, 2012) and the ENCODE production laboratory(s) generating the particular dataset(s) as well as the contributors of the UCSC Table Browser (Karolchik et al., 2004; Kuhn, Haussler, & Kent, 2013) data. We also acknowledge the contributors of Ensembl (Zerbino et al., 2018), ReplicationDomain (Weddington et al., 2008), Cistrome Data Browser (Mei et al., 2017) for making their data publicly available.

