## Supplementary Material for "Genome-wide alterations of uracil distribution patterns in human DNA upon chemotherapeutic treatments"

**Table of Content**

1. Effects of UNG inhibition and drug treatment on genomic uracil content ... 4

Supplementary Figure S1. Elevation of genomic uracil content upon stable UGI expression and drug treatments. ... 4

2. Detailed analysis pipeline – methods of U-DNA-Seq data analysis ... 6

Supplementary Figure S2. Detailed analysis pipeline. ... 7

Supplementary Table S1. Details on the applied tools. ... 8

2.1. Preprocessing ... 9

Supplementary Figure S3. Creation of cell line specific blacklists. ... 13

Supplementary Table S2. Number of reads in samples during the preprocessing steps. ... 14

Supplementary Figure S4. Pearson correlation among processed BAM files. ... 16

2.2. Determination of uracil enrichment: log2 ratio track and derived regions versus peaks called by MACS2 tool. ... 16

Supplementary Figure S5. Representative IGV views from aligned reads to a 10 Mb cluster of uracil enriched regions compared to usual peak calling results. ... 19

Supplementary Table S3. Comparison of regions of uracil enrichment derived from log2 signal ratio and peaks called by MACS2. ... 21

Supplementary Figure S6. IGV view of log2 ratio and regions of uracil enrichment on chromosome 1 (for all chromosomes see Appendix 1). ... 23

Supplementary Figure S7. Pearson correlation among log2 ratio tracks of replicates. ... 24

2.3. Genome-wide analysis of uracil-DNA pattern comparing to ChIP-seq data and other genomic features ... 24

Supplementary Table S4. Full collection of genomic features compared to the four samples of U-DNA-Seq. ... 29

3. Characterization of the new uracil-sensor construct ... 32

Supplementary Figure S8. Design and validation of the new FLAG-ΔUNG-SNAP uracil-sensor. ... 32

4. *In situ* detection of the cellular U-DNA content by STED microscopy ... 33

Supplementary Figure S9. Representative image for Supplementary Movies SM1-SM4. ... 33

Supplementary Movie SM1. Confocal Z-stack series of 3 cells showing uracil distribution. ... 33

Supplementary Movie SM2. 3D projection of Supplementary Movie SM1. ... 33

Supplementary Movie SM3. Z-stack series of 1 cell acquired by STED showing uracil distribution. ... 33

Supplementary Movie SM4. 3D projection of Supplementary Movie SM3. ... 33

5. Re-analysis and reinterpretation of dU-seq data published in (15) ... 34

Supplementary Table S5. Statistics on preprocessing of dU-seq data. ... 36

### 1. Effects of UNG inhibition and drug treatment on genomic uracil content

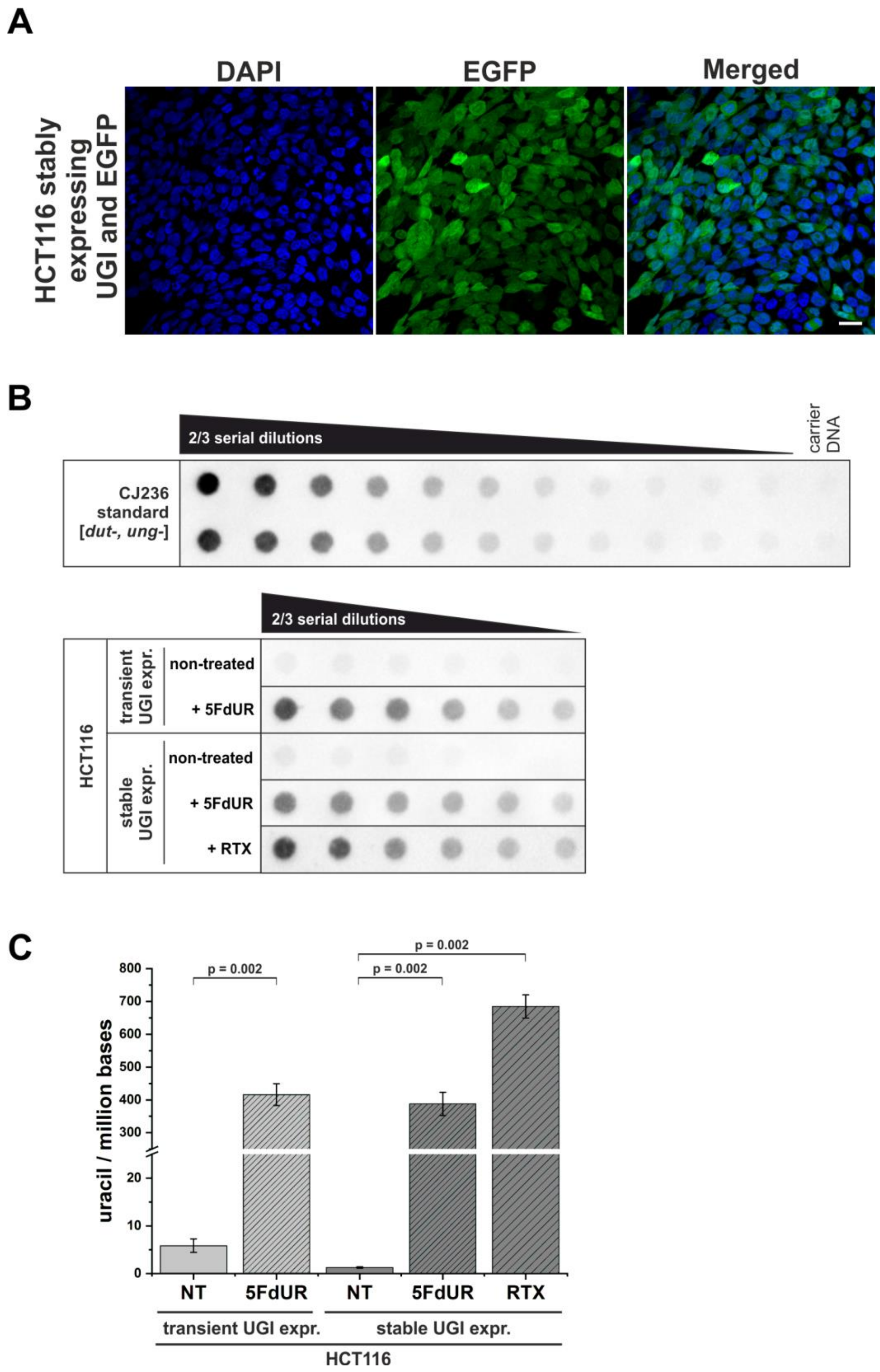

**Supplementary Figure S1. Elevation of genomic uracil content upon stable UGI expression and drug treatments. (A)** HCT116 cells stably expressing UGI along with EGFP following retroviral transduction. GFP positive cells were selected by fluorescence-activated cell sorting and cultured for further analysis. DAPI was used for DNA staining. Scale bar represents 20 µm. **(B)** Dot blot assay for measuring genomic uracil levels (1) of non-treated and drug (5FdUR or RTX) treated HCT116 cells either transiently or stably expressing UGI. Genomic DNA (8 ng) isolated from log-phase growing CJ236 [*dut-, ung-*] *Escherichia coli* strain was applied as uracil standard in a ½ dilution series (upper panel). Two-third dilution series from HCT116 samples started with 600 ng DNA for non-treated or 5 ng DNA for 5FdUR or RTX treated samples (lower panel). The dot blot image presented is a representative of six independent biological experiments. **(C)** Bar graph shows the uracil moieties / million bases of each sample. Drug treatment led to significantly elevated uracil levels in HCT116 cells either transiently or stably expressing UGI (~400 uracil moieties/million for 5FdUR and ~700 uracil moieties/million for RTX) as compared to non-treated (NT) cells (~2-5 uracil moieties/million). Error bars indicate standard errors of the mean. Calculations were based on six independent datasets (n = 6). p = 0.002.

### 2. Detailed analysis pipeline – methods of U-DNA-Seq data analysis

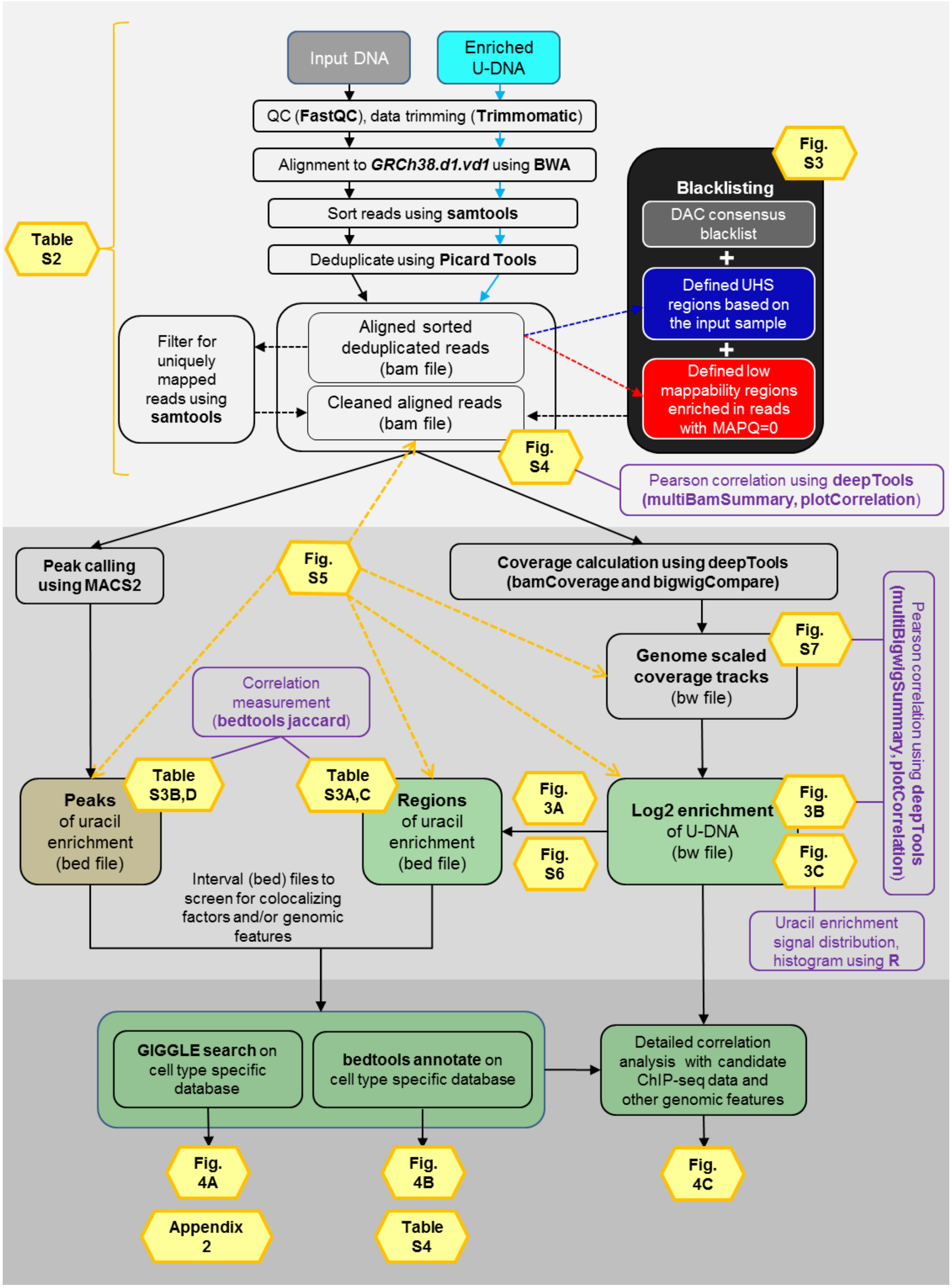

**Supplementary Figure S2. Detailed analysis pipeline.** Input DNA and enriched U-DNA mean genomic DNA fragmented by sonication and uracil containing fragments pulled down by FLAG-ΔUNG sensor, respectively. Preprocessing steps (light grey field) include quality check, data trimming, read alignment, sorting, deduplication, filtering out ambiguously mapped reads (mapping quality, MAPQ=0) and blacklisting based on consensus DAC blacklist recommended by ENCODE Consortium (2), the ultra-high signal (UHS) and low mappability (more than 50% of the reads are with MAPQ=0) regions. Pearson correlation among samples at the level of aligned reads in bam files can be calculated using deepTools package (3). Data processing (darker grey field) can follow two main streams: 1) peak calling originally designed for ChIP-seq (using MACS2 (4, 5), resulting in peaks' interval files in bed format); and 2) coverage track calculation and analysis (using deepTools, resulting in coverage track files in bigwig/bw format). Interval (bed) files can be derived not only from peak calling (peaks.bed) but from log2 coverage signal ratio (enriched/input) bigwig files also (regions.bed). Additional calculations for sample characterization are indicated in purple boxes. In the third part (darkest grey field), uracil enrichment profile was further analysed by comparing to available data. First, a quick screen using interval (bed) files was performed, and then a detailed correlation analysis was carried out with the promising candidates using coverage track (bw) files. For the initial screen, we collected a set of HCT116 specific ChIP-seq and DNA-IP-seq data, as well as other genomic features in the format of interval (bed) files. GIGGLE search (6) and bedtools annotate (7) were used for scoring the similarities between the query uracil-DNA and the database intervals.

| Program package | tool | purpose | version | Link, reference |
|---|---|---|---|---|
| FastQC | | Quality checking | 0.11.7 | https://www.bioinformatics.babraham.ac.uk/projects/fastqc |
| Trimmomatic | | Adapter and quality trimming | 0.36 | https://github.com/timflutre/trimmomatic (8) |
| BWA | MEM | *Burrows-Wheeler Aligner* | 0.7.17 | https://github.com/lh3/bwa (9) |
| samtools | view | Filtering reads in bam files | 1.9 | https://github.com/samtools/samtools (12) |
| | merge | Concatenating bam files | | |
| | sort | Sorting reads in a bam file (required by most of the downstream application) | | |
| | index | Indexing bam files (required by most of the downstream application) | | |
| | idxstats | Reporting the numbers of mapped and unmapped reads in an indexed bam file along the chromosomes and scafolds in the reference genome | | |
| Picard Tools | MarkDuplicates | Filtering out reads corresponding to PCR or optical duplicates | 1.95 | http://broadinstitute.github.io/picard |
| deepTools | multiBamSummary | Genome-wide comparison of multiple bam files regarding the readcoverage in defined sized bins | 3.2.1 | https://github.com/deeptools/deepTools/releases (3) |
| | bamCoverage | Calculating genome scaled readcoverage tracks in databins and with the option of smoothing resulting in bedgaph or bigWig files | | |
| | bigwigCompare | Comparing two bigWig files in many different ways e.g. log2 ratio or subtract | | |
| | multiBigWigSummary | Genome-wide comparison of multiple bw files in defined sized bins | | |
| | plotCorrelation | Calculating and ploting the correlation coefficients from the results of multiBigWigSummary or multiBamSummary | | |
| bedtools2 | merge | Merging intervals in files in many different ways | 2.28.0 | https://github.com/arq5x/bedtools2 (7) |
| | subtract | Subtracting intervals in files in different ways | | |
| | complement | Taking the complement of an interval file comparing to a reference genome | | |
| | intersect | Extracting overlaping fractions of interval files in many different ways | | |
| | jaccard | Calculating Jaccard indices (ratio of base numbers in the intersect over the union of two interval files) | | |
| | annotate | Comparing query interval file to a set of database interval files, and reporting overlap ratio and/or the number of overlaping intervals for each interval in the query bed file | | |
| GIGGLE | sort_bed | A script utilizing also bgzip, to sort and compress bed files for giggle search | 1.0 | https://github.com/ryanlayer/giggle (6) |
| | Index | Special indexing applied for the library of the database interval files | | |
| | search | Scoring collocalization between a query and indexed database interval files | | |
| kentUtils | bigWigToWig | conversion tool from the binary coded bigWig to a text format Wiggle file | | https://github.com/ucscGenomeBrowser/kent |
| | bigWigAverageOverBed | Averaging scores in a bw files for the intervals given in a bed files | | |
| | liftOver | Converting genomic coordinates in a bed file from one to another reference genome version | | |
| MACS2 | callpeak | Calling peaks of readcoverage, standard tool in ChIP-seq data analysis | 2.1.2 | https://github.com/taoliu/MACS (4,5) |
| R | | Environment for statistical computing | 3.5.1 | https://www.R-project.org/ (14) |
| Linux command-line utilities | awk | Text pattern scanning and processing tool to handle big data in text format in many different ways | 4.0.2 | Copyright © 2016 Free Software Foundation, Inc. |
| | sort | Sorting information of a text file in many different ways | 8.22 | |
| | grep | Handling and processing big data in text format in many different ways | 2.20 | |

**Supplementary Table S1. Details on the applied tools.**

### 2.1. Preprocessing

Raw sequencing data for both input and enriched samples were first quality checked (using FastQC) and trimmed (using Trimmomatic (8)), then aligned to the human reference genome (using BWA (9)). The GRCh38.d1.vd1 reference genome sequence (basically the GCA_000001405.15_GRCh38_no_alt_analysis_set (10)) was selected that contains additional decoy segments (GenBank Accession GCA_000786075) and virus sequences to help eliminating potential contaminating reads from the core alignment (https://gdc.cancer.gov/about-data/data-harmonization-and-generation/gdc-reference-files (11)). Aligned reads were sorted (using samtools sort (12)), and duplicates were marked (using Picard Tools) resulting in bam files (raw aligned reads). Reads with MAPQ=0 were removed from raw bam files using samtools view as follows.

```
$ samtools view -b -h -q 1 NAME.sorted.dedup.bam -L list_of_chr_bam.bed -o
NAME.MAPQfiltered.bam
```

*# list_of_chr_bam.bed is a 3 column tab delimited text file with indication of the name of chromosomes, their starts and their ends within the applied reference genome assembly*.

Hereafter, all applied command lines are provided in a generalized way, where „NAME" consists of the following indications: treatments_cellType_replicationNo_sampleType. In this study, „treatments" can be WT, NT_UGI, RTX_UGI, or 5FdUR_UGI; cellTypes can be HCT116 or K562; replicationNo can be rep1, rep2, or merged; sampleType can be IP (=enriched), son (=input), or combination of these in case of log2 ratio or other files derived from two samples (e.g. IP_vs_son). Where distinction is necessary, a note is inserted in brackets after the „NAME" (e.g. NAME(rep1)…), otherwise the command was applied on all of the samples. The names of the files deposited into the Gene Expression Omnibus (GEO, accession number GSE126822) also follow this scheme.

Cell type specific blacklists were created by combination of the universal DAC blacklist (https://www.encodeproject.org/files/ENCFF419RSJ) suggested for general use by ENCODE consortium (2) and a cell type specific blacklist defined based on ultrahigh signal (UHS) regions and low mappability regions detected in the input sequencing data (Supplementary Figure S3). This procedure involves deepTools (3), some tools from the kentUtils package of the UCSC (13), R and linux command-line utilities. The steps are as follows:

**Method to define Ultra High Signal (UHS) regions:**

(1) Compute coverage tracks without smoothing for input samples only.

```
$ bamCoverage -b NAME.sorted.dedup.bam -o NAME.bin100bp.no_smooth.RPGC.bw --binSize
100 --verbose --normalizeUsing RPGC --effectiveGenomeSize 2913022398 -p 16
```

(2) Compute histogram on coverage signals to define a threshold above which UHS regions are considered (Supplementary Figure S3C).

```
$ bigWigToWig NAME.bin100bp.no_smooth.RPGC.bw NAME.bin100bp.no_smooth.RPGC.wig
```

*In R:*

```
> NAME <- read.delim("NAME.bin100bp.no_smooth.RPGC.wig", header=FALSE)
> hist(NAME$V4, breaks = 100)
> hist(NAME$V4, breaks = 3000, xlim = c(-0.2, 400), ylim = c(0, 5000))
```

A threshold at coverage signal = 50 was decided.

(3) Compute interval (bed) files describing UHS regions.

*# deleting lines that are only for indication the bedGraph sections and then selecting data bins that are above the threshold 50*

```
$ grep -vwF "bedGraph" NAME.bin100bp.no_smooth.RPGC.wig | awk ' $4 > 50 ' >
NAME.bin100bp.no_smooth.RPGC.UHS.bed
```

*# merging neighbouring data bins to a single interval, then sorting, then printing column 1, 2, and 3, and also the line number in each line of the bed file*

```
$ bedtools merge -i NAME.bin100bp.no_smooth.RPGC.UHS.bed | sort -k1,1 -k2,2n | awk
'{print $1 "\t" $2 "\t" $3 "\t" NR}' > NAME.bin100bp.no_smooth.RPGC.UHS.numbered.bed
```

*# calculating average log2 uracil enrichment value for the intervals in the bed file, it is added to the column 5*

```
$ bigWigAverageOverBed -bedOut=NAME.bin100bp.no_smooth.RPGC.UHS.scored.bed
NAME.bin100bp.no_smooth.RPGC.bw NAME.bin100bp.no_smooth.RPGC.UHS.numbered.bed DEL.tab
```

*# sorting, then printing again with the right format of the float numbers in the column 5*

```
$ sort -k1,1 -k2,2n NAME.bin100bp.no_smooth.RPGC.UHS.scored.bed | awk '{printf "%s\t",
$1; printf "%s\t", $2; printf "%s\t", $3; printf "%s\t", $4; printf "%f\n", $5}' >
NAME.bin100bp.no_smooth.RPGC.UHS.scored2.bed
```

**Method to define low-mappability regions:**

(1) Compute coverage tracks without smoothing and also without normalizing for the input bam files, original and filtered ones (in filtered one, the MAPQ=0 reads were removed using samtools view, see above).

```
$ bamCoverage -b NAME.sorted.dedup.bam -o NAME.bin100bp.no_smooth.no_norm.bw --binSize
100 --verbose  --effectiveGenomeSize 2913022398 –p 16
```

```
$ bamCoverage -b NAME.MAPQfiltered.bam -o
NAME.MAPQfiltered.bin100bp.no_smooth.no_norm.bw --binSize 100 --verbose  --
effectiveGenomeSize 2913022398 –p 16
```

(2) Compute log2 ratio track of coverage original / filtered (using deepTools/BamCompare for bins 100bp).

```
$ bigwigCompare -b1 NAME.bin100bp.no_smooth.no_norm.bw -b2
NAME.MAPQfiltered.bin100bp.no_smooth.no_norm.bw -o NAME.original_vs_filtered.log2.bw -
of bigwig --binSize 100 --skipZeroOverZero --pseudocount 2 1 -v -p 16
```

(3) Compute histogram on log2 ratio signals (Supplementary Figure S3D).

```
$ bigWigToWig NAME.original_vs_filtered.log2.bw NAME.original_vs_filtered.log2.wig
```

*In R* (14)*:*

```
> NAME_of <- read.delim("NAME.original_vs_filtered.log2.wig", header=FALSE)
> hist(NAME_of$V4, breaks = 100)
> hist(NAME_of$V4, breaks = 100, xlim = c(-0.2, 4), ylim = c(0, 1500000))
```

A threshold at log2 ratio signal = 1.0 was decided, that means that half of the reads in the given bin have MAPQ=0.

(4) Compute interval (bed) files that describe regions with more than 50% ambiguously mapped reads considered as low-mappability regions.

```
$  grep -vwF "bedGraph" NAME.original_vs_filtered.log2.wig | awk ' $4 > 1 ' >
NAME.original_vs_filtered.log2.blackMAPQ.bed
```

```
$ bedtools merge -i NAME.original_vs_filtered.log2.blackMAPQ.bed | sort -k1,1 -k2,2n |
awk '{print $1 "\t" $2 "\t" $3 "\t" NR}' >
NAME.original_vs_filtered.log2.blackMAPQ.numbered.bed
```

```
$ bigWigAverageOverBed -bedOut=NAME.original_vs_filtered.log2.blackMAPQ.scored.bed
NAME.original_vs_filtered.log2.bw
NAME.original_vs_filtered.log2.blackMAPQ.numbered.bed DEL.tab
```

```
$ sort -k1,1 -k2,2n NAME.original_vs_filtered.log2.blackMAPQ.scored.bed | awk '{printf
"%s\t", $1; printf "%s\t", $2; printf "%s\t", $3; printf "%s\t", $4; printf "%f\n",
$5}' > NAME.original_vs_filtered.log2.blackMAPQ.scored2.bed
```

Cell type specific blacklists were then created by merging the DAC blacklist (ENCFF419RSJ), the UHS and the low-mappability regions using bedtools merge with the parameter - d500 to avoid 500 bases or shorter gaps with obviously no biological meaning (cf. purple and black intervals on IGV view at Supplementary Figure S3B). For HCT116 cell line specific blacklist, all the corresponding input samples were used and the derived intervals were merged together.

```
$ cat NAME1.bin100bp.no_smooth.RPGC.UHS.scored2.bed
NAME2.bin100bp.no_smooth.RPGC.UHS.scored2.bed {…}
```

```
NAMEn.bin100bp.no_smooth.RPGC.UHS.scored2.bed | sort -k1,1 -k2,2n >
united_sorted_UHS_HCT116.bed
```

```
$ bedtools merge -i united_sorted_UHS_HCT116.bed > UHS_HCT116.bed
```

```
$ cat NAME1.original_vs_filtered.log2.blackMAPQ.scored2.bed
NAME2.original_vs_filtered.log2.blackMAPQ.scored2.bed {…}
NAMEn.original_vs_filtered.log2.blackMAPQ.scored2.bed | sort -k1,1 -k2,2n >
united_sorted_blackMAPQ_HCT116.bed
```

```
$ bedtools merge -i united_sorted_blackMAPQ_HCT116.bed > blackMAPQ_HCT116.bed
```

```
$ cat ENCFF419RSJ.bed UHS_HCT116.bed blackMAPQ_HCT116.bed | sort -k1,1 -k2,2n >
united_sorted_blacklist_HCT116.bed
```

```
$ bedtools merge -i united_sorted_blacklist_HCT116.bed -d 500 > blacklist_HCT116.bed
```

The effective genome size was calculated by subtracting the blacklisted and the originally masked regions of the reference genome.

```
$ bedtools subtract -a list_of_chr_bam.bed -b blacklist_HCT116.bed >
not_blacklisted_HCT116.bed
```

```
$ bedtools nuc -fi GRCh38.d1.vd1.fa -bed not_blacklisted_HCT116.bed >
not_blacklisted_HCT116_nuc.bed
```

```
$ awk '{(sum1+=$6) (sum2+=$9) (sum3+=$7) (sum4+=$8) (sum5+=$10) (sum6+=$11)
(sum7+=$12)} END {print sum1 "\t" sum2"\t" sum3 "\t" sum4 "\t" sum5 "\t" sum6 "\t"
sum7}' not_blacklisted_HCT116_nuc.bed
```

```
825630405    826937345    570444697    570830493    165010872    99    2958853872
```

*# Note that awk will sum up the number from the head line too – so column number has to be subtracted.*

| number of A | number of T | number of C | number of G | number of N | Nº other | length |
|---|---|---|---|---|---|---|
| 825630399 | 826937336 | 570444690 | 570830485 | 165010862 | 88 | 2958853860 |

Thereby, the effective genome size was calculated for the analysis of the HCT116 samples as 2793842910 (length – number of N – Nº other).

GC content for the effective part of the reference genome was found to be 40.85%, calculated according to the formula: (number of  C + number of G) / effective genome size.

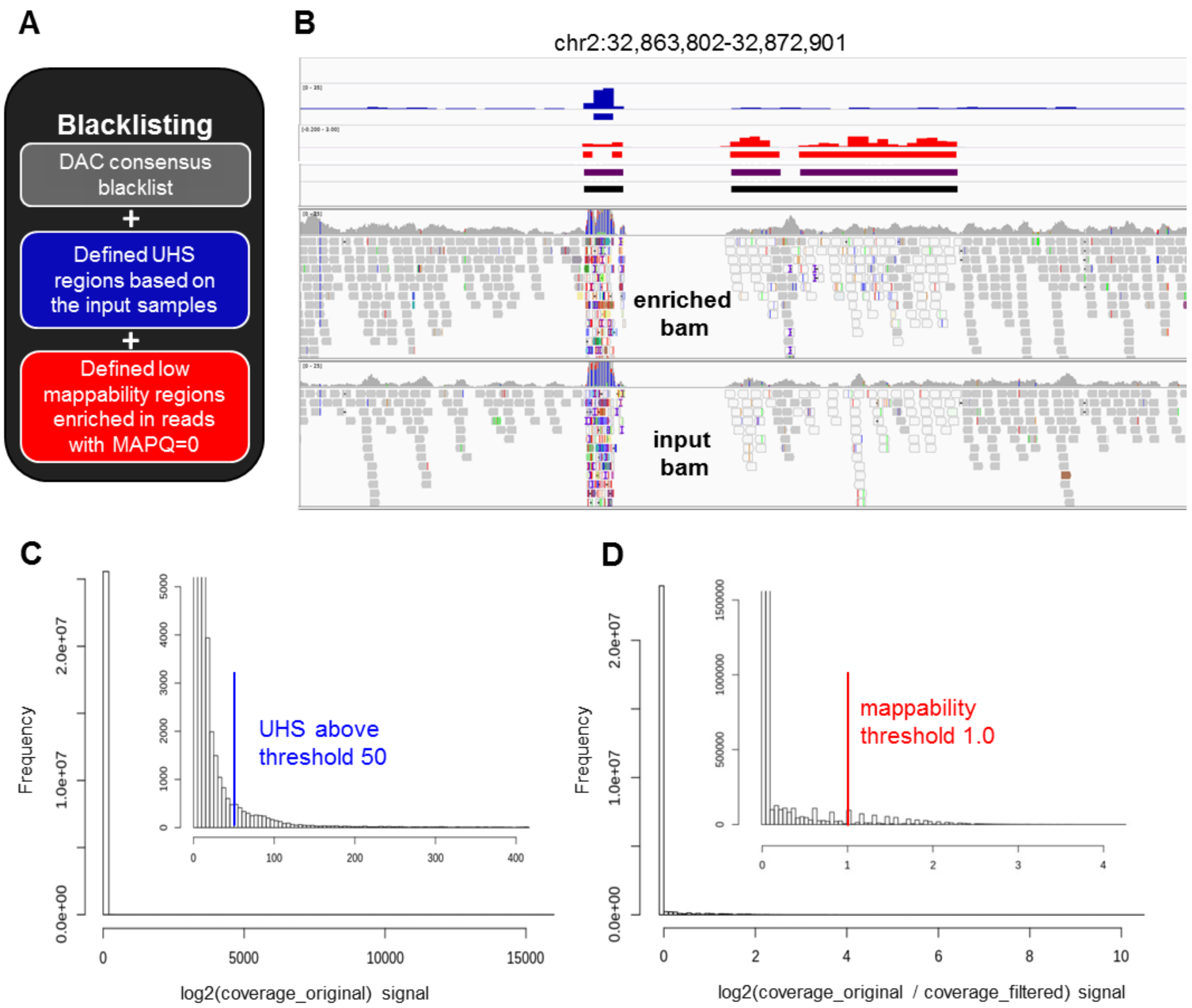

**Supplementary Figure S3. Creation of cell line specific blacklists. (A)** Concept of blacklisting. United cell line specific blacklist was composed from universal DAC blacklist (https://www.encodeproject.org/files/ENCFF419RSJ, grey), input sample derived Ultra High Signal (UHS, blue) regions and low mappability regions where more than 50% of the reads are ambiguously mapped (MAPQ=0, red). **(B)** Representative IGV view of genomic region (chr2:32,863,802-32,872,901) where both UHS and low mappability regions are present. Tracks from the top are as follows: DAC blacklist (no intervals in the presented region); coverage track (bw file, blue), and the derived UHS regions (bed file, blue); log2 ratio (original/filtered) track (bw file, red), and the derived low mappability regions (bed file, red); intervals after combination of DAC, UHS, and low mappability regions (bed file, purple); and the same after merging intervals separated by less than 500 bases (bed file, the final blacklist, black). The bw files and the derived bed files are shown for one input sample, while the blacklists presented here are calculated from all of the input data corresponding to HCT116 cell line. Raw alignments (bam files) of the corresponding enriched and input samples are also shown. Reads in white are characterized with MAPQ=0. Alterations in reads comparing to the reference genome are indicated with different colours according to the IGV standard. In regions suffering from alignment artefacts, reads often appears with multiple alterations, as it is visible here for the UHS region. **(C)** Histogram of coverage signal of a representative input sample (the same as above). Both full data range and a zoomed version (inset) with better resolution are shown. In the zoomed histogram, the defined UHS threshold (blue) is also indicated. **(D)** Histogram of log2 ratio (original/filtered) signal of a representative input sample (the same as above). Both full data range and a zoomed version (inset) with better resolution are shown. In the zoomed histogram, the defined low-mappability threshold (red) is also indicated.

This cell type specific united blacklist was applied in samtools view to BAM files that were also filtered for MAPQ=0 reads previously.

```
$ samtools view -b -h NAME.MAPQfiltered.bam -L blacklist_HCT116.bed -o
NAME.blacklist.bam -U NAME.filtered_blacklisted.bam

$ samtools index NAME.filtered_blacklisted.bam

$ samtools idxstats NAME.filtered_blacklisted.bam >
NAME.filtered_blacklisted.bam.idxstats.csv
```

| sample | replicates | number of raw reads | number of mapped reads | unmapped reads | | uniquely mapped reads | | uniquely mapped reads after blacklisting | |
|---|---|---|---|---|---|---|---|---|---|
| | | | | number | % | number | % | number | % |
| WT input | WT1_son | 138283424 | 138113944 | 169480 | 0.12 | 131604925 | 95.17 | 126302380 | 91.34 |
| | WT2_son | 185174607 | 184959442 | 215165 | 0.12 | 175302618 | 94.67 | 168698159 | 91.10 |
| WT enriched | WT1_IP | 144612745 | 144094135 | 518610 | 0.36 | 138611548 | 95.85 | 131765827 | 91.12 |
| | WT2_IP | 159514985 | 159314208 | 200777 | 0.13 | 152796029 | 95.79 | 145489972 | 91.21 |
| NT_UGI input | NT1_son | 164023406 | 163757733 | 265673 | 0.16 | 156045404 | 95.14 | 149734348 | 91.29 |
| | NT2_son | 173254485 | 173088530 | 165955 | 0.10 | 165373102 | 95.45 | 158978819 | 91.76 |
| NT_UGI enriched | NT1_IP | 260763674 | 260300247 | 463427 | 0.18 | 251164014 | 96.32 | 239327438 | 91.78 |
| | NT2_IP | 136148357 | 134759365 | 1388992 | 1.02 | 129486254 | 95.11 | 123064064 | 90.39 |
| 5FdUR_UGI input | 5FdUR1_son | 128706895 | 128669770 | 37125 | 0.03 | 122476766 | 95.16 | 118558597 | 92.12 |
| | 5FdUR1_son | 201926203 | 201560665 | 365538 | 0.18 | 193086643 | 95.62 | 184756297 | 91.50 |
| 5FdUR_UGI enriched | 5FdUR1_IP | 150596242 | 150522522 | 73720 | 0.05 | 144554269 | 95.99 | 141582874 | 94.01 |
| | 5FdUR2_IP | 138651760 | 138410833 | 240927 | 0.17 | 133200761 | 96.07 | 128584894 | 92.74 |
| RTX_UGI input | RTX1_son | 145920877 | 145775676 | 145201 | 0.10 | 139168642 | 95.37 | 133567232 | 91.53 |
| | RTX2_son | 147882518 | 147674678 | 207840 | 0.14 | 141097936 | 95.41 | 135259752 | 91.46 |
| RTX_UGI enriched | RTX1_IP | 166544868 | 166305588 | 239280 | 0.14 | 160567280 | 96.41 | 155171205 | 93.17 |
| | RTX2_IP | 151875638 | 151666578 | 209060 | 0.14 | 146619425 | 96.54 | 141987664 | 93.49 |
| K562 input | K562_son | 106137622 | 105875437 | 262185 | 0.25 | 100326105 | 94.52 | 97429855 | 91.8 |
| K562 enriched | K562_IP | 109490393 | 109306854 | 183539 | 0.17 | 105310296 | 96.18 | 102013265 | 93.17 |

**Supplementary Table S2. Number of reads in samples during the pre-processing steps.** All samples and replicates are shown here that were sequenced in the frame of the present publication. Number of raw reads means read number before starting alignment (the sum of the mapped and unmapped read numbers). Uniquely mapped read means that MAPQ is not zero. The samples are as follows: non-treated wild-type HCT116 cells (WT), non-treated UGI-expressing HCT116 cells (NT_UGI), 5FdUR treated UGI-expressing HCT116 cells (5FdUR_UGI), RTX treated UGI-expressing HCT116 cells (RTX_UGI), and non-treated wild-type K562 cells (K562). Genomic DNA was isolated and sonicated to about 300 kb fragments (input), uracil-DNA was enriched by immunoprecipitation via FLAG-tagged U-DNA sensor (enriched). Here, we included K562 data too that was addressed to have a kind of reference point to the previously published dU-seq data (15) with which detailed comparison is also made at the end of the supplementary material (cf. Supplementary Figure S11, S12 and Supplementary Table S5).

Correlation can be calculated among bam files using multiBamSummary and plotCorrelation tools of the deepTools package (3). Pearson correlation coefficients were calculated with 5000 bases bin size between uniquely mapped reads of samples after blacklisting as follows:

```
$ multiBamSummary bins --binSize 5000 -b NAME1.filtered_blacklisted.bam
NAME2.filtered_blacklisted.bam {…} NAMEn.filtered_blacklisted.bam  -o
multiBamSummary_bin5000.npz --scalingFactors
scalingFactors_from_multiBamSummary_bin5000.txt --outRawCounts
raw_counts_from_multiBamSummary_bin5000.csv --ignoreDuplicates --maxFragmentLength
2000 --extendReads -v -p 16
```

```
$ plotCorrelation --corData multiBamSummary_bin5000.npz --corMethod pearson --
whatToPlot heatmap -o multiBamSummary_bin5000_heatmap.png -T multiBamSummary_bin5000 -
-skipZeros --removeOutliers --plotNumbers --colorMap RdPu
```

Pearson correlation coefficients between replicates were measured as follows: WT enriched: 0.92, input: 0.89; NT_UGI enriched: 0.79, input: 0.82; 5FdUR_UGI enriched: 0.87, input: 0.88; RTX_UGI enriched: 0.97, input: 0.89. All further data processing and analysis steps were done on the two biological replicates separately, as well as on merged bam files of corresponding replicates. All the results were in good agreement between replicates, so hereafter, in the main figures, we show results for the merged data.

Merging replicates were performed at the level of cleaned aligned reads (filtered_blacklisted.bam files) using samtools merge (12).

```
$ samtools merge -r -1 -c --threads 16
NAME(merged).filtered_blacklisted.non_sorted.bam NAME(rep1).filtered_blacklisted.bam
NAME(rep2).filtered_blacklisted.bam
```

```
$ samtools sort -l1 -o NAME(merged).filtered_blacklisted.bam -O BAM -@16
NAME(merged).filtered_blacklisted.non_sorted.bam
```

```
$ samtools index NAME(merged).filtered_blacklisted.bam
```

Comparison of the samples at the level of merged, filtered and blacklisted bam files (Supplementary Figure S4) shows clear differences among input and enriched files, as well as treated and non-treated samples. All input files belong to the HCT116 cell line are quite similar, while the input sample of K562 cells shows significant difference that is another argument for cell type specific blacklisting.

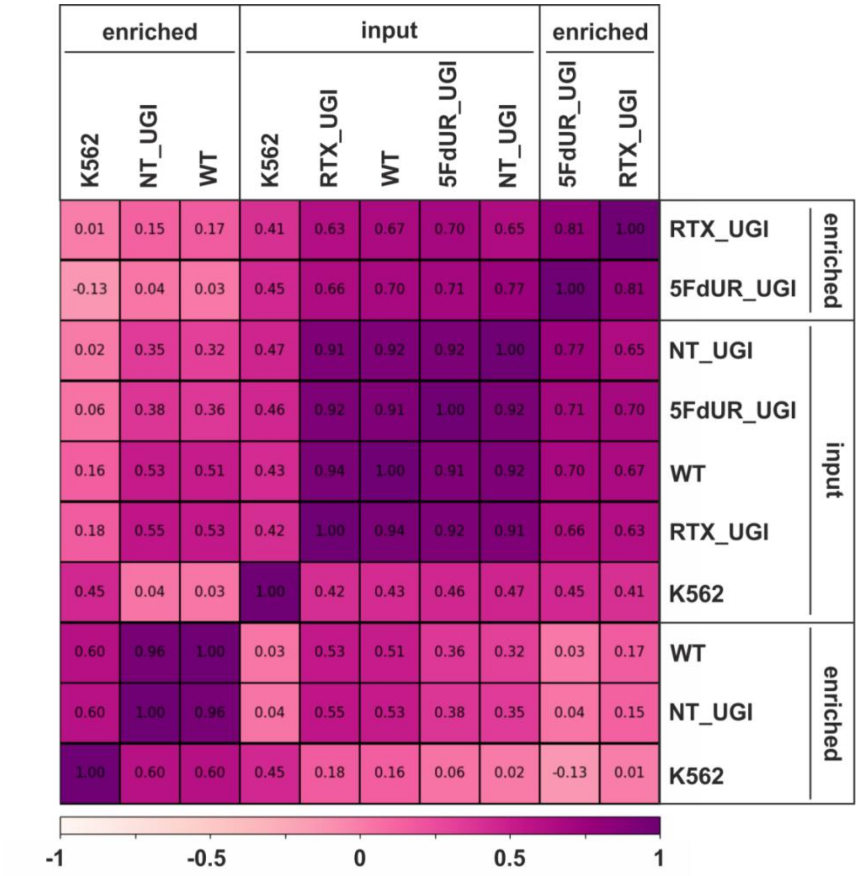

**Supplementary Figure S4. Pearson correlation among processed BAM files.** Calculation was done on uniquely mapped reads after blacklisting and merging replicates, using deepTools multiBamSummary, and heatmap was plotted using plotCorrelation as described in the text above. The input and the corresponding enriched samples are as follows: non-treated wild-type HCT116 cells (WT), non-treated UGI-expressing HCT116 cells (NT_UGI), 5FdUR treated UGI-expressing HCT116 cells (5FdUR_UGI), RTX treated UGI-expressing HCT116 cells (RTX_UGI), and non-treated wild-type K562 cells (K562).

#### 2.2. Determination of uracil enrichment: log2 ratio track and derived regions versus peaks called by MACS2 tool.

Uracil enrichment should be determined from the increased coverage of enriched data versus the input using cleaned aligned reads (filtered_blacklisted.bam files). For that, basically two main ways are available: 1) conventional peak calling algorithms (e.g. MACS2), especially if relatively intense and sharp peaks of enrichment are expected; 2) calculation and comparison of genome scaled coverage tracks for both enriched and input sequencing data e.g. in the form of log2 ratio tracks (Supplementary Figure S5). This latter option results in more detailed information on the enrichment in the format of bedgraph or bigwig (bw). However, such log2 ratio tracks (bw files) can hardly be used to screen large databases for colocalizing genomic features or factor binding profiles (cf. Supplementary Figure S2).

In case of the present samples (either non-treated or treated by thymidylate biosynthesis inhibitors), we found broad genomic regions with elevated log2 signals rather than intense sharp peaks (Figure 3A, Supplementary Figure S5, Supplementary Figure S6). Hence, we decided to derive interval (bed) files from the log2 ratio tracks (bw) using a threshold reasonable based on log2 ratio signal histograms (cf. Figure 3C). These intervals might be able to describe such broad regions of uracil enrichment better than the peak calling results (cf. Supplementary Figure S5), and simultaneously allow efficient screening of large datasets for colocalizing features.

To further access the appropriate approach of data processing and extracting information on genomic uracil enrichment, we performed both 1) broad peak calling, and 2) extraction of even broader regions based on log2 ratio tracks. Hereafter, the two terms 'peak' and 'region' will be consequently applied for the results of these two approaches, respectively.

1) Peak calling was performed using broad peak option in MACS2 at two different broad-cutoff values (grey intervals at Supplementary Figure S5). Note that --cutoff-analysis option can also be used to estimate the number and length of the peaks at different q and p cutoff values.

```
$ MACS2 callpeak -t NAME(IP).filtered_blacklisted.bam -c
NAME(son).filtered_blacklis.bam --broad -g 2793842910 --broad-cutoff 0.05 -n NAME.0p05
--outdir {PATH} --nomodel -f BAMPE
```

```
$ MACS2 callpeak -t NAME(IP).filtered_blacklisted.bam -c
NAME(son).filtered_blacklis.bam --broad -g 2793842910 --broad-cutoff 0.5 -n NAME.0p5 -
-outdir {PATH} --nomodel -f BAMPE
```

2) Determination of broad regions based on log2 ratio tracks was performed as follows using bamCoverage and bigwigCompare tools of deepTools package (3), some tools from the kentUtils package of the UCSC (13), R and linux command-line utilities.

```
$ bamCoverage -b NAME.filtered_blacklisted.bam -o NAME.bin100bp.smooth5000.RPGC.bw --
binSize 100 --verbose --smoothLength 5000 --normalizeUsing RPGC --effectiveGenomeSize
2793842910 -p 16 --extendReads
```

```
$ bigwigCompare -b1 NAME(IP).bin100bp.smooth5000.RPGC.bw -b2
NAME(son).bin100bp.smooth5000.RPGC.bw -o NAME.bin100bp.smooth5000.RPGC.log2.bw -of
bigwig --binSize 100 -v -p 16
```

```
$ bigWigToWig NAME.bin100bp.smooth5000.RPGC.log2.bw
NAME.bin100bp.smooth5000.RPGC.log2.wig
```

*In R (Figure 3C):*

```
> NAME(short) <- read.delim("NAME.bin100bp.smooth5000.RPGC.log2.wig", header=FALSE)
> hist(NAME(short)$V4, breaks = 100, xlim = c(-1.5, 1.5), ylim = c(0, 2500000))
```

The histograms are shown in Figure 3C. The applied thresholds are shown in Supplementary Table S3A.

Extraction of the data bins with log2 ratio signal higher than the threshold was done as follows:

*# deleting lines that is only for indication the bedGraph sections and then selecting data bins that are above the threshold (in this example, it is 0.2)*

```
$ grep -vwF "bedGraph" NAME.bin100bp.smooth5000.RPGC.log2.wig | awk ' $4 > 0.2 ' >
NAME.bin100bp.smooth5000.RPGC.log2.0p2.bed
```

*# merging neighbouring data bins to a single interval, then sorting, then printing column 1, 2, and 3, and also the line number in each line of the bed file*

```
$ bedtools merge -i NAME.bin100bp.smooth5000.RPGC.log2.0p2.bed | sort -k1,1 -k2,2n |
awk '{print $1 "\t" $2 "\t" $3 "\t" NR}' >
NAME.bin100bp.smooth5000.RPGC.log2.0p2.numbered.bed
```

*# calculating average log2 uracil enrichment value for the intervals in the bed file, it is added to the column 5*

```
$ bigWigAverageOverBed -bedOut=NAME.bin100bp.smooth5000.RPGC.log2.0p2.scored.bed
NAME.bin100bp.smooth5000.RPGC.log2.bw
NAME.bin100bp.smooth5000.RPGC.log2.0p2.numbered.bed DEL.tab
```

*# sorting, then printing again with the right format of the float numbers in the column 5*

```
$ sort -k1,1 -k2,2n NAME.bin100bp.smooth5000.RPGC.log2.0p2.scored.bed | awk '{printf
"%s\t", $1; printf "%s\t", $2; printf "%s\t", $3; printf "%s\t", $4; printf "%f\n",
$5}'  > NAME.bin100bp.smooth5000.RPGC.log2.0p2.region.bed
```

*# only if top ranked intervals have to be selected: sorting by average log2 uracil enrichment scores in decreasing order, then selecting the top 50000 intervals (other numbers of top intervals can be defined as it is desired), then sorting back in alphabetic order (that is required by several possible further applications e.g. bedtools)*

```
$ sort -k 5 -nr NAME.bin100bp.smooth5000.RPGC.log2.0p2.region.bed | head -n 50000 |
sort -k1,1 -k2,2n > NAME.bin100bp.smooth5000.RPGC.log2.0p2.top50k.bed
```

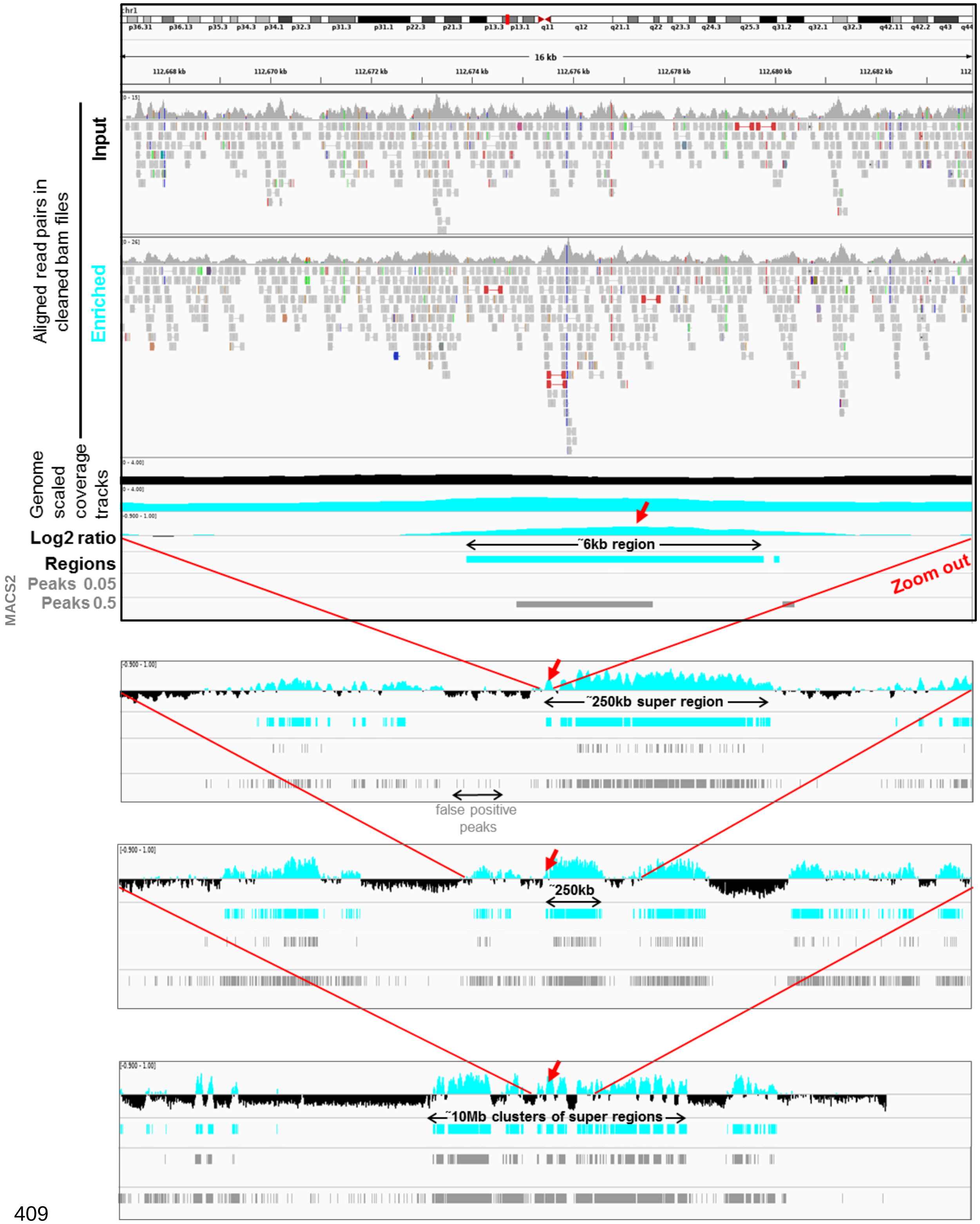

**Supplementary Figure S5. Representative IGV views from aligned reads to a 10 Mb cluster of uracil enriched regions compared to usual peak calling results.** Upper panel shows IGV view of 112.667.000-112.684.000 genomic segment to demonstrate the processing steps. The tracks from the top are as follows: cleaned aligned reads (bam files) for a pair of input (black title) and enriched (cyan title) samples; genome scaled coverage tracks for the input (black), and for the enriched (cyan) samples computed by deepTools/bamCoverage; log2 ratio track (enriched coverage / input coverage, positive values are shown in cyan, negative values are in black, computed by deepTools/bigwigCompare); regions of uracil enrichment derived from log2 ratio using a threshold, as described above (cyan); peaks called by MACS2 with 0.05 and also with the more permissive 0.5 broad-cutoff (grey). In the presented 16 kb genomic segment a 6 kb region of uracil enrichment is detected, while peak calling either fails or detects only a shorter peak depending on the applied broad-cutoff value. The following panels are zooming out on the same 6 kb region (red arrows) allowing visualization of its broader context. This 6 kb region of uracil enrichment (red arrows) is a part of a 250 kb super-region that is surrounded by similar super-regions separated by negative log2 value segments. On a larger scale, even these super-regions are clustered (to a 10 Mb cluster here, bottom panel). It is also visible that using more permissive broad-cutoff in peak calling allows higher coverage of the uracil enriched regions, however detects false positive peaks as well.

We argue that peak calling using MACS2 is suboptimal for description of distribution of genomic uracil, even if broad peak calling is applied (Supplementary Figure S5). Based on theoretical expectations (cf. main text) as well as on the initial processing of the actual U-DNA-Seq data, we recommend to use the log2 ratio of the genome scaled coverage tracks and the derived regions of uracil enrichment rather than the peak calling approach.

To further strengthen this choice, we made a detailed comparison on the defined regions of uracil enrichment (based on log2 ratio tracks) and the peak calling results (Supplementary Table S3). A statistics, including the applied thresholds, Jaccard indices between replicates, and the extent of the regions, are shown for region.bed files derived from the log2 ratio tracks (Supplementary Table S3A). Regarding peak calling, we found, that using the same broad-cutoff parameter, the numbers of called peaks are extremely different (from 35 000 to 250 000) among the samples, even between parallels. This difference in peak numbers does not seem to correlate with the elevated uracil level in treated samples (cf. higher number of peaks in WT and NT_UGI samples than in the treated ones). Using the „--cutoff-analysis" option in MACS2, we tried to harmonize the number of called peaks in different samples using sometimes very different broad-cutoff parameters (Supplementary Table S3B). Comparing the two statistics for the two approaches, the reproducibility of peak calling was still much worse (cf. Jaccard index values between replicates, in case of peak calling (Supplementary Table S3B) versus log2 regions of uracil enrichment (Supplementary Table S3A)). Lower reproducibility of peak calling results in lower descriptive value for the uracil distribution, as it is also reflected in comparison of drug-treated and non-treated samples (Supplementary Table S3D vs S3C).

Overlapping bases and Jaccard indices were calculated for the interval files by bedtools jaccard tool as follows:

```
$ bedtools jaccard -a NAME1.bin100bp.smooth5000.RPGC.log2.0p2.region.bed -b
NAME2.bin100bp.smooth5000.RPGC.log2.0p2.region.bed
```

**A**

| sample | replicates | treshold | number of regions | number of bp in regions | Jaccard index | Jaccard indext to the merge | GC content within the intervals % |
|---|---|---|---|---|---|---|---|
| WT | WT_HCT116_rep1 | 0.33 | 137,224 | 331,725,091 | 0.37 | 0.63 | 33.8 |
| | WT_HCT116_rep2 | 0.25 | 145,757 | 328,484,272 | | 0.61 | 33.9 |
| | WT_HCT116_merged | 0.27 | 121,310 | 319,476,171 | | | 33.4 |
| NT_UGI | NT_UGI_HCT116_rep1 | 0.29 | 119,216 | 321,518,016 | 0.37 | 0.71 | 33.5 |
| | NT_UGI_HCT116_rep2 | 0.32 | 140,951 | 325,942,833 | | 0.53 | 33.9 |
| | NT_UGI_HCT116_merged | 0.33 | 110,035 | 327,691,223 | | | 33.3 |
| 5FdUR_UGI | 5FdUR_UGI_HCT116_rep1 | 0.25 | 99,303 | 510,748,006 | 0.42 | 0.68 | 46.1 |
| | 5FdUR_UGI_HCT116_rep2 | 0.14 | 150,860 | 503,980,890 | | 0.62 | 46.2 |
| | 5FdUR_UGI_HCT116_merged | 0.18 | 90,286 | 525,051,904 | | | 46.5 |
| RTX_UGI | RTX_UGI_HCT116_rep1 | 0.25 | 96,960 | 506,205,489 | 0.63 | 0.78 | 44.8 |
| | RTX_UGI_HCT116_rep2 | 0.25 | 84,788 | 555,525,646 | | 0.81 | 45.2 |
| | RTX_UGI_HCT116_merged | 0.25 | 65,073 | 520,671,621 | | | 45.2 |
| K562 | K562 | 0.33 | 132,725 | 323,883,311 | | | 33.7 |

**B**

| sample | replicates | broad cutoff | number of peaks | number of bp in peaks | Jaccard index | Jaccard indext to the merge | GC content within the intervals % |
|---|---|---|---|---|---|---|---|
| WT | WT_HCT116_rep1 | 0.045 | 82,110 | 63,007,157 | 0.15 | 0.38 | 29.9 |
| | WT_HCT116_rep2 | 0.07 | 75,415 | 51,232,054 | | 0.35 | 29.9 |
| | WT_HCT116_merged | 0.004 | 82,296 | 59,311,730 | | | 29.2 |
| NT_UGI | NT_UGI_HCT116_rep1 | 0.001 | 74,797 | 74,741,677 | 0.12 | 0.55 | 29.8 |
| | NT_UGI_HCT116_rep2 | 0.35 | 81,788 | 66, 080,148 | | 0.19 | 33.3 |
| | NT_UGI_HCT116_merged | 0.0005 | 84,821 | 80,814,174 | | | 29.9 |
| 5FdUR_UGI | 5FdUR_UGI_HCT116_rep1 | 0.09 | 110,038 | 100,603,863 | 0.16 | 0.37 | 50.7 |
| | 5FdUR_UGI_HCT116_rep2 | 0.15 | 94,120 | 105,426,693 | | 0.40 | 52.8 |
| | 5FdUR_UGI_HCT116_merged | 0.03 | 100,289 | 117,311,431 | | | 52.5 |
| RTX_UGI | RTX_UGI_HCT116_rep1 | 0.15 | 86,529 | 123,079,165 | 0.22 | 0.43 | 44.5 |
| | RTX_UGI_HCT116_rep2 | 0.08 | 78,301 | 121,000,794 | | 0.44 | 45.5 |
| | RTX_UGI_HCT116_merged | 0.015 | 73,063 | 133,717,635 | | | 45.1 |
| K562 | K562 | 0.03 | 96,886 | 112,539,705 | | | 31.6 |

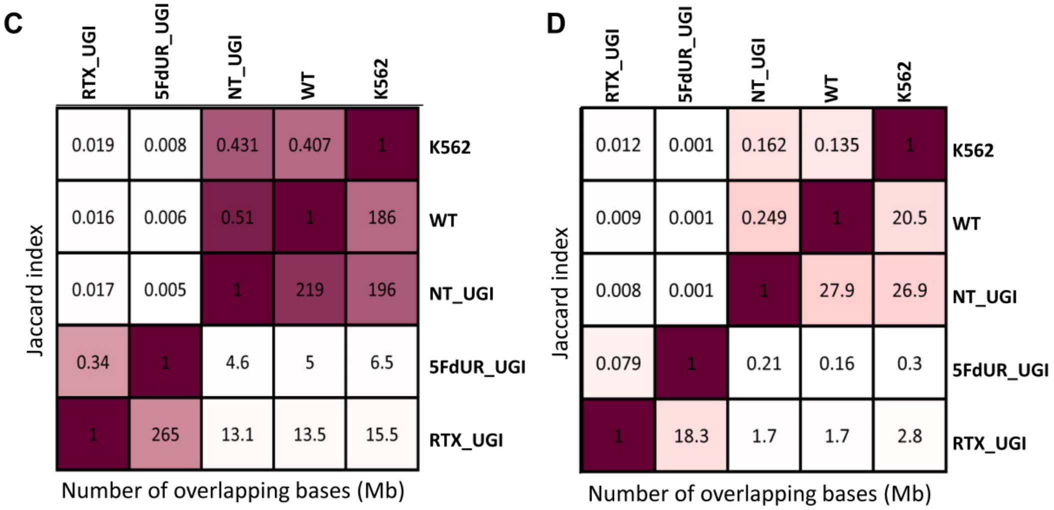

**Supplementary Table S3. Comparison of regions of uracil enrichment derived from log2 signal ratio and peaks called by MACS2.** The samples are as follows: non-treated wild-type (WT), non-treated UGI-expressing (NT_UGI), 5FdUR treated UGI-expressing (5FdUR_UGI), RTX treated UGI-expressing HCT116 cells (RTX_UGI), non-treated wild-type K562 cells (K562). **(A)** Statistics on log2 ratio derived regions of uracil enrichment. The applied threshold values were fine-tuned to have approximately the same length of intervals in the corresponding replicates and merged data. Bedtools Jaccard indices were calculated between replicates, as well as between individual replicate and the corresponding merge data. GC contents were also calculated for each interval files of regions **(B)** Statistics on peaks called by MACS2. Broad-cutoff values were adjusted to have approximately the same length of intervals in the corresponding replicates and merged data. Bedtools Jaccard indices were calculated between replicates, as well as between individual replicates and the corresponding merged data. GC contents were also calculated for each interval files of peaks. **(C)** Comparison of samples (replicates were merged) based on log2 ratio derived regions. Jaccard indices and numbers of millions of intersecting bases are indicated above and below the diagonal, respectively. Treated and non-treated samples are well separated. **(D)** Comparison of samples (replicates were merged) based on peaks. Jaccard indices and numbers of millions of intersecting bases are indicated above and below the diagonal, respectively. Treated and non-treated samples are not that well separated anymore.

In the QC report of sequencing from Novogene, the GC contents of the sequenced samples were documented. All samples, except the non-treated enriched ones, were around 42% characteristic for the human genome. However, in case of non-treated enriched samples, the GC content was consequently decreased to around 37%. We were curious, if such difference might occur due to different GC content of the regions enriched in uracils in the non-treated versus drug-treated samples. Indeed, GC contents of regions were decreased to around 33% and increased to about 44-46% in case of non-treated and drug-treated samples, respectively (Supplementary Table 3A). For comparison, GC content of the not blacklisted and non-masked part of the reference genome was 40.85% ((number of C + number of G) / effective genome size).

GC% was calculated for the interval files of each sample using bedtools nuc tool and awk as follows:

```
$ bedtools nuc -fi GRCh38.d1.vd1.fa -bed
NAME.bin100bp.smooth5000.RPGC.log2.0p2.region.bed | awk '{(sum1+=$8) (sum2+=$11)
(sum3+=$9) (sum4+=$10)} END {print sum1 "\t" sum2"\t" sum3 "\t" sum4}' >>
summary.region.bed.nuc.csv
```

```
$ bedtools nuc -fi GRCh38.d1.vd1.fa -bed NAME1.0p05_peaks.broadPeak | awk
'{(sum1+=$12) (sum2+=$15) (sum3+=$13) (sum4+=$14)} END {print sum1 "\t" sum2"\t" sum3
"\t" sum4}' >> summary.peaks.bed.nuc.csv
```

Based on the comparison reported in Supplementary Table S3, we decided that log2 ratio tracks and the derived interval files will be used for further analysis. For visualization, IGV views are shown for all the samples (replicates were merged) in a selected genomic region (Figure 3A), as well as for all the chromosomes in Supplementary Figure S6 (the first chromosome is shown here, and the rest is in the Appendix 1).

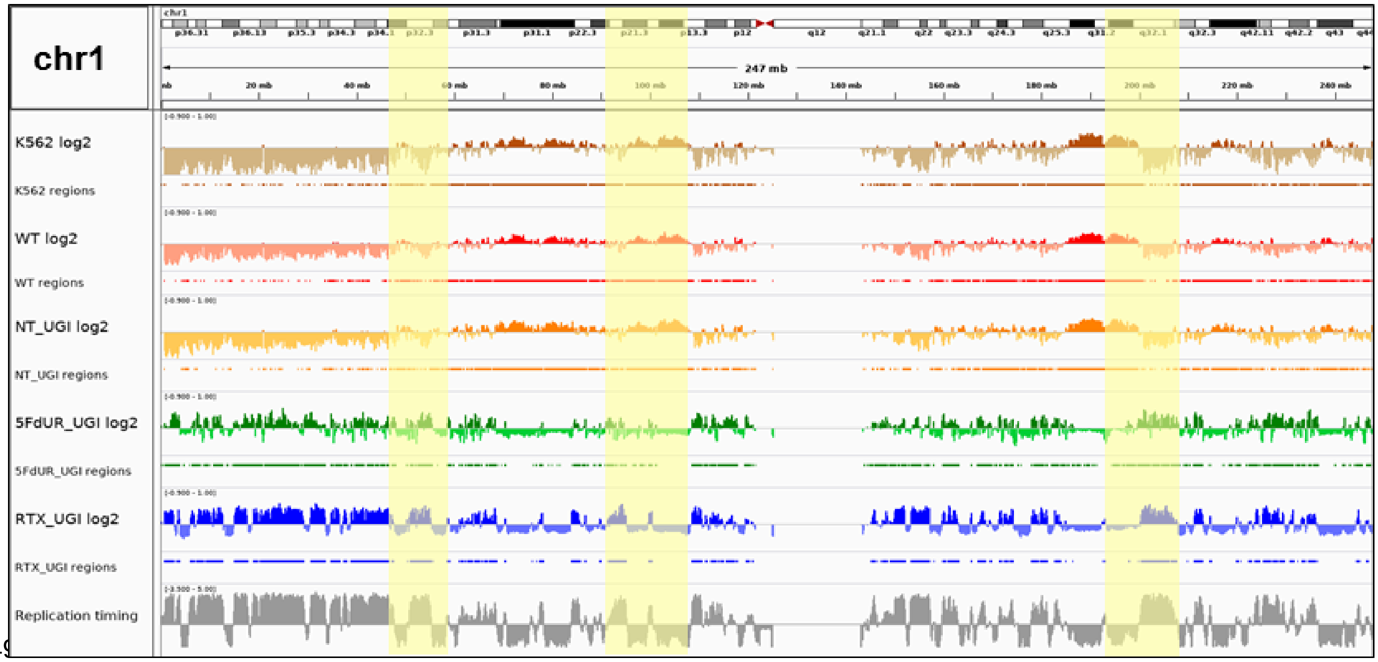

**Supplementary Figure S6. IGV view of log2 ratio and regions of uracil enrichment on chromosome 1 (for all chromosomes see Appendix 1).** At the upper part cytogenetic bands by Giemsa staining are visible, the staining intensity (from white to black) correlates to the chromatin structure. Log2 ratio tracks (enriched coverage / input coverage, computed by deepTools/bigwigCompare) are coloured by samples (non-treated wild type K562 cells (K562) brown, non-treated wild type (WT) red, non-treated UGI-expressing (NT_UGI) orange, 5FdUR treated UGI-expressing (5FdUR_UGI) green, RTX treated UGI-expressing HCT116 cells (RTX_UGI) blue) and negative values are shown with a bit lighter colour than the positives. For all the log2 ratio tracks, the same range was applied from -0.9 to 1. Regions of uracil enrichment derived from log2 ratio tracks are shown with the same colour directly below the corresponding log2 tracks. The bottom track shows replication timing data (grey) for HCT116 downloaded from Replication Domain database (16). Correlation among samples, replication timing and the cytogenetic bands are well visible (highlighted regions, cf. Figure 4B-C, and Supplementary Table S4) and will be discussed in the next session.

Furthermore, we used multiBigwigSummary and plotCorrelation to show Pearson correlation on log2 ratio tracks (see the command lines below). Heatmaps for individual replicates (Supplementary Figure S7) and also for merged replicates (Figure 3B) revealed that the treated and non-treated enriched samples are well separated in terms of global uracil distribution pattern.

```
$ multiBigwigSummary bins -b NAME1.filtered_blacklisted.bw
NAME2.filtered_blacklisted.bw {…} NAMEn.filtered_blacklisted.bw -o
mbws_filtered_blacklisted_bw_data.npz -v -p 16
```

```
$ plotCorrelation --corData mbws_filtered_blacklisted_bw_data.npz --corMethod pearson
--whatToPlot heatmap -o mbws_filtered_blacklisted_bw_heatmap.png -T
mbws_filtered_blacklisted_bw --skipZeros --removeOutliers --plotNumbers --colorMap
RdPu
```

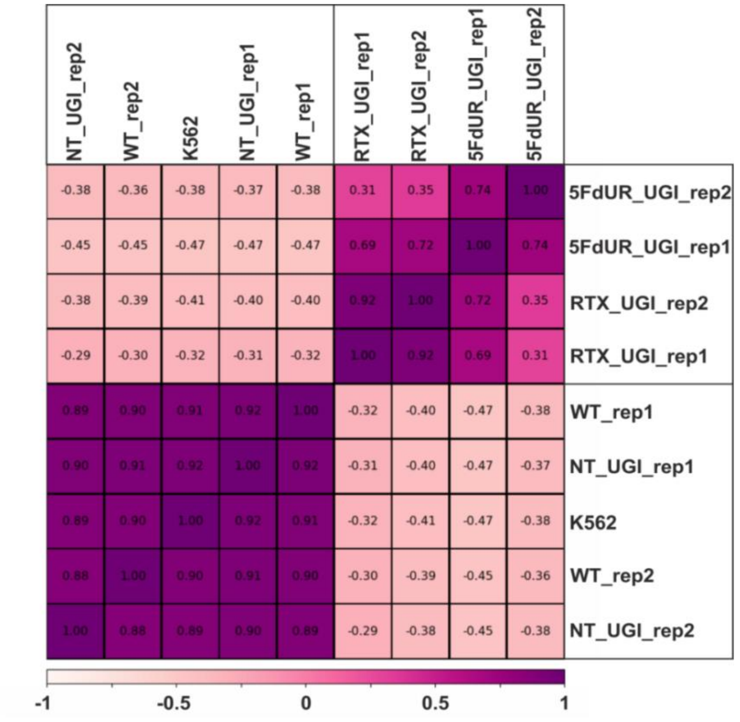

**Supplementary Figure S7. Pearson correlation among log2 ratio tracks of replicates.** The log2 ratio tracks were calculated from the genome scaled coverage tracks (enriched vs input) using deepTools/bigwigCompare tool for the individual replicates. Pearson correlation was calculated by deepTools/multiBigwigSummary and heatmap was plotted by plotCorrelation. The samples are as follows: non-treated wild-type (WT), non-treated UGI-expressing (NT_UGI), 5FdUR treated UGI-expressing (5FdUR_UGI), RTX treated UGI-expressing HCT116 cells (RTX_UGI), non-treated wild-type K562 cells (K562). Replicates are indicated as rep1 and rep2.

#### 2.3. Genome-wide analysis of uracil-DNA pattern comparing to ChIP-seq data and other genomic features

To find colocalizing binding factors and other genomic features, first a HCT116 specific or relevant set of data were collected. On the one hand, from Cistrome database (http://cistrome.org/db/#/, (17), data reflect the state of 17 July 2019), overall 542 ChIP-seq data made in HCT116 for transcription factors or histone markers were downloaded as interval (bed) files. Although, these data are still heterogeneous regarding quality and the applied treatments, it is definitely more reasonable than searching in the whole Cistrome database without any restriction for cell types. Only those data were comprised that contained more than 400 intervals (471 files remained). It is also has to be noted that no controls (such as input samples in our case) are considered in the evaluation pipeline of Cistrome. To strengthen this dataset, further HCT116-specific ChIP-seq data (36 bed files) were downloaded from ENCODE (https://www.encodeproject.org/search/?type=Experiment&status=released&replicates.library.biosample.donor.organism.scientific_name=Homo+sapiens&assembly=GRCh38&biosample_ontology.classification=cell+line&biosample_ontology.term_name=HCT116 (18)), where evaluation pipeline (https://www.encodeproject.org/pages/pipelines/) considers controls and includes many quality measures, and the resulted “replicated peaks” reflect only the consensus peaks of replicates and pseudo-replicates. Further 27 bed files were downloaded from the Ensembl database ((19), release 97, July 2019, ftp://ftp.ensembl.org/pub/release-97/regulation/homo_sapiens/Peaks/HCT116/). Moreover, for colorectal tissues, comprehensive epigenomic data focusing on five core histone marks (H3K4me3, H3K4me1, H3K27me3, H3K9me3, and H3K36me3) were constructed by Roadmap Epigenomics (https://egg2.wustl.edu/roadmap/web_portal/processed_data.html, (20)). From these data, overall 40 bed files (broad- and gappedPeaks) corresponding to E075 Colonic Mucosa (7 experiments); E076: Colon Smooth Muscle (7 experiments); and E106: Sigmoid Colon (6 experiments)) were also integrated to our dataset for comparison with U-DNA-Seq data. These files were originally aligned to the hg19 reference genome, therefore liftOver (https://github.com/ucscGenomeBrowser/kent) was applied to convert the coordinates to hg38 as follows.

```
$ liftOver NAME_of_DB_intervals.bed hg19ToHg38.over.chain NAME_ofDB_intervals_hg38.bed
unMapped
```

*# The file hg19ToHg38.over.chain can be downloaded from the UCSC (http://hgdownload.soe.ucsc.edu/goldenPath/hg19/liftOver/hg19ToHg38.over.chain.gz).*

Finally, to address the published centromeric localization of uracil (15), ChIP-seq data on CENPA in HuRef cells were also downloaded (GSM1105684, bw file (21)). Note that CENPA data are not available for HCT116 cells. In this study, reads were aligned to human reference genome hg19, and enrichment was given in bedgraph format containing only the alpha satellite segments. Following the paper (21), those data bins were selected that showed at least two-fold enrichment, and then bed files were generated (using the same procedure that we used for derivation of region interval files from the log2 coverage tracks). Then liftOver was applied to convert hg19 coordinates to hg38 (GSM1105684_2fold_enriched_merged_hg38.bed). Data from the same experiment appear also in the Cistrome database (40153), where data were simply realigned to hg38 reference genome, although the original paper reported much more careful procedure on mapping reads in the highly repetitive centromeres (21). Other CENPA data in Cistrome database represent results on ectopically expressed CENPA, outside of the centromeres.

After neglecting files that contain less than 400 intervals, overall 576 bed files remained in the dataset. Because of the limitation of the applied GIGGLE search tool ((6) version 1.0, cf. issue #46 https://github.com/ryanlayer/GIGGLE/issues/46), top 100,000 intervals were selected from those interval (bed) files that contained more than 100,000 intervals (overall 50 bed files were cut in this way).

Then GIGGLE search was performed on this set of relevant and good quality data with all the U-DNA-Seq samples corresponding to HCT116 cells. A digestion of the results is shown in Figure 4A, while the whole table with the combo scores can be found in Appendix 2. The GIGGLE search was done as follows:

*# The database interval files as well as the query interval files have to be sorted and gzipped using a script belongs to the GIGGLE package using also bgzip tool that has to be installed in advance (https://github.com/samtools/htslib/releases/, htslib-1.9.tar.bz2).*

```
$  {PATH}/GIGGLE/scripts/sort_bed "{PATH}/*.bed" bed_sorted 4
$  {PATH}/GIGGLE/scripts/sort_bed "{PATH}/*.region.bed" own_bed_sorted 4
```

*# Indexing the database*

```
$ GIGGLE index -i "bed_sorted/*gz" -o bed_sorted_b -f -s
```

*# Running GIGGLE search in this indexed library*

```
$ GIGGLE search -i bed_sorted_b -q
own_bed_sorted/NAME.filtered_blacklisted.bin100bp.smooth5k.RPGC.log2.0p2.region.bed.gz
-s > NAME.log2.0p2.regions.GIGGLE_results.csv
```

Note: In some cases, GIGGLE search might end in an error message “too many open files”. To solve this problem, a soft limit (-Sn) of the possible open files has to be checked and changed on the linux operating system (it is possible to do up to the hard limit (-Hn)).

*# Checking:*

```
$ ulimit -Sn
$ ulimit -Hn
```

*# Changing:*

```
$ ulimit -Sn 4096
```

We also wanted to measure colocalization with other genomic features such as cytogenetic bands, coding regions vs non-coding, repeat classes, or replication timing. Data were collected from UCSC Table Browser ((22) http://genome.ucsc.edu/cgi-bin/hgTables), as it is labelled in Supplementary Table S4. These tab delimited text files were rearranged to fulfil the minimum requirements of the bed format, and sub-categories were also selected into separated bed files using awk.

Replication timing data specific for HCT116 cells were obtained from Replication Domain database (Int90617792 and Int97243322, https://www2.replicationdomain.com/database.php (16)). From these wiggle files, early, middle and late replicating segments were extracted to bed files as follows:

```
$ awk ' $4 > 2 ' Int90617792.wig > Int90617792_early_RT.bed
$ awk ' $4 < 0 ' Int90617792.wig > Int90617792_neg_RT.wig
$ awk ' $4 > -2 ' Int90617792_neg_RT.wig > Int90617792_late_RT.wig
```

*# Note that awk use ”>” for the absolute value of the negative numbers.*

```
$ awk ' $4 > 0 ' Int90617792.wig > Int90617792_pos_RT.wig
$ awk ' $4 <= 2 ' Int90617792_pos_RT.bed > Int90617792_mid_pos_RT.bed
$ awk ' $4 <= -2 ' Int90617792_neg_RT.bed > Int90617792_mid_neg_RT.bed
$ cat Int90617792_mid_pos_RT.bed Int90617792_mid_neg_RT.bed | sort -k1,1 -k2,2n >
Int90617792_mid_RT_sorted.bed
```

*# From this state, the processing steps are the same, as in case of the log2 track derived bed files involving bedtools merge, bigWigAverageOverBed, sort and awk.*

Additional data were collected from Ensembl database (19): genomic annotations (ftp://ftp.ensembl.org/pub/release-97/gff3/homo_sapiens/Homo_sapiens.GRCh38.97.gff3.gz) and HCT116 specific regulatory features corresponding to transcriptional activity (ftp://ftp.ensembl.org/pub/release-97/regulation/homo_sapiens/RegulatoryFeatureActivity/HCT116/homo_sapiens.GRCh38.HCT116.Regulatory_Build.regulatory_activity.20190329.gff.gz). From this latter file, relevant interval files corresponding to different categories (e. g. promoter, enhancer, etc., cf. Supplementary Table S4) were derived as follows:

```
$ awk '{gsub(/\;/, "\t", $9)} {print "chr"$1 "\t" $4 "\t"$5 "\t"$3 "\t" $9}'
homo_sapiens.GRCh38.HCT116.Regulatory_Build.regulatory_activity.20190329.gff >
homo_sapiens.GRCh38.HCT116.Regulatory_Build.regulatory_activity_separated.20190329.gff.bed
```

```
$ awk '$5=="activity=ACTIVE" {print $1 "\t" $2 "\t"$3 "\t"$4 "\t" $5}'
homo_sapiens.GRCh38.HCT116.Regulatory_Build.regulatory_activity_separated.20190329.gff.bed
> homo_sapiens.GRCh38.HCT116.Regulatory_Build.regulatory_activity_ACTIVE.20190329.gff.bed
```

```
$ awk '$4=="promoter" {print $1 "\t" $2 "\t"$3 "\t"$4 "\t" $5}'
homo_sapiens.GRCh38.HCT116.Regulatory_Build.regulatory_activity_ACTIVE.20190329.gff.bed >
homo_sapiens.GRCh38.HCT116.Regulatory_Build.regulatory_activity_PROMOTER.20190329.gff.bed
```

Data for RNA genes (cf. long non-coding RNAs (lnc_RNA) were similarly derived from the Homo_sapiens.GRCh38.97.gff3 file.

To further access the alpha satellites and the assembled higher order repeat segments (HORs), another interval file corresponding to the publication (23) was also downloaded (https://genome.ucsc.edu/cgi-bin/hgTrackUi?hgsid=771843343_kD7KrH9deXCkpCCcKNTjvq4t3jOi&c=chr1&g=ct_HMMERSF1HORst281_7253).

These features often contain too many and/or too large intervals for which, GIGGLE 1.0 was not proved to be efficient (cf. issue #46 https://github.com/ryanlayer/GIGGLE/issues/46). Therefore, bedtools annotate was applied to count the overlaps between the U-DNA-Seq and the database intervals. The numbers of overlapping bases between each sample and each database interval file were summarized and scores were calculated according to the following formula:

$$\frac{N^{o}\ of\ overlaping\ bases\ *100}{N^{o}\ of\ bases\ in\ sample\ intervals} * \frac{N^{o}\ of\ overlaping\ bases\ *100}{N^{o}\ of\ bases\ in\ feature\ intervals}$$

A systematic selection of the tested features is shown in Figure 4B, while the results of the full analysis are provided in Supplementary Table S4. Bedtools annotate was used as follows:

```
$ bedtools annotate –names {short names for the database interval files} -both -i
NAME.filtered_blacklisted.bin100bp.smooth5k.RPGC.log2.0p2.region.bed -files {list of
database interval files} > bedtools_annot.NAME.bed
```

*# For one representative database interval file, the number of overlapping bases is calculated as follows:*

```
$ awk -v OFS="\t" '{$8=$3-$2}1' bedtools_annot_NAME.bed | awk '{($7=$7*$8)}1' | awk
'{(sum1 += $7)} END {print sum1}' > bedtools_annot_NAME_results.csv
```

| | | WT | NT_UGI | 5FU_UGI | RTX_UGI | |
|---|---|---|---|---|---|---|
| | baseNo in intervals | 3.19E+08 | 3.28E+08 | 5.25E+08 | 5.21E+08 | |
| Protein Gene | 1.72E+09 | 3.18E+02 | 4.43E+02 | 8.51E+02 | 1.28E+03 | UCSC, Table Browser: Genes and Gene Predictions / GENCODE v32 / knownGene |
| INTRON | 1.64E+09 | 3.12E+02 | 4.41E+02 | 7.60E+02 | 1.15E+03 | |
| EXON | 1.46E+08 | 9.37E+00 | 1.18E+01 | 2.03E+02 | 3.02E+02 | |
| CDS | 3.97E+07 | 2.12E+00 | 2.72E+00 | 5.60E+01 | 9.35E+01 | |
| UTR | 1.26E+08 | 8.32E+00 | 1.04E+01 | 1.73E+02 | 2.66E+02 | |
| RNA genes | 1.29E+06 | 1.32E-01 | 1.39E-01 | 1.29E+00 | 1.77E+00 | UCSC Table Browser: Repeats / RepeatMasker / rmsk |
| srpRNA | 2.91E+05 | 1.68E-02 | 2.13E-02 | 3.64E-01 | 7.31E-01 | |
| snRNA | 3.66E+05 | 6.43E-02 | 7.33E-02 | 3.40E-01 | 5.98E-01 | |
| scRNA | 1.40E+05 | 1.15E-02 | 8.90E-03 | 1.60E-01 | 3.58E-01 | |
| tRNA | 1.22E+05 | 8.10E-03 | 8.89E-03 | 2.12E-01 | 1.26E-01 | |
| rRNA | 2.55E+05 | 1.59E-02 | 1.36E-02 | 1.79E-01 | 9.26E-02 | |
| RNA | 1.18E+05 | 2.53E-02 | 2.48E-02 | 6.31E-02 | 5.87E-02 | |
| lincRNA (colon) | 2.01E+08 | 7.81E+01 | 1.03E+02 | 1.23E+02 | 5.78E+01 | UCSC Table Browser: Genes and Gene Predictions / lincRNA RNA-Seq / Colon (lincRNAsCTColon) |
| tRNAs | 4.68E+04 | 5.02E-05 | 1.62E-04 | 2.30E-01 | 1.10E-01 | UCSC Table Browser: Genes and Gene Predictions / tRNA Genes / tRNAs |
| lnc_RNA | 1.08E+09 | 2.61E+02 | 3.61E+02 | 5.16E+02 | 7.12E+02 | Ensembl |
| miRNA | 3.03E+04 | 8.85E-04 | 1.77E-03 | 6.51E-02 | 6.72E-02 | UCSC Table Browser: Expression / miRNA Tissue Atlas / sample1 (miRnaAtlasSample1BarChart) |
| mi_snoRNA | 1.95E+05 | 1.33E-02 | 1.67E-02 | 3.33E-01 | 4.47E-01 | UCSC Table Browser: Genes and Gene Predictions / sno/miRNA / wgRna |
| CpGIsland (unmasked) | 3.36E+07 | 2.34E-01 | 1.99E-01 | 9.60E+01 | 5.04E+01 | UCSC Table Browser: Regulation / Unmasked CpG / cpgIslandExtUnmasked |
| rmsk_LINE | 6.72E+08 | 3.81E+02 | 4.33E+02 | 1.77E+02 | 2.00E+02 | UCSC Table Browser: Repeats / RepeatMasker / rmsk |
| rmsk_SINE | 4.17E+08 | 3.59E+01 | 3.33E+01 | 4.54E+02 | 6.99E+02 | |
| rmsk_LTR | 2.82E+08 | 3.68E+01 | 3.52E+01 | 1.58E+02 | 9.70E+01 | |
| rmsk_Retroposon | 4.54E+06 | 1.74E-02 | 1.20E-02 | 7.06E+00 | 5.45E+00 | |
| rmsk_DNA | 1.07E+08 | 4.28E+01 | 4.07E+01 | 5.66E+01 | 5.55E+01 | |
| rmsk_DNAq | 4.59E+05 | 2.81E-01 | 2.74E-01 | 1.21E-01 | 7.61E-02 | |
| rmsk_Helitron | 3.81E+05 | 2.34E-01 | 2.09E-01 | 1.82E-01 | 1.42E-01 | |
| rmsk_Satellite | 7.89E+07 | 3.02E-01 | 2.64E-01 | 1.60E-01 | 7.28E-02 | |
| rmsk_Simple_repeat | 3.95E+07 | 2.39E+01 | 2.31E+01 | 2.67E+01 | 1.97E+01 | |
| rmsk_Low_complexity | 6.39E+06 | 2.31E+00 | 2.33E+00 | 4.81E+00 | 3.53E+00 | |
| rmsk_Unknown | 7.53E+05 | 3.57E-01 | 3.48E-01 | 2.93E-01 | 1.73E-01 | |
| simpleRepeat | 1.49E+08 | 5.09E+00 | 6.71E+00 | 3.48E+01 | 3.64E+01 | UCSC Table Browser: Repeats / Simple Repeats / simpleRepeat |
| nestedRepeat | 8.89E+08 | 2.21E+02 | 3.43E+02 | 4.26E+02 | 4.99E+02 | UCSC Table Browser: Repeats / Interrupted Rpts / nestedRepeats |
| microsatellites | 1.70E+06 | 6.06E-01 | 8.14E-01 | 1.01E+00 | 9.28E-01 | UCSC Table Browser: Repeats / Microsatellite / microsat |
| centromeres | 5.95E+07 | 1.02E-05 | 8.01E-06 | 0.00E+00 | 0.00E+00 | UCSC Table Browser: Mapping and Sequencing / Centromeres / centromeres |
| Satellite_centro_repeats | 7.42E+07 | 2.22E-04 | 6.25E-05 | 5.89E-03 | 5.27E-04 | UCSC Table Browser: Repeats / RepeatMasker / rmsk |
| HORs | 2.04E+07 | 1.53E-08 | 3.59E-08 | 0.00E+00 | 2.26E-08 | (23) |
| cytoBand_gneg | 1.47E+09 | 9.77E+01 | 1.33E+02 | 1.42E+03 | 1.53E+03 | UCSC Table Browser: Mapping and Sequencing / Chromosome Band / cytoBand |
| cytoBand_gpos25 | 2.14E+08 | 6.60E+00 | 1.07E+01 | 3.05E+02 | 4.21E+02 | |
| cytoBand_gpos50 | 4.10E+08 | 4.62E+01 | 6.52E+01 | 2.12E+02 | 1.78E+02 | |
| cytoBand_gpos75 | 4.11E+08 | 2.14E+02 | 3.41E+02 | 8.41E+01 | 4.09E+01 | |
| cytoBand_gpos100 | 4.96E+08 | 8.21E+02 | 1.12E+03 | 1.86E+01 | 1.00E+01 | |
| early_RT1 | 5.80E+08 | 4.78E+00 | 5.37E+00 | 1.85E+03 | 4.04E+03 | Replication Domain Database |
| early_RT2 | 5.74E+08 | 4.72E+00 | 5.21E+00 | 1.80E+03 | 4.03E+03 | |
| middle_RT1 | 1.02E+09 | 1.19E+02 | 1.65E+02 | 8.64E+02 | 5.10E+01 | |
| middle_RT2 | 1.03E+09 | 1.22E+02 | 1.74E+02 | 8.83E+02 | 5.33E+01 | |
| late_RT1 | 6.19E+08 | 3.65E+02 | 5.23E+02 | 1.50E+00 | 1.08E-04 | |
| late_RT2 | 6.09E+08 | 3.60E+02 | 5.01E+02 | 1.76E+00 | 1.14E-04 | |
| CTCF_bind | 4.86E+07 | 2.95E+00 | 3.70E+00 | 9.90E+01 | 8.88E+01 | Ensembl, regulatory build for HCT116 |
| TF_bind | 3.25E+06 | 2.99E-01 | 3.74E-01 | 4.43E+00 | 3.74E+00 | |
| ENHANCER | 1.25E+06 | 6.71E-02 | 1.05E-01 | 1.41E+00 | 3.98E+00 | |
| PROMOTER | 2.28E+07 | 8.02E-02 | 9.87E-02 | 6.50E+01 | 5.58E+01 | |
| PROMOTER_flank | 2.48E+07 | 4.93E-01 | 6.89E-01 | 3.04E+01 | 7.47E+01 | |
| OPEN | 9.71E+04 | 1.46E-02 | 1.80E-02 | 1.12E-01 | 1.15E-01 | |
| DNAseHS_HCT116 | 2.35E+07 | 5.55E-01 | 7.14E-01 | 4.78E+01 | 4.80E+01 | UCSC Table Browser: Regulation / DNase HS / HCT-116 Pk (wgEncodeRegDnaseUwHct116Peak) |
| RegDnaseCluster | 4.58E+08 | 5.91E-01 | 7.51E-01 | 6.12E+01 | 7.96E+01 | UCSC Table Browser: Regulation / DNase Clusters / wgEncodeRegDnaseClustered |

**Supplementary Table S4. Full collection of genomic features compared to the four samples of U-DNA-Seq.** Genomic features were downloaded from the UCSC (22), the Ensembl (19), and the

Replication Domain (16) databases. Higher order repeat segments (HORs, HMMERSF1HORst281top100k.bed, (23)) were downloaded from UCSC. Scores were calculated according the formula given in the text. Abbreviated features are the following: coding sequences (CDS), untranslated regions (UTR), signal recognition particle RNAs (srpRNA), small nuclear (snRNA), small conditional (scRNA), long non-coding RNA (lncRNA), long intergenic non-coding RNAs found in colon tissues (lincRNA (colon)), micro-RNA (miRNA), micro and small nucleolar (mi_snoRNA) RNAs, short interspersed nuclear elements (SINE) and long interspersed nuclear elements (LINE), long terminal repeat element (LTR), putative DNA repeat elements (DNAq), cytological bands stained by Giemsa (cytoBand_gneg: non-stained, cytoBand_pos25 up to pos100: show increasing staining intensity), early, middle and late replication timing (RT), CCCTC-Binding factor binding sites that might correspond to DNA loops, insulators, chromatin anchoring point and borders between hetero- and euchromatin (CTCF-binding), opened chromatin structure (OPEN), DNase hypersensitive sites (DNaseHS), transcription factor binding sites (TF_binding_site).

Detailed correlation analysis between uracil-DNA enrichment and replication timing was also done using the R script below (cf. Figure 4C and Supplementary Figure S11C).

```
#used packages
library(tidyverse)
library(BSgenome.Hsapiens.NCBI.GRCh38)

#input the replication timing data
rt <- read.table(„RT_data/RT_HCT116_Epithelial cells from colon carcinoma_Int90617792_hg38.bedgraph”)
names(rt) <- c(„Chr”, „Start”, „End”, „RT”)
#list of bigwig files
list.files(path = "readfiltered_bwcomp//", pattern = "*.bw") -> bwlist
strsplit(bwlist, split = "\\.") %>% sapply("[", 1) -> namelist

dir.create("readfiltered_U_vs_RT_plots/")

#list to store the aggregated data
ll <- vector(mode = "list", length = length(bwlist))
names(ll) <- namelist

#the for cycle will match each of the U-DNA-Seq data to RT and make pictures for all one by one
for (b in seq_along(bwlist)) {
  #input bigwig files
  bw1 <- import.bw(paste0("readfiltered_bwcomp/", bwlist[b]))
  GRanges(rt) -> rtgr
  #ensuring that only the common chromosomes and scaffolds will be considered
  bw1 <- keepSeqlevels(x = bw1, value = seqlevels(rtgr), pruning.mode = "coarse")
  #the RT data are given in bins of 5000 bases, while the uracil enrichment was calculated for bins of 100 bases
  #the principle of the comparison is first to define the overlaps between the intervals of the two datasets, then to
average the uracil enrichment scores for the bins of RT data. In this way, the object tt will contain both RT and the
corresponding average of uracil enrichment scores for each bin of 5000 bases.
  olaps <- findOverlaps(subject = rtgr, query = bw1)
  as.data.frame(olaps) %>% mutate(U_score = bw1$score[queryHits]) %>% group_by(subjectHits) %>%
summarize(U_score_mean = mean(U_score))
  %>%
    mutate(RT = rtgr$RT[subjectHits]) -> tt
  #the resulting data frame is copied to the appropriate position of the summarizing list
  ll[[namelist[b]]] <- tt
  #labelling the RT values to have 10 equal-sized groups for the further plotting
  .bincode(tt$RT, breaks = quantile(tt$RT, probs = seq(0, 1, length.out = 21))) -> tt$RT_Cut
```

```
  #plots one by one
  tt %>% ggplot(aes(x = RT, y = U_score_mean)) + stat_density2d(aes(alpha = ..level.., fill = ..level..), geom = "polygon") + scale_fill_viridis_c() + theme_bw() + guides(alpha = FALSE, fill = FALSE) + ggtitle(paste0("Uracil log2 overrepresentation vs. RT for ", namelist[b])) + coord_cartesian(ylim = c(-1.2, 1.2))
  ggsave(paste0(file = paste0("readfiltered_U_vs_RT_plots/", namelist[b], ".chr.U_vs_RT.genome.pdf")), width = 8, height = 5)
  ggplot(tt, aes(x = RT_Cut, y = U_score_mean, group = RT_Cut)) + geom_boxplot() + coord_cartesian(ylim = c(-2, 2))
  ggsave(paste0(file = paste0("readfiltered_U_vs_RT_plots/_", namelist[b], ".chr.U_vs_RT.boxplot.pdf")), width = 8, height = 5)
}

#plots for multiple samples

#labelling the 10 equal sized groups according to the RT data on each elements of the summarizing list.
ll2 <- lapply(ll, function(x) mutate(x, RT_Cut = .bincode(RT, breaks = quantile(RT, probs = seq(0, 1, length.out = 11)))))
do.call(rbind.data.frame, ll2) -> df
#additional information on formatting
df <- mutate(df,
             Sample = gsub("\\.[0-9]+", "", rownames(df)),
             Condition = NA,
             Condition = ifelse(Sample == "K562_IP_vs_son",
                                "K562",
                                ifelse(Sample %in% c("3CG_vs_1CG", "U_IP_cs_son"),
                                       "UGI + 5FdU",
                                       ifelse(Sample %in% c("4CG_vs_2CG", "NT_IP_vs_son"),
                                              "UGI",
                                              ifelse(Sample %in% c("URTX1_IP_vs_son", "URTX2_IP_vs_son"),
                                                     "UGI + RTX",
                                                     "WT")))),
             Condition = factor(Condition, levels = c("K562", "WT", "UGI", "UGI + 5FdU", "UGI + RTX"))) %>%
  filter(!is.na(Condition), Sample != "4CG_vs_2CG", Sample != "U_IP_vs_son", !is.na(RT_Cut)) %>%
  mutate(Group = NA,
         Group = ifelse(Condition %in% c("UGI + RTX", "UGI + 5FdU"),
                        "inhibited",
                        "non-inhibited"))

#plot and save
ggplot(df, aes(x = factor(RT_Cut), y = U_score_mean, fill = Condition)) + geom_boxplot(size = .3, outlier.shape = NA) + theme_classic() + coord_cartesian(ylim = c(-1.5, 1.2), clip = "off") + theme(axis.text.y = element_text(size = 12, face = "bold"), axis.title = element_text(size = 16), axis.text.x = element_blank(), axis.ticks.x = element_blank(), axis.line = element_line(size = 1)) + ylab(bquote(log[2] * "(Uracil overrepresentation)")) + xlab("")  + scale_y_continuous(breaks = round(seq(-1.6, 1.2, by = .4), digits = 1)) + theme(legend.text = element_text(size = 12), legend.title = element_text(size = 12)) -> g1
ggsave(filename = "Uracil_overrepresentation.pdf", width = 8, height = 5, plot = g1)
```

#### 3. Characterization of the new uracil-sensor construct

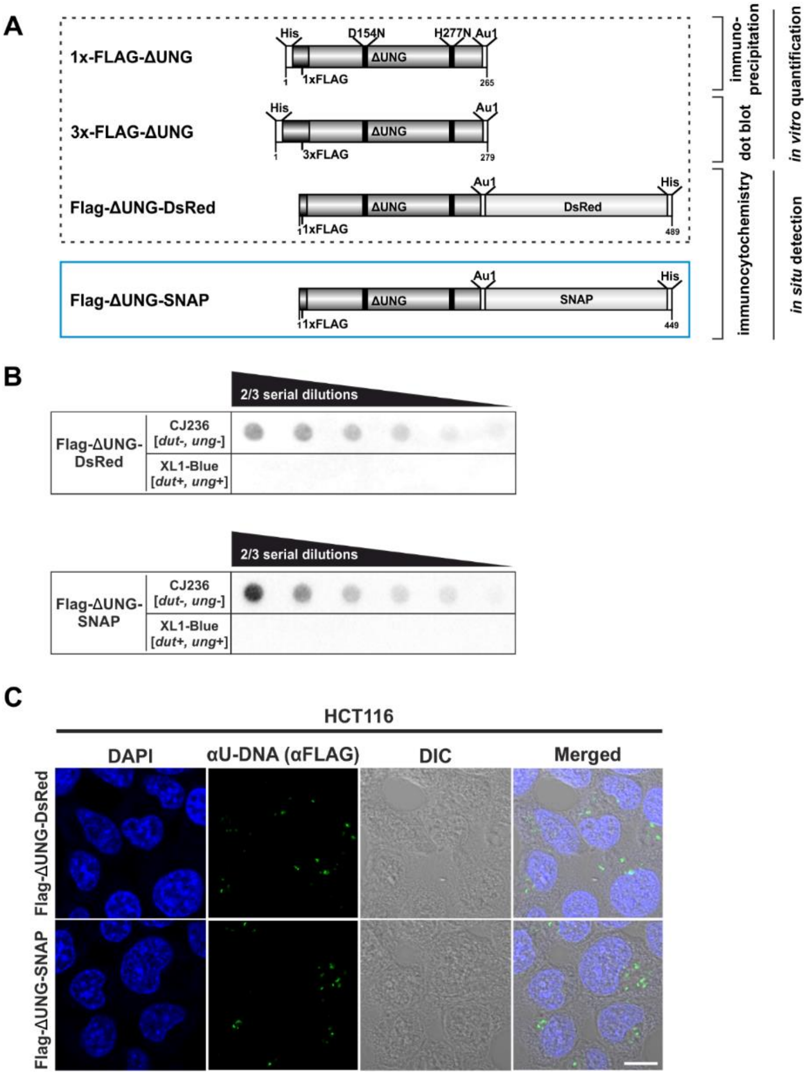

**Supplementary Figure S8. Design and validation of the new FLAG-ΔUNG-SNAP uracil-sensor. (A)** Scheme of the used constructs in this study, with their applications highlighted. The catalytically inactive (D154N and H277N, mutated sites indicated with black lines) and truncated ΔUNG (the N-terminal 84 residues, responsible for the binding to RPA and PCNA, were also removed) was created from human UNG2. The ΔUNG uracil-recognizing core was fused to different epitope tags: His-tag for affinity purification, 1x/3xFLAG and Au1 for antibody-based detection, DsRed-monomer or SNAP-tag for direct fluorescent detection. Our newly developed FLAG-ΔUNG-SNAP sensor construct is indicated with blue frame, separately from the constructs characterized previously (1). **(B)** Dot blot assay was used to compare uracil binding capability of FLAG-ΔUNG-DsRed and FLAG-ΔUNG-SNAP sensor constructs. CJ236 [*dut−, ung−*] (positive control) and XL1-Blue [*dut+, ung+*] (negative control) *E. coli* genomic DNA samples (8 ng) were measured in two-third dilution series. **(C)** Comparison of the FLAG-ΔUNG-DsRed and the FLAG-ΔUNG-SNAP constructs in detecting uracil-rich plasmid DNA aggregates in HCT116 in an immunocytochemistry assay. The sensors were visualized through the FLAG epitope tag. DAPI was used to counterstain DNA. Scale bar represents 10 µm.

### 4. *In situ* detection of the cellular U-DNA content by STED microscopy

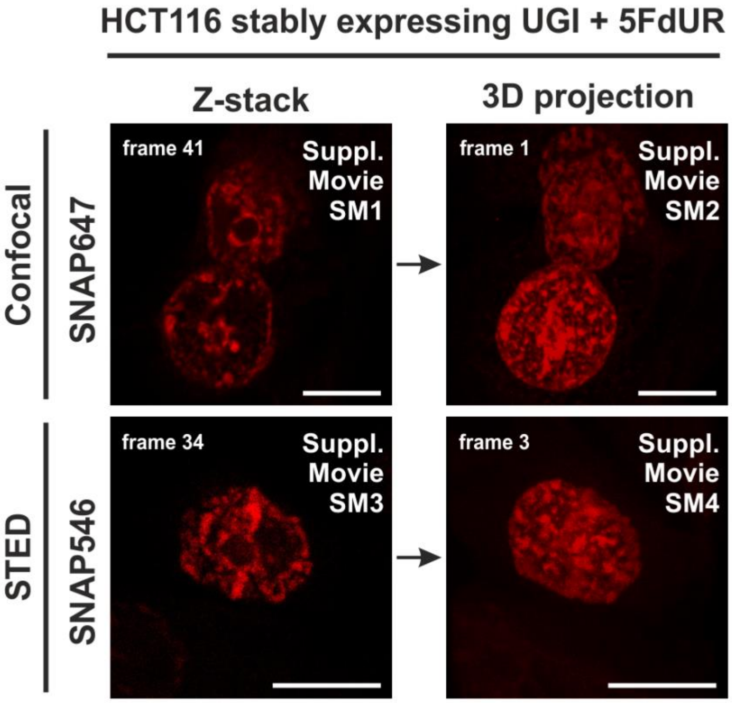

**Supplementary Figure S9. Representative image for Supplementary Movies SM1-SM4.** Genomic uracil levels of HCT116 cells were elevated with 5FdUR treatment and UGI expression. Videos show the following: confocal Z-stack series of 3 cells (Supplementary Movie SM1), 3D projection of this confocal series (Supplementary Movie SM2), Z-stack series of 1 cell acquired by STED microscopy (Supplementary Movie SM3) and 3D projection of this series acquired by STED microscopy (Supplementary Movie SM4). The related movies are indicated at the top right corner of the images. Scale bar represents 10 µm.

**Supplementary Movie SM1. Confocal Z-stack series of 3 cells showing uracil distribution.** Figure S9 serves as an overall representative image for Supplementary Movies.

**Supplementary Movie SM2. 3D projection of Supplementary Movie SM1.** Figure S9 serves as an overall representative image for Supplementary Movies.

**Supplementary Movie SM3. Z-stack series of 1 cell acquired by STED showing uracil distribution.** Figure S9 serves as an overall representative image for Supplementary Movies.

**Supplementary Movie SM4. 3D projection of Supplementary Movie SM3.** Figure S9 serves as an overall representative image for Supplementary Movies.

### 5. Re-analysis and reinterpretation of dU-seq data published in (15)

Shu and co-workers recently published the dU-seq method and reported centromeric location of genomic uracil in non-treated cells (15). In this study, surprisingly, a short-read sequencing technology was used to map centromeric localization of a genomic feature. Centromeres are known to be highly repetitive, poorly mappable regions of the human genome, even if centromeric model sequences (24) are implemented into the hg38 assembly (25). Therefore, blacklisting and mappability filters are highly recommended to be used, especially if the analysis is focusing on such critical regions, like in this case.

The dU-seq data analysis pipeline, as it was published in the *Shu et al*, included the following pre-processing steps: 1) pre-alignment of the 150 bp paired-end sequencing data to the spike-in sequences applied in their experiments; 2) trimming the adaptors and the low quality segments; 3) alignment of the remaining reads to reference human genome hg38 (it is not clear, to which set exactly) using bowtie2. There was no mention either about 1) deduplication of the data, or 2) filtering for uniquely mapped reads, or 3) applying recommended blacklists. However, in widely accepted ChIP-seq pipelines, only uniquely mapped reads are considered as valid information (26).

The detection of uracil enrichment within these aligned reads was done by peak calling using MACS2, separately for the uracil "pull-down" and the "control" samples. It was not mentioned if the corresponding input was included as a control in the peak calling process, or not. It is also not clear what options and parameters were used in their MACS2 runs (e. g. broad, no-model or broad-cutoff). Then they subtracted the peaks detected in the "control" from the ones called in the "pulled-down" samples. We claim that this approach is clearly suboptimal considering the lower descriptive value and the lower reproducibility of peak calling for the uracil-DNA distribution as shown in Supplementary Figure S5 and Supplementary Table S3. To judge the reproducibility of peak calling in dU-seq data, is also not trivial because they did not uploaded all the peak data for their replicates (GSE99011). We calculated the Jaccard indices for their uploaded non-treated HEK293 (2 replicates) and UNG-knock-out HEK293 (4 replicates) data, and found really low values (0.063 for the 2 replicates, and 0.030, 0.059, 0.075, 0.029, 0.029, 0.055 in the pairwise comparison of the 4 replicates). Jaccard indices between the individual replicates and the united data of the non-treated HEK293 sample were 0.134 and 0.092. These values are even lower than in case of similar peak calling on our U-DNA-Seq data (cf. Supplementary Table 3B).

The correlation with the CENPA bound genomic regions, published as a key point in their paper, is also questionable. Raw reads of CENPA ChIP-seq data (GSE45497) were downloaded and mapped to hg38. Details are not provided in the paper, however, they most probably used the same procedure as in case of the dU-seq data, namely, aligning reads without blacklisting and mapping quality filters, followed by peak calling. Moreover, the CENPA data were originally aligned to hg19 reference genome using a more careful algorithm filtering out potential artefacts (21). It would have been recommended to either follow their approach or simply lift over the original alignment from hg19 to hg38.

Unfortunately, *Shu et al* did not upload the aligned reads (bam files) to the GEO database, and the peak calling results for the replicates were also incompletely uploaded. The uploaded peaks were not always correlating with the ones they published on the manuscript's figures (cf. Fig. 2a, where they show a K562 peak on chromosome 21 that is actually not present in their uploaded peak data, and another peak on chromosome 16 that corresponds to aligned reads characterized with MAPQ=0, at least in our alignment. Given missing data uploads, it is challenging to reproduce their results.

Still, we were curious, whether their dU-seq data itself (not the interpretation of that) correlates with our U-DNA-Seq data, or not. Therefore, we used our novel analysis pipeline described in the present study, to process and re-analyse raw data from (15), notably the following data:

K562 „input": SRR5572773/GSM2630035 and SRR5572774/GSM2630036
K562 „control": SRR5572775/GSM2630037 and SRR5572776/GSM2630038
K562 „PD": SRR5572777/GSM2630039 and SRR5572778/GSM2630040
HEK293T UNGKO 5FU „input": SRR5998406/GSM2769605 and SRR5998407/GSM2769606
HEK293T UNGKO 5FU „control": SRR5998408/GSM2769607 and SRR5998409/GSM2769608
HEK293T UNGKO 5FU „PD": SRR5998410/GSM2769609 and SRR6026694/GSM2769610

We remapped the fastq reads to the human reference genome hg38 (GRCh38.d1.vd1.fa, GDC reference, https://gdc.cancer.gov/about-data/data-harmonization-and-generation/gdc-reference-files) using BWA, filtered out ambiguously mapped reads, and applied a blacklist created as in case of our data (cf. Supplementary Figure S3). The statistics of this pre-processing are summarized in Supplementary Table S5. The relative amount of the spike-in sequences is significantly higher in mock pull-down („control") and the non-treated K562 pull-down samples, where the sample DNA itself was not labelled or was present at extremely low level, respectively. The overall depth of sequencing was lower in these cases, which decrease the reliability or reproducibility of further data analysis.

| sample | replicates | number of raw reads | number of mapped* reads | Unmapped* reads | | uniquely mapped reads | | uniquely mapped reads after blacklisting | |
|---|---|---|---|---|---|---|---|---|---|
| | | | | number | % | number | % | number | % |
| K562 input | SRR5572773 | 95,922,009 | 90,663,272 | 5,258,737 | 5.48 | 84,045,278 | 87.62 | 82,192,353 | 85.69 |
| | SRR5572774 | 95,030,662 | 89,816,930 | 5,213,732 | 5.49 | 83,306,810 | 87.66 | 81,457,577 | 85.72 |
| K562 Control | SRR5572775 | 76,394,870 | 46,189,867 | 30,205,003 | 39.54 | 41,385,042 | 54.17 | 40,405,241 | 52.89 |
| | SRR5572776 | 78,962,287 | 41,053,994 | 37,908,293 | 48.01 | 36,260,497 | 45.92 | 35,393,512 | 44.82 |
| K562 PD | SRR5572777 | 87,466,276 | 54,113,837 | 33,352,439 | 38.13 | 48,446,026 | 55.39 | 47,324,075 | 54.11 |
| | SRR5572778 | 82,499,155 | 52,929,849 | 29,569,306 | 35.84 | 47,555,693 | 57.64 | 46,466,915 | 56.32 |
| 5FdUR_UNGKO HEK293 input | SRR5998406 | 125,631,380 | 108,320,783 | 17,310,597 | 13.78 | 101,027,428 | 80.42 | 99,012,730 | 78.81 |
| | SRR5998407 | 70,349,101 | 61,039,638 | 9,309,463 | 13.23 | 56,970,384 | 80.98 | 55,807,073 | 79.33 |
| 5FdUR_UNGKO HEK293 Control | SRR5998408 | 113,654,134 | 62,679,292 | 50,974,842 | 44.85 | 55,333,969 | 48.69 | 54,129,569 | 47.63 |
| | SRR5998409 | 129,196,940 | 58,003,222 | 71,193,718 | 55.10 | 49,706,846 | 38.47 | 48,600,418 | 37.62 |
| 5FdUR_UNGKO HEK293 PD | SRR5998410 | 80,035,762 | 67,939,558 | 12,096,204 | 15.11 | 63,453,866 | 79.28 | 62,184,497 | 77.70 |
| | SRR6026694 | 66,242,483 | 56,303,837 | 9,938,646 | 15.00 | 52,653,804 | 79.49 | 51,598,007 | 77.89 |
| 5FdUR_UGI input | 5FdUR1_son | 128,706,895 | 128,669,770 | 37,125 | 0.03 | 122,476,766 | 95.16 | 118,558,597 | 92.12 |
| | 5FdUR1_son | 201,926,203 | 201,560,665 | 365,538 | 0.18 | 193,086,643 | 95.62 | 184,756,297 | 91.50 |
| 5FdUR_UGI enriched | 5FdUR1_IP | 150,596,242 | 150,522,522 | 73,720 | 0.05 | 144,554,269 | 95.99 | 141,582,874 | 94.01 |
| | 5FdUR2_IP | 138,651,760 | 138,410,833 | 240,927 | 0.17 | 133,200,761 | 96.07 | 128,584,894 | 92.74 |
| K562 input | K562_son | 106,137,622 | 105,875,437 | 262,185 | 0.25 | 100,326,105 | 94.52 | 97,504,876 | 91.87 |
| K562 enriched | K562_IP | 109,490,393 | 109,306,854 | 183,539 | 0.17 | 105,310,296 | 96.18 | 102,117,055 | 93.27 |

**Supplementary Table S5. Statistics on pre-processing of dU-seq data.** Samples from the study of *Shu et al* (15) (non-treated K562 (K562), and 5FdUR treated UNG-knock-out HEK293 cells (5FdUR_UNGKO HEK293)) are compared to our samples (5FdUR treated UGI-expressing HCT116 cells (5FdUR_UGI), and non-treated wild-type K562 cells (K562), which are listed here again for easier comparison (grey, see also in Supplementary Table S2). In case of dU-seq samples, inputs are genomic DNA fragmented and treated according to the dU-seq protocol, also containing additional spike-in sequences; controls are pulled-down in a mock experiment excluding UNG treatment; while PD means the pull-down samples according to the dU-seq protocol. Number of raw reads means read number before starting alignment (the sum of the mapped and unmapped reads). Uniquely mapped read means that MAPQ is not zero. The dU-seq and the U-DNA-Seq samples markedly differ in the ratio of mapped (*) and unmapped (*) reads due to the spike-in DNA applied in dU-seq only.

dU-seq and U-DNA-Seq were performed in completely independent laboratories, even on different continents; applying different conditions; in case of drug treated samples different cell lines; and obviously different experimental protocols. Still, the resulting log2 ratio tracks are in surprisingly good correlation, if we use our robust analysis pipeline. This is demonstrated in Supplementary Figure S10 showing an IGV view, the Pearson correlation analysis, and the histograms of uracil enrichment signal distribution, following the scheme of Figure 3 for better comparison. The clear difference between treated and non-treated samples is also obvious. Furthermore, we have demonstrated that the centromeric peaks published in *Shu et al* (15) localize in blacklisted area (Supplementary Figure S11A). However, the re-analysed dU-seq data could confirm our interpretation on genomic uracil distribution in both non-treated and treated cells, using the herein developed robust analysis pipeline.

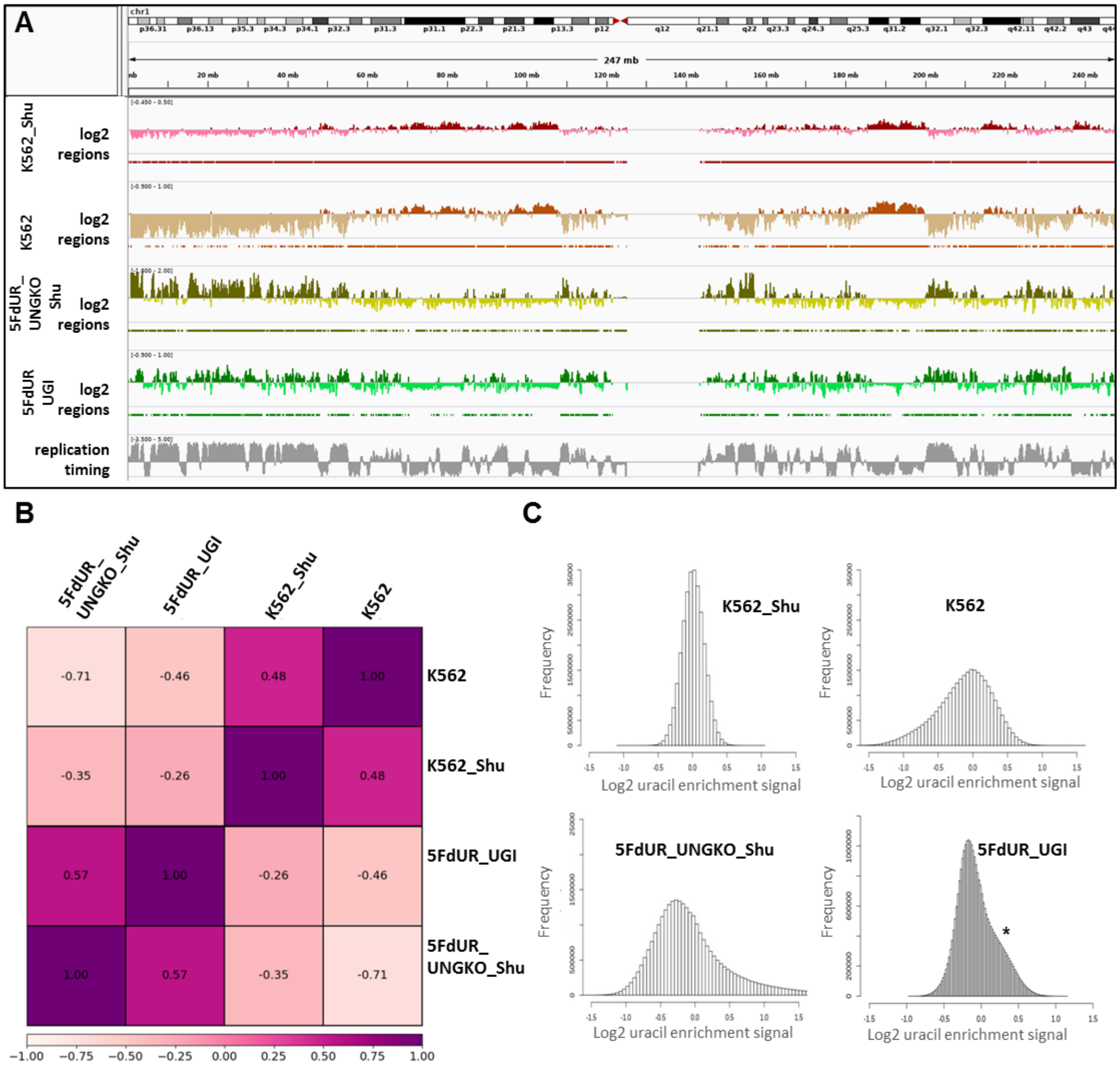

**Supplementary Figure S10. Re-analysis of the published dU-seq data (15) reveals that corresponding samples from dU-seq and U-DNA-Seq show similar patterns of uracil distribution. (A)** IGV view of dU-seq data (non-treated K562 cells (K562_Shu, wine track), and 5FdUR treated UNG-knock-out HEK293 cells (5FdUR_UNGKO_Shu, olive track)), compared to our own U-DNA-Seq data (non-treated K562 cells (K562, brown track), and 5FdUR treated UNG inhibited HCT116 cells (5FdUR_UGI, green track)) on chromosome 1. Log2 ratio tracks and the derived regions of uracil enrichment are also indicated. The bottom track shows replication timing data (grey) for HCT116 downloaded from Replication Domain database (16). **(B)** Pearson correlation among dU-seq and U-DNA-Seq log2 ratio tracks calculated from merged replicates. The treated and non-treated samples are well separated again. Pearson correlation between corresponding dU-seq and U-DNA-Seq samples are unexpectedly high, especially considering the cell line difference in case of drug-treated cells, and the overall low signal intensity in case of non-treated K562. **(C)** Log2 ratio signal distribution of dU-seq and U-DNA-Seq data. The non-treated K562 samples result in a normal like distribution of uracil enrichment signals, while in case of 5FdUR treated cells, these distributions show asymmetry: either a clear shoulder (asterisk), or a more elongated tail towards increased signals in both U-DNA-Seq and dU-seq data, respectively.

A

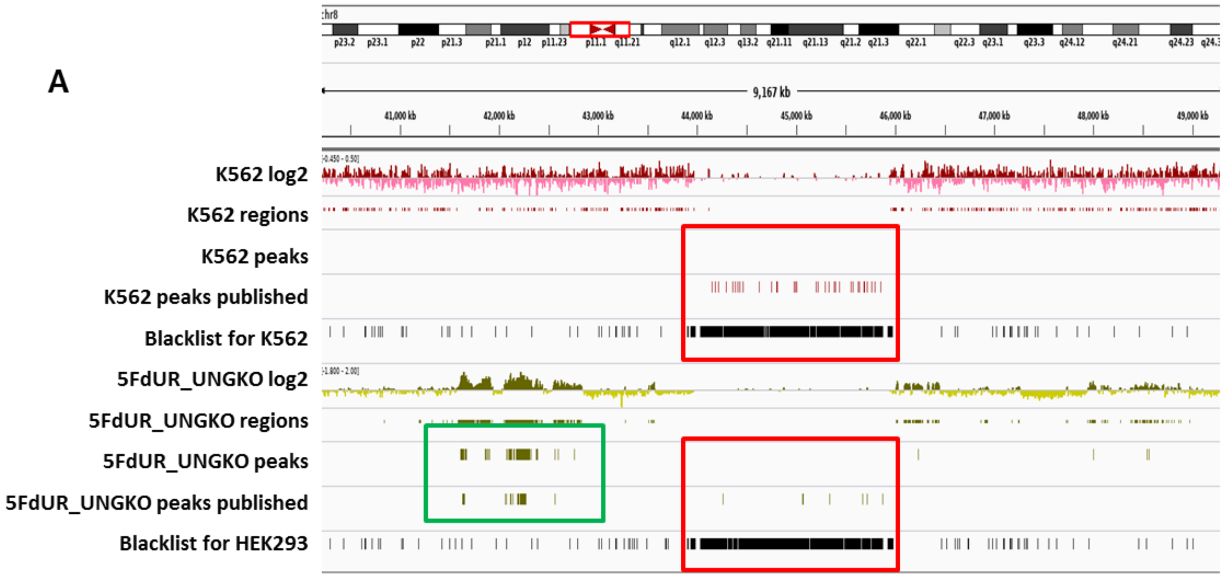

B

| | | K562 Shu | K562 | 5FdUR_UNGKO_Shu | 5FdUR_UGI |
|---|---|---|---|---|---|
| | baseNo of interval files | 3.27E+08 | 3.24E+08 | 5.36E+08 | 5.25E+08 |
| Protein Genes | 1.72E+09 | 526.58 | 475.82 | 962.80 | 850.7 |
| INTRON | 1.64E+09 | 519.28 | 470.83 | 832.89 | 759.9 |
| EXON | 1.46E+08 | 19.57 | 14.22 | 344.47 | 202.5 |
| CDS | 3.97E+07 | 4.23 | 3.32 | 110.72 | 56.0 |
| RNA Genes | 1.29E+06 | 0.25 | 0.14 | 1.56 | 1.29 |
| CpG Island | 2.42E+07 | 2.22 | 0.30 | 251.82 | 96.0 |
| SINE | 4.17E+08 | 87.65 | 39.56 | 578.14 | 454.4 |
| LTR | 2.82E+08 | 86.16 | 38.92 | 155.15 | 157.8 |
| LINE | 6.72E+08 | 344.42 | 390.95 | 112.41 | 177.4 |
| satellite | 7.89E+07 | 1.53 | 0.28 | 1.01 | 0.16 |
| cytoBand_gneg | 1.47E+09 | 257.54 | 149.45 | 1561.93 | 1416.7 |
| cytoBand_gpos25 | 2.14E+08 | 36.44 | 13.47 | 394.05 | 305.3 |
| cytoBand_gpos50 | 4.10E+08 | 133.33 | 76.61 | 179.72 | 212.3 |
| cytoBand_gpos75 | 4.11E+08 | 280.67 | 327.63 | 38.16 | 84.1 |
| cytoBand_gpos100 | 4.96E+08 | 490.80 | 985.30 | 14.83 | 18.6 |
| DNAseHS | 2.35E+07 | 3.11 | 1.12 | ND | 47.8 |

C

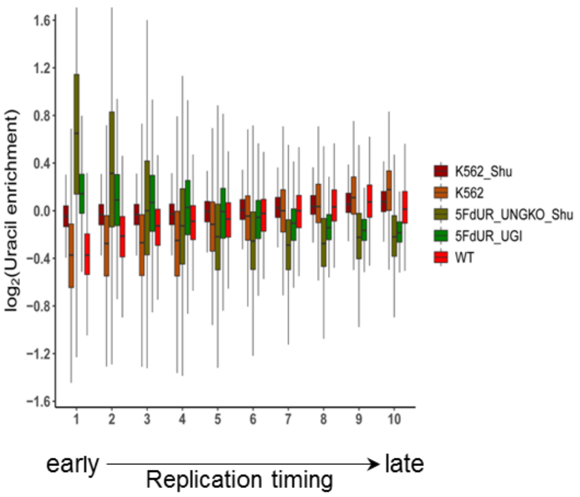

**Supplementary Figure S11. Reinterpretation of dU-seq data. (A)** Comparison of the re-analysed and the published dU-seq data in a representative IGV view of chromosome 8 (cf. also Fig. 3a in *Shu et al* (15)). All centromeric peaks in K562 published for chromosome 8 in *Shu et al* are found in blacklisted regions (red box). Overall, 75% of their published peaks in K562 are overlapping with blacklisted regions determined by our protocol (cf. Supplementary Figure S3). Accordingly, no peaks were called in the presented region during the re-analysis of the sequencing data (red box). Similarly, in drug-treated samples, published centromeric peaks were not reproducible (red box), while other peaks outside of the centromeres were similar in the published and the re-analysed data (green box). **(B)** dU-seq data shows similar correlation to genomic features as compared to the corresponding U-DNA-Seq data. Similarity were measured by bedtools annotate tool and the scores were calculated in the same way as it was in Figure 4B. For each sample, cell type (HCT116 or K562) specific DNase hypersensitive site data were used. For treated HEK293 cells similarity was not addressed (grey). **(C)** Correlation between uracil distribution and replication timing was confirmed by dU-seq data as well, although this correlation is weaker than the U-DNA-Seq results (cf. Figure 4C).

### 6. Appendix

**Appendix 1. Accompanying Supplementary Figure S6. IGV views of log2 ratio and regions of uracil enrichment on all the chromosomes.**

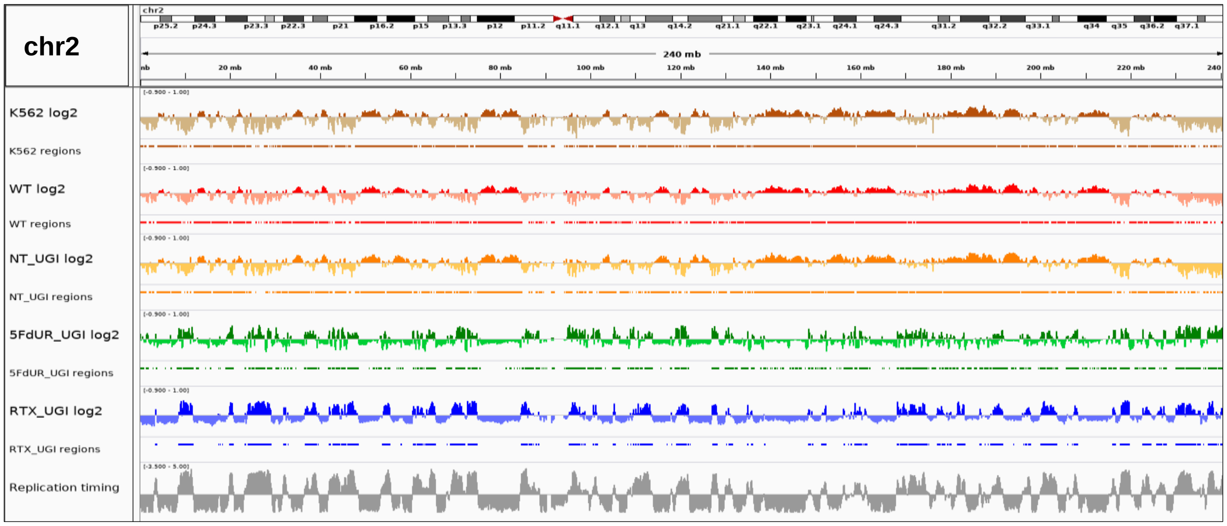

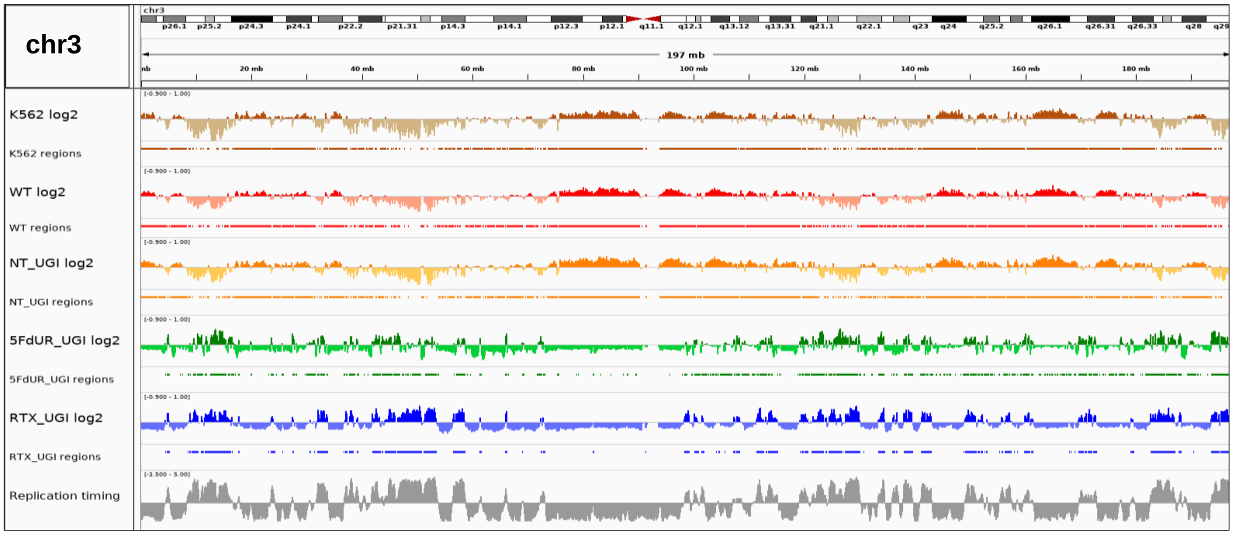

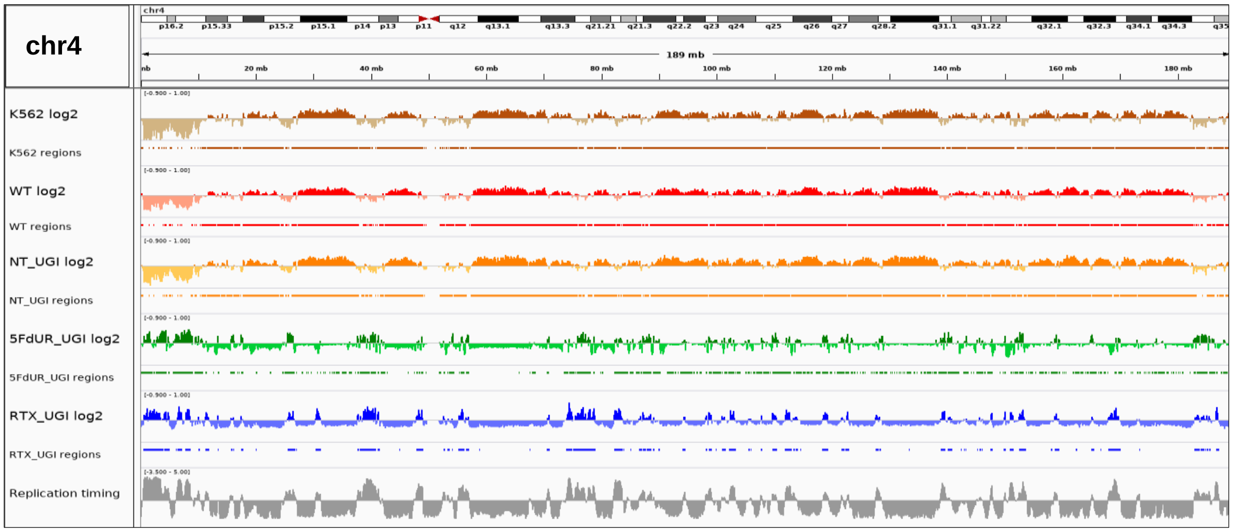
chr4
189 mb
K562 log2
K562 regions
WT log2
WT regions
NT_UGI log2
NT_UGI regions
5FdUR_UGI log2
5FdUR_UGI regions
RTX_UGI log2
RTX_UGI regions
Replication timing

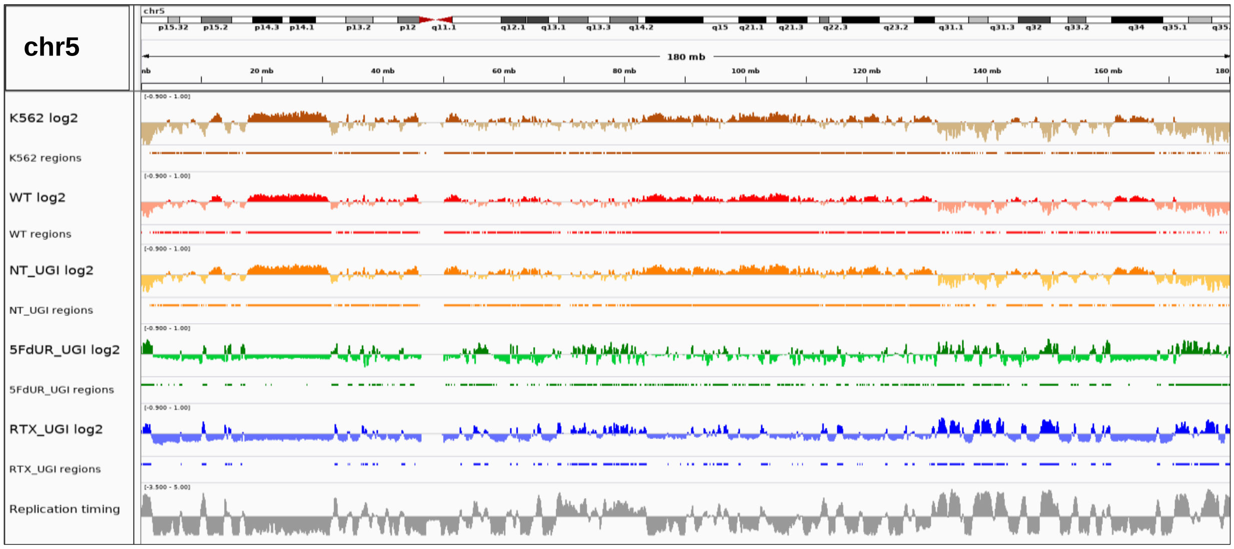
chr5
180 mb
K562 log2
K562 regions
WT log2
WT regions
NT_UGI log2
NT_UGI regions
5FdUR_UGI log2
5FdUR_UGI regions
RTX_UGI log2
RTX_UGI regions
Replication timing

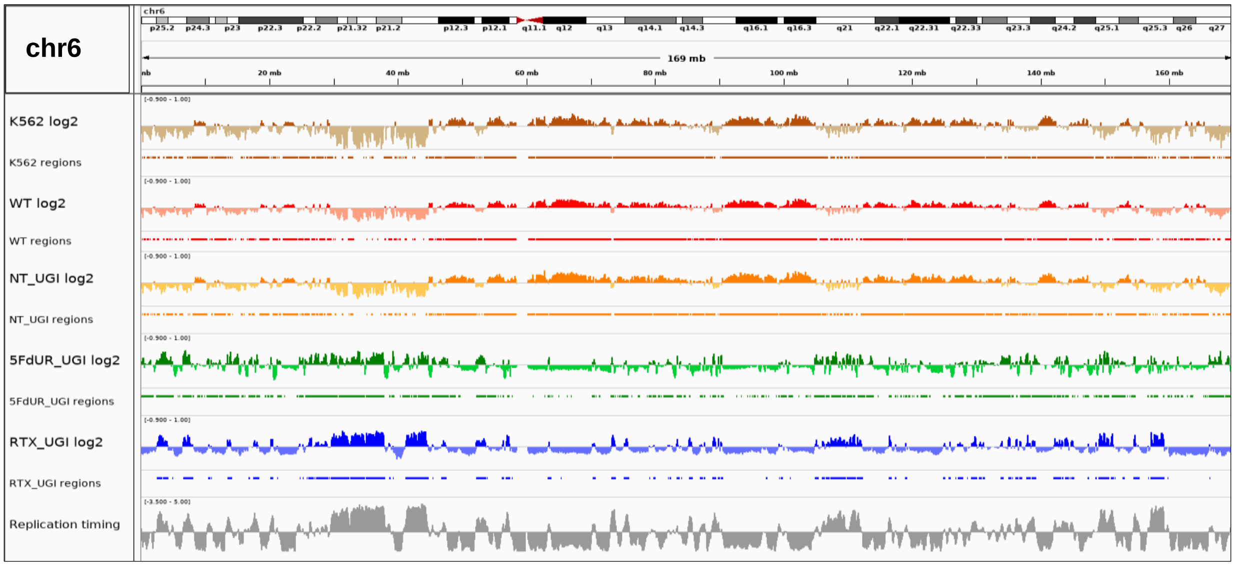
chr6
169 mb
K562 log2
K562 regions
WT log2
WT regions
NT_UGI log2
NT_UGI regions
5FdUR_UGI log2
5FdUR_UGI regions
RTX_UGI log2
RTX_UGI regions
Replication timing

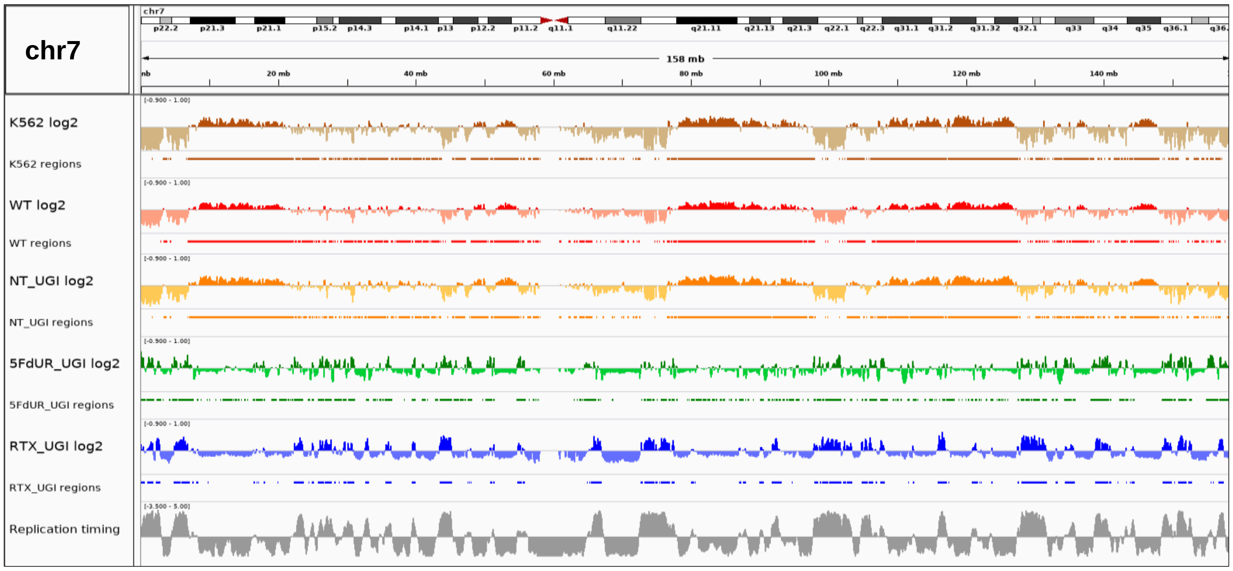
chr7
158 mb
K562 log2
K562 regions
WT log2
WT regions
NT_UGI log2
NT_UGI regions
5FdUR_UGI log2
5FdUR_UGI regions
RTX_UGI log2
RTX_UGI regions
Replication timing

919

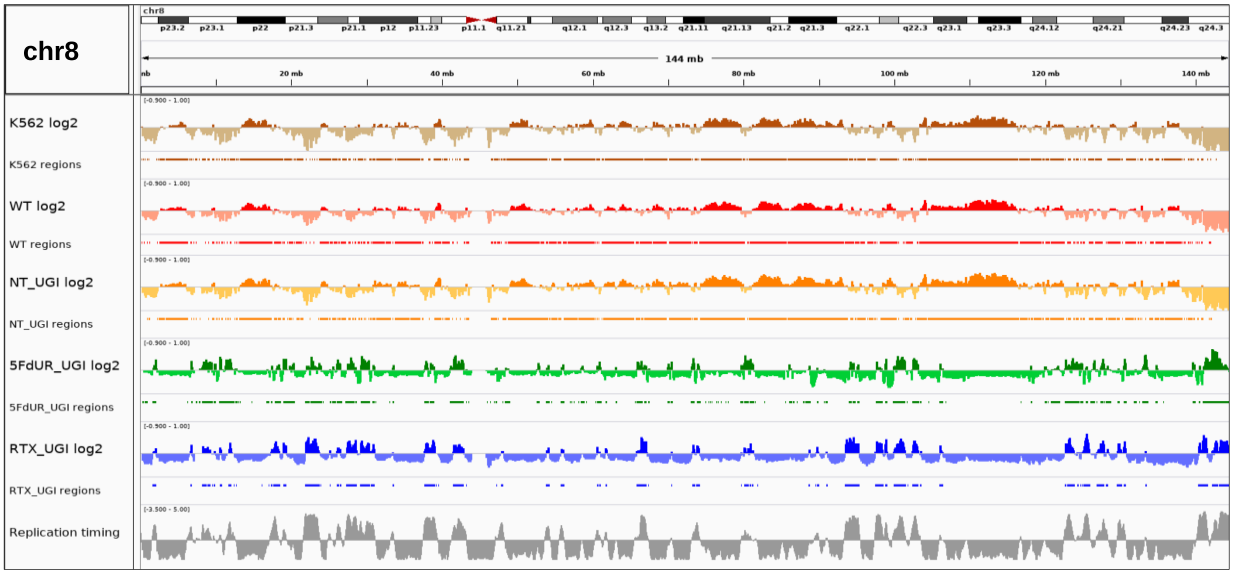
chr8
144 mb
K562 log2
K562 regions
WT log2
WT regions
NT_UGI log2
NT_UGI regions
5FdUR_UGI log2
5FdUR_UGI regions
RTX_UGI log2
RTX_UGI regions
Replication timing

920

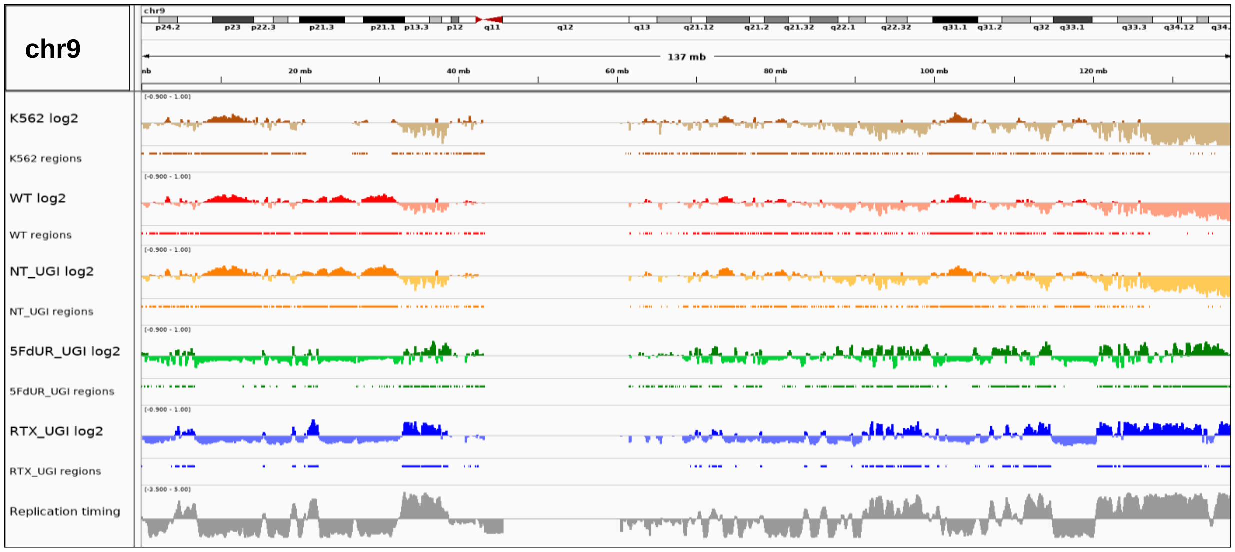
chr9
137 mb
K562 log2
K562 regions
WT log2
WT regions
NT_UGI log2
NT_UGI regions
5FdUR_UGI log2
5FdUR_UGI regions
RTX_UGI log2
RTX_UGI regions
Replication timing

921

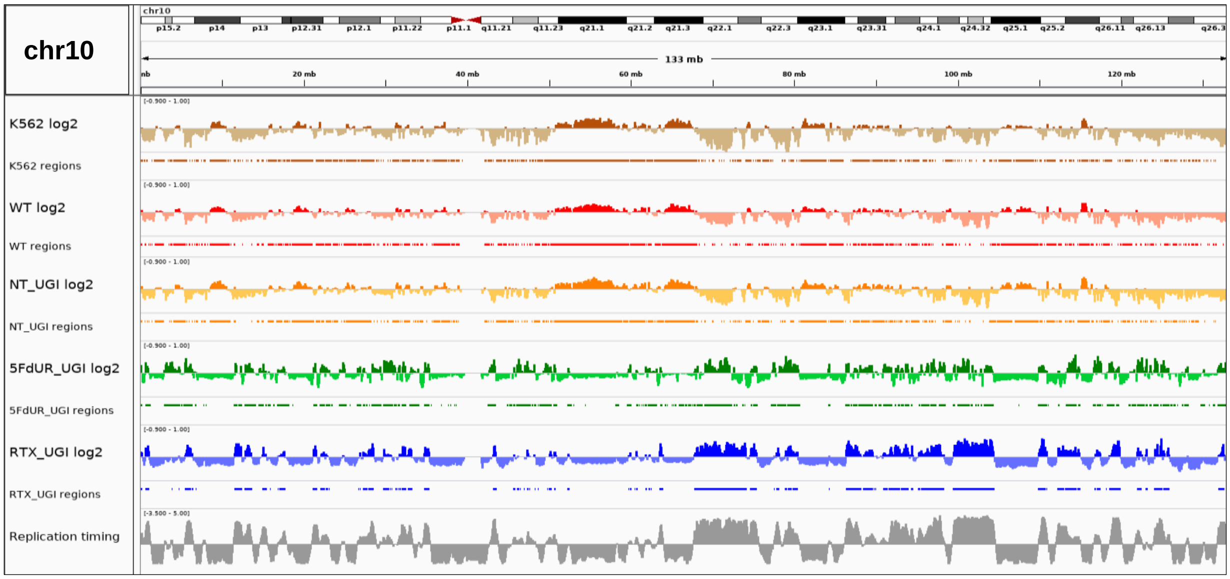
chr10
K562 log2
K562 regions
WT log2
WT regions
NT_UGI log2
NT_UGI regions
5FdUR_UGI log2
5FdUR_UGI regions
RTX_UGI log2
RTX_UGI regions
Replication timing

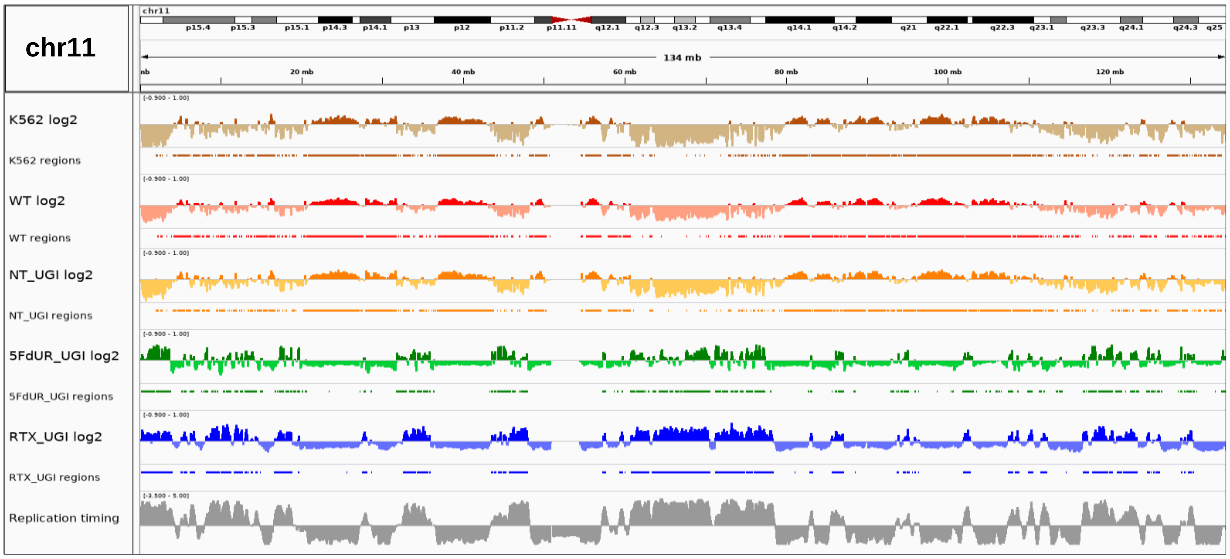
chr11
K562 log2
K562 regions
WT log2
WT regions
NT_UGI log2
NT_UGI regions
5FdUR_UGI log2
5FdUR_UGI regions
RTX_UGI log2
RTX_UGI regions
Replication timing

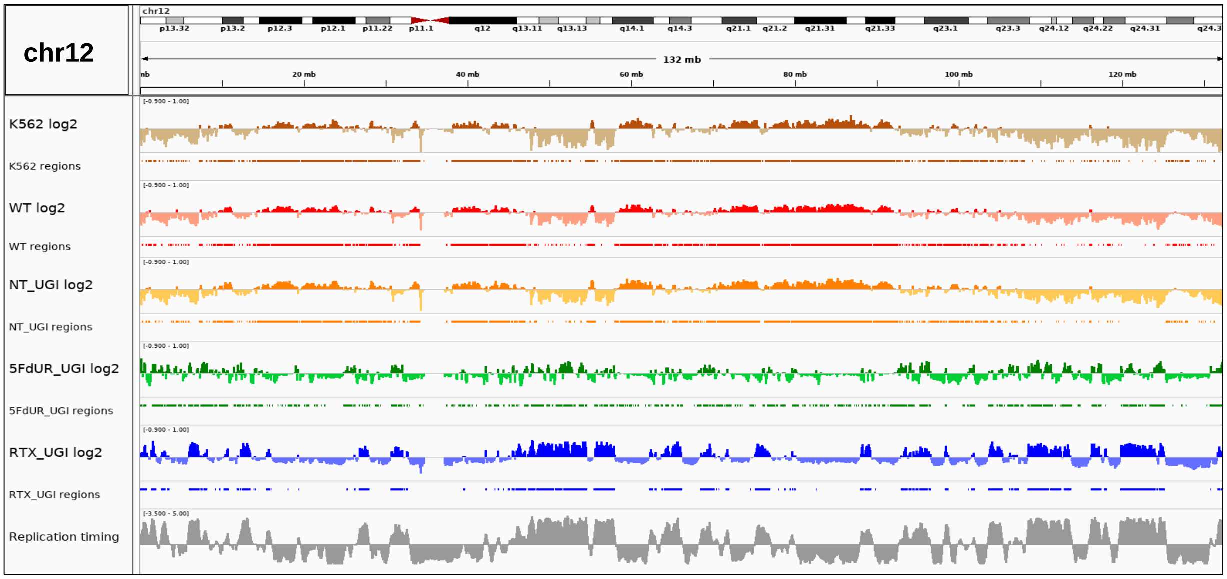
chr12
K562 log2
K562 regions
WT log2
WT regions
NT_UGI log2
NT_UGI regions
5FdUR_UGI log2
5FdUR_UGI regions
RTX_UGI log2
RTX_UGI regions
Replication timing

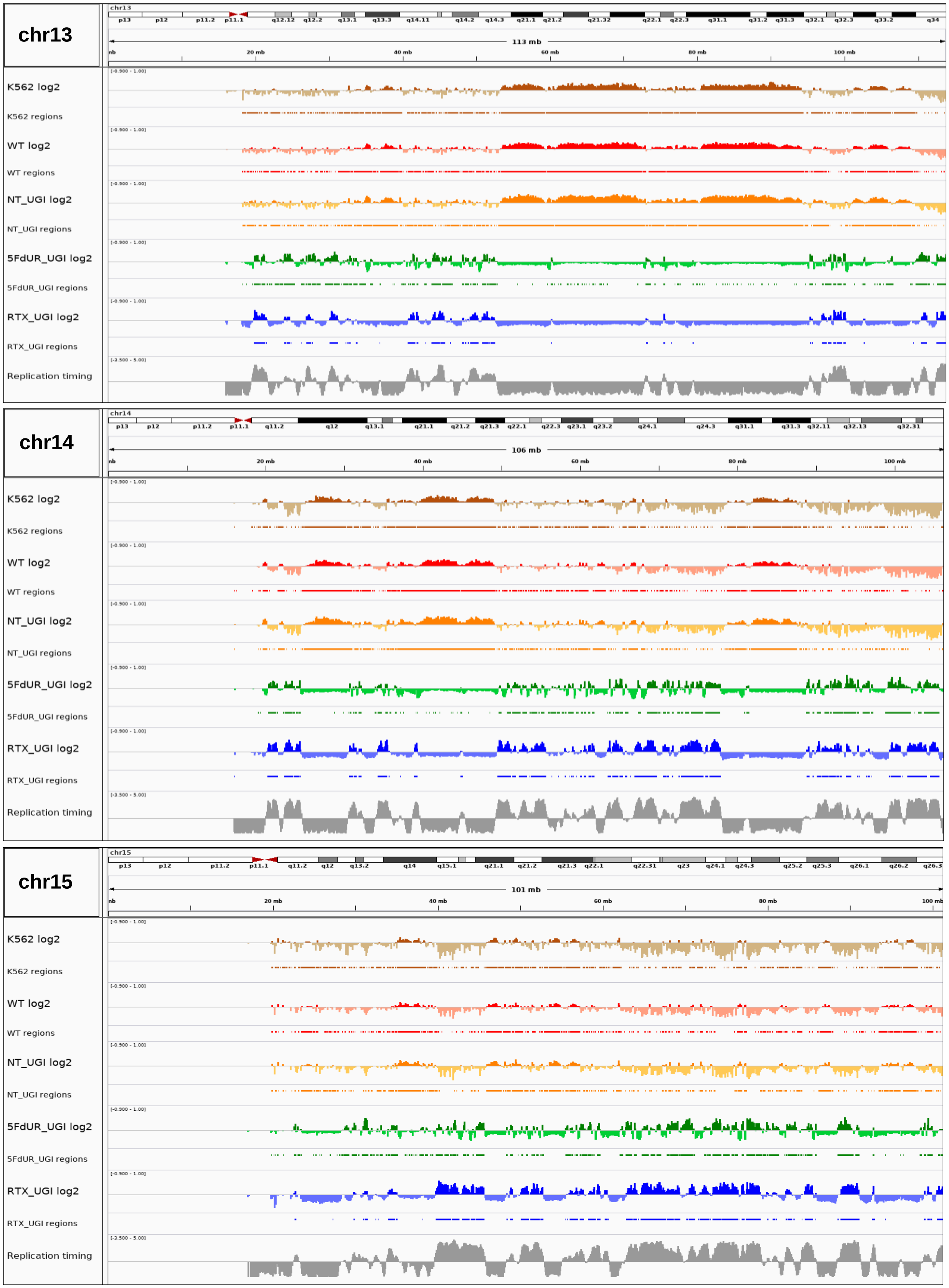
chr13
113 mb
K562 log2
K562 regions
WT log2
WT regions
NT_UGI log2
NT_UGI regions
5FdUR_UGI log2
5FdUR_UGI regions
RTX_UGI log2
RTX_UGI regions
Replication timing
chr14
106 mb
K562 log2
K562 regions
WT log2
WT regions
NT_UGI log2
NT_UGI regions
5FdUR_UGI log2
5FdUR_UGI regions
RTX_UGI log2
RTX_UGI regions
Replication timing
chr15
101 mb
K562 log2
K562 regions
WT log2
WT regions
NT_UGI log2
NT_UGI regions
5FdUR_UGI log2
5FdUR_UGI regions
RTX_UGI log2
RTX_UGI regions
Replication timing

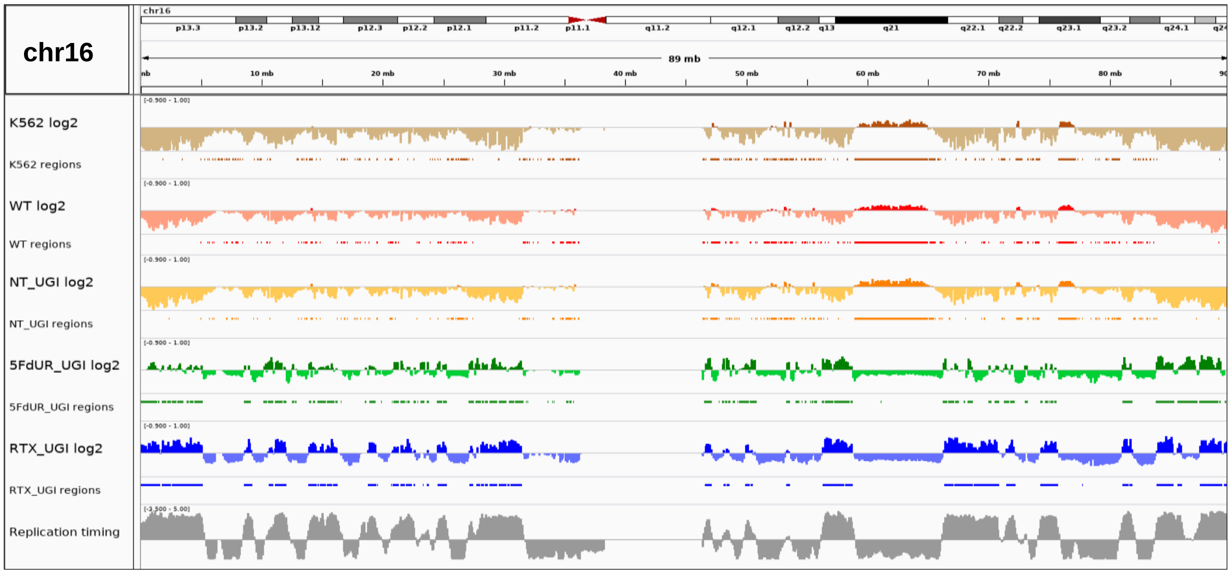
chr16
89 mb
K562 log2
K562 regions
WT log2
WT regions
NT_UGI log2
NT_UGI regions
5FdUR_UGI log2
5FdUR_UGI regions
RTX_UGI log2
RTX_UGI regions
Replication timing

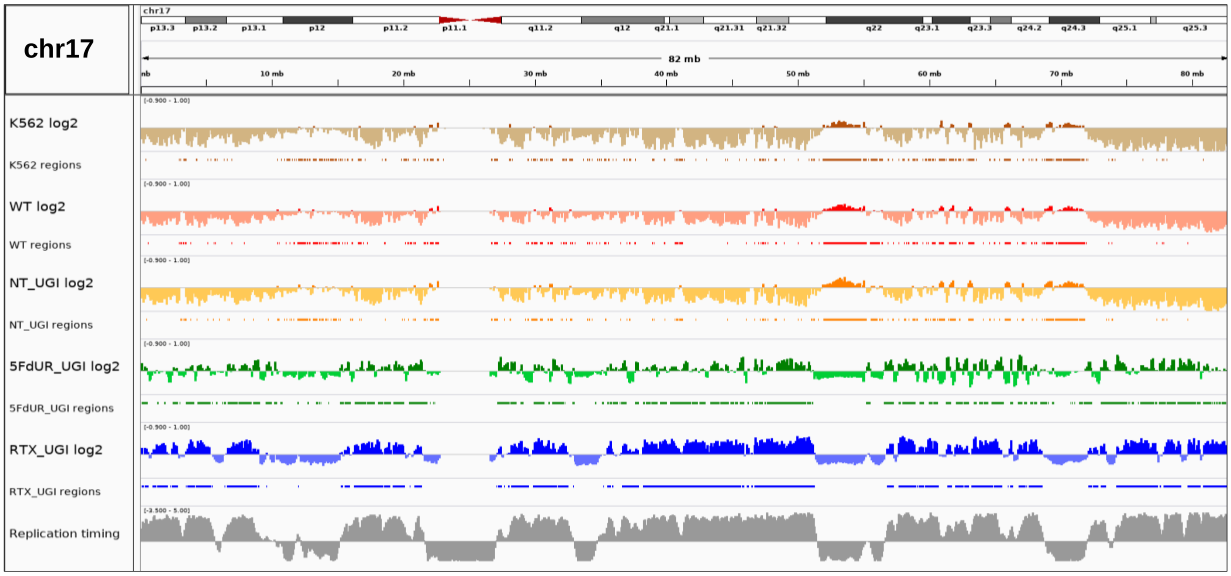
chr17
82 mb
K562 log2
K562 regions
WT log2
WT regions
NT_UGI log2
NT_UGI regions
5FdUR_UGI log2
5FdUR_UGI regions
RTX_UGI log2
RTX_UGI regions
Replication timing

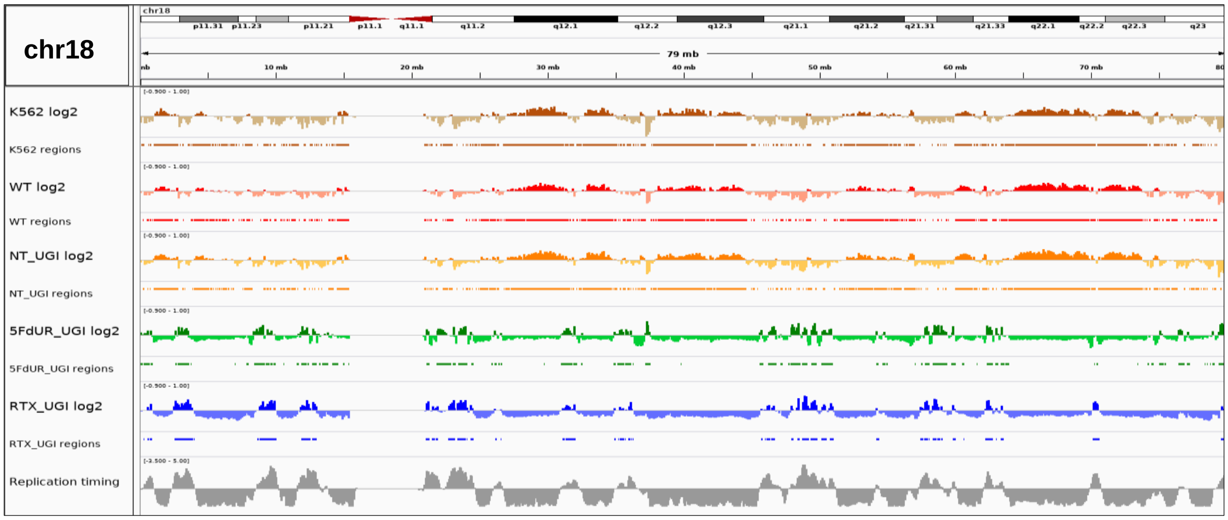
chr18
79 mb
K562 log2
K562 regions
WT log2
WT regions
NT_UGI log2
NT_UGI regions
5FdUR_UGI log2
5FdUR_UGI regions
RTX_UGI log2
RTX_UGI regions
Replication timing

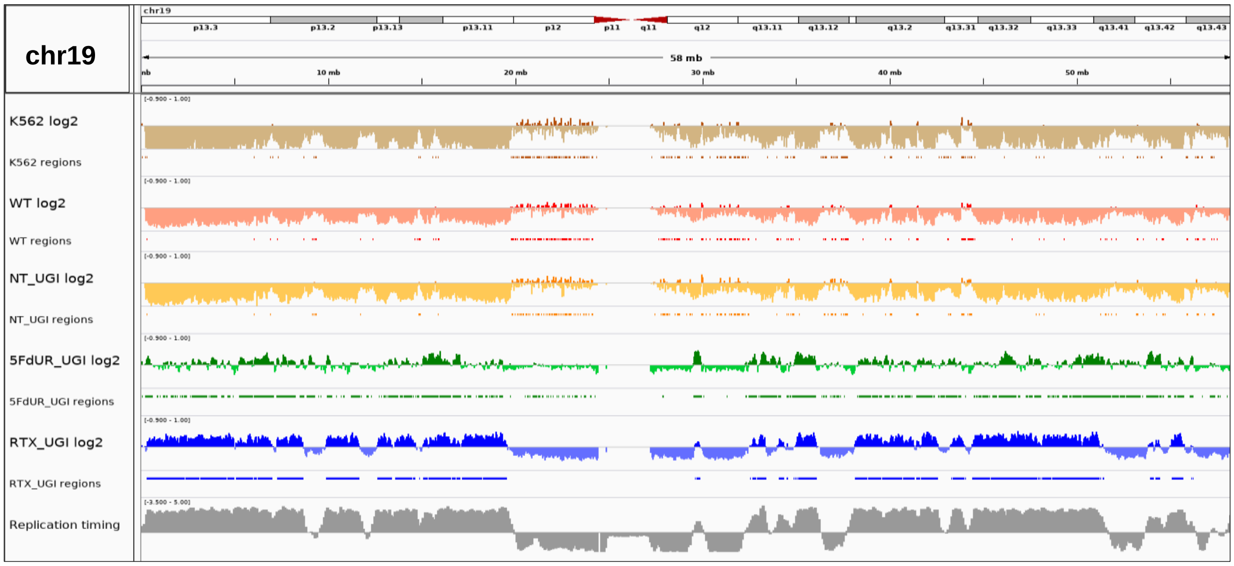
chr19
58 mb
K562 log2
K562 regions
WT log2
WT regions
NT_UGI log2
NT_UGI regions
5FdUR_UGI log2
5FdUR_UGI regions
RTX_UGI log2
RTX_UGI regions
Replication timing

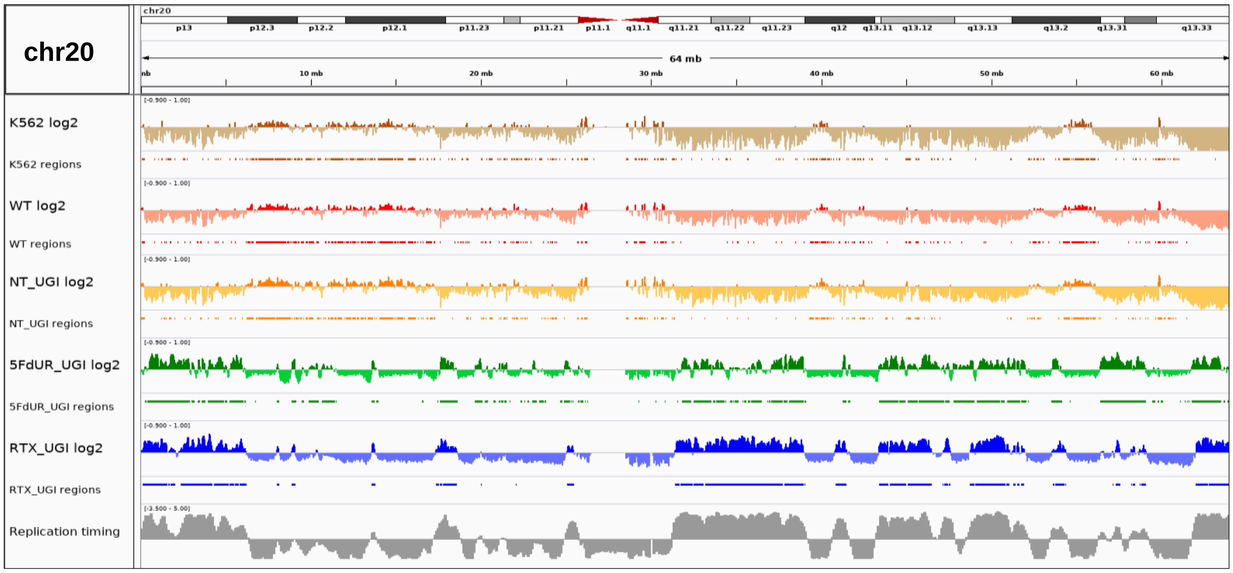
chr20
64 mb
K562 log2
K562 regions
WT log2
WT regions
NT_UGI log2
NT_UGI regions
5FdUR_UGI log2
5FdUR_UGI regions
RTX_UGI log2
RTX_UGI regions
Replication timing

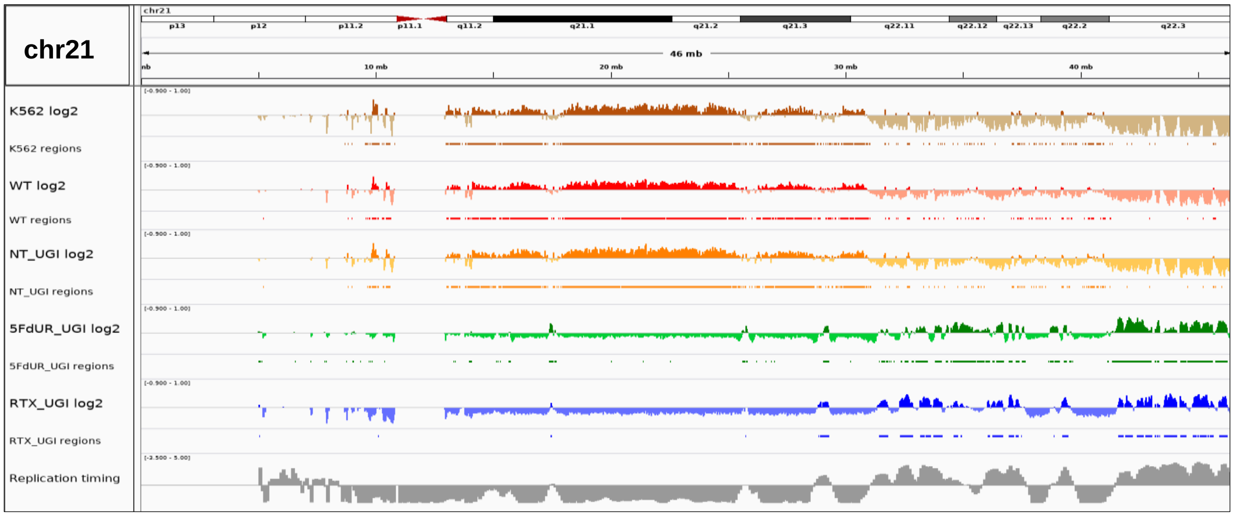
chr21
46 mb
K562 log2
K562 regions
WT log2
WT regions
NT_UGI log2
NT_UGI regions
5FdUR_UGI log2
5FdUR_UGI regions
RTX_UGI log2
RTX_UGI regions
Replication timing

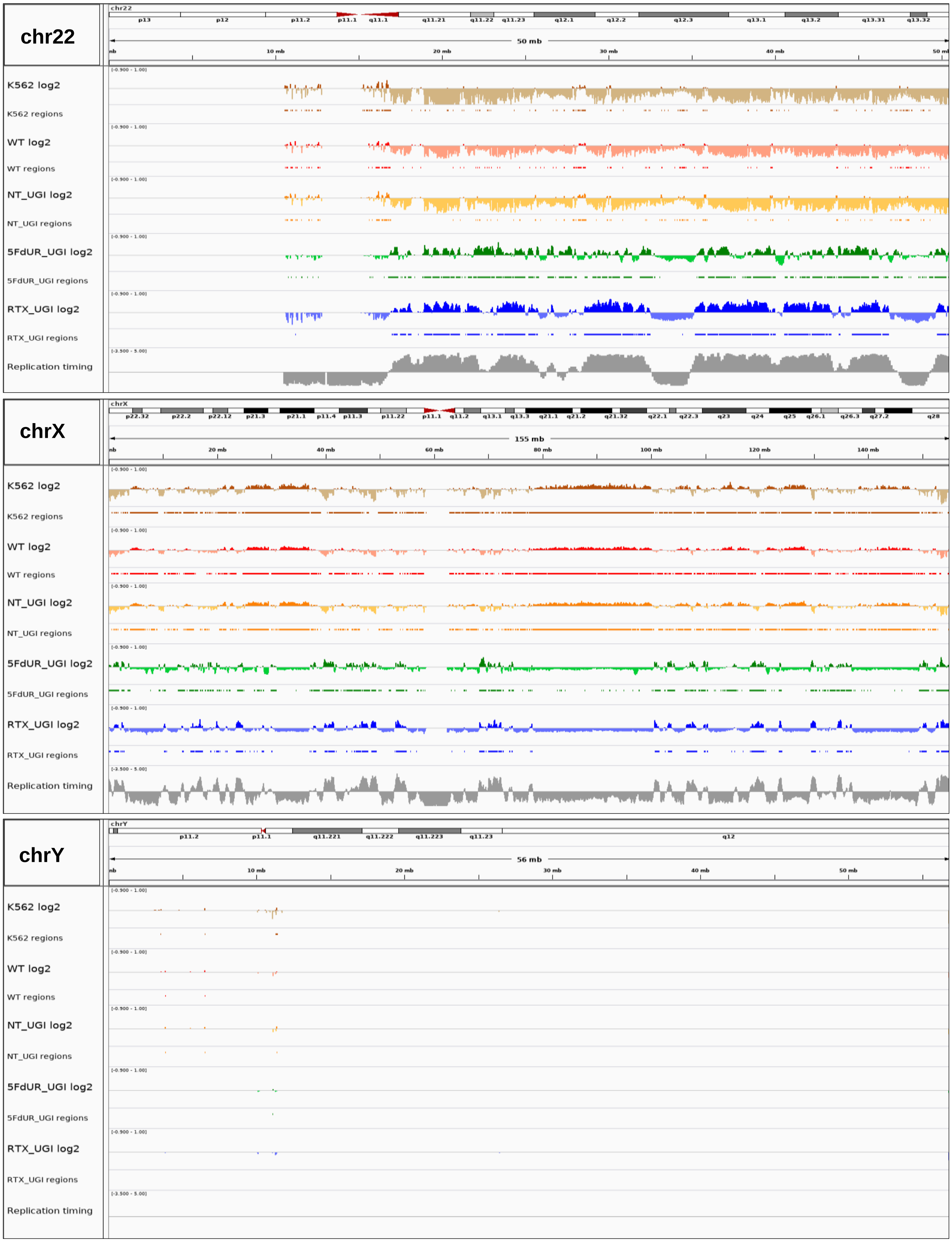
chr22
chrX
chrY
K562 log2
K562 regions
WT log2
WT regions
NT_UGI log2
NT_UGI regions
5FdUR_UGI log2
5FdUR_UGI regions
RTX_UGI log2
RTX_UGI regions
Replication timing
50 mb
155 mb
56 mb

**Appendix 2. Full table on combo scores from GIGGLE search.**

| factor | WT regions | NT_UGI regions | 5FdUR_UGI regions | RTX_ UGI regions | number of intervals | primary DB | primary_DB ID |
|---|---|---|---|---|---|---|---|
| histone markers | | | | | | | |
| H2A | -662.4 | -681.8 | 386.1 | 338.6 | 16198 | Cistrome | 51143 |
| H2A | -805.1 | -855.3 | 342.7 | 316.8 | 7396 | Cistrome | 51159 |
| H2AFZ | -411.8 | -422.5 | 443.7 | 375.9 | 16633 | Cistrome | 100929 |
| H2AFZ | -463.5 | -475.5 | 469.1 | 341.5 | 12876 | Cistrome | 100930 |
| H2AFZ | 151.7 | 162.2 | 407.3 | 329.8 | 70941 | ENCODE | ENCFF549VEQ.bed.gz |
| H2AZ | -569.3 | -589.3 | 418.2 | 423.6 | 56439 | Cistrome | 84716 |
| H2AZ | -523.2 | -541.9 | 414.8 | 417.0 | 59933 | Cistrome | 84715 |
| H2AZ | -72.3 | -83.5 | 411.4 | 352.1 | 100000 | Cistrome | 83065 |
| H2AZ | -735.2 | -819.4 | 460.5 | 352.4 | 16374 | Cistrome | 84720 |
| H2AZ | -923.8 | -943.5 | 437.0 | 318.1 | 3440 | Cistrome | 84719 |
| H2AZ | -915.8 | -931.1 | 457.5 | 322.7 | 9478 | Cistrome | 84717 |
| H2AZ | 3.4 | 5.0 | 374.7 | 294.7 | 100000 | Cistrome | 88446 |
| H2B | -12.5 | -19.7 | 10.3 | 1.9 | 1007 | Cistrome | 40190 |
| H2B | 0.1 | 0.1 | 0.0 | 0.0 | 571 | Cistrome | 40192 |
| H2Bub | -37.1 | -38.9 | 85.0 | 202.3 | 5475 | Cistrome | 40191 |
| H3.3 | -557.8 | -572.0 | 278.7 | 298.5 | 12756 | Cistrome | 77204 |
| H3.3 | -47.8 | -51.8 | 45.0 | 14.4 | 1016 | Cistrome | 74191 |
| H3K18ac | -1057.4 | -1029.4 | 311.6 | 346.7 | 15021 | Cistrome | 81141 |
| H3K18ac | -232.5 | -237.2 | 15.4 | 49.7 | 673 | Cistrome | 84122 |
| H3K18cr | -760.2 | -781.0 | 359.1 | 372.9 | 27293 | Cistrome | 88253 |
| H3K18cr | -1026.8 | -1007.8 | 307.7 | 335.0 | 12848 | Cistrome | 88250 |
| H3K18cr | -1204.4 | -1189.0 | 322.3 | 319.3 | 6848 | Cistrome | 88251 |
| H3K27ac | -775.5 | -815.0 | 467.8 | 587.3 | 100000 | Roadm. Epi. | E075-H3K27ac.narrowPeak.bed.gz |
| H3K27ac | -819.8 | -856.9 | 436.3 | 575.0 | 100000 | Roadm. Epi. | E106-H3K27ac.narrowPeak.bed.gz |
| H3K27ac | -464.0 | -473.2 | 278.9 | 580.6 | 45508 | ENCODE | ENCFF349LKU.bed.gz |
| H3K27ac | -530.0 | -544.7 | 322.6 | 528.8 | 40985 | ENCODE | ENCFF083ADY.bed.gz |
| H3K27ac | -470.1 | -474.3 | 364.6 | 521.8 | 100000 | Roadm. Epi. | E075-H3K27ac.broadPeak.bed.gz |
| H3K27ac | -535.0 | -553.0 | 344.1 | 515.9 | 100000 | Roadm. Epi. | E106-H3K27ac.broadPeak.bed.gz |
| H3K27ac | -558.2 | -583.1 | 370.5 | 482.6 | 100000 | Roadm. Epi. | E076-H3K27ac.narrowPeak.bed.gz |
| H3K27ac | -610.4 | -615.6 | 361.7 | 450.5 | 57085 | Cistrome | 61965 |
| H3K27ac | -569.6 | -570.2 | 281.1 | 460.8 | 14808 | Ensemble | homo_sapiens.GRCh38.HCT116.H3K27ac.SWEmbl_R0005.peaks.20190329.bed.gz |
| H3K27ac | -472.4 | -476.7 | 396.7 | 450.1 | 67892 | Cistrome | 66813 |
| H3K27ac | -499.3 | -505.1 | 318.1 | 437.3 | 50049 | Cistrome | 66991 |
| H3K27ac | -478.8 | -488.6 | 294.1 | 429.3 | 36552 | Cistrome | 101737 |
| H3K27ac | -564.6 | -564.2 | 362.4 | 417.4 | 51101 | Cistrome | 83014 |
| H3K27ac | -546.5 | -553.0 | 315.4 | 413.1 | 45759 | Cistrome | 45272 |
| H3K27ac | -447.0 | -450.7 | 310.0 | 414.2 | 49495 | Cistrome | 66992 |
| H3K27ac | -480.1 | -486.5 | 306.6 | 411.2 | 42374 | Cistrome | 66993 |
| H3K27ac | -450.7 | -454.2 | 299.3 | 402.4 | 41740 | Cistrome | 85866 |
| H3K27ac | -291.8 | -295.5 | 267.8 | 422.2 | 100000 | Roadm. Epi. | E076-H3K27ac.broadPeak.bed.gz |
| H3K27ac | -587.6 | -623.9 | 280.8 | 392.1 | 35902 | Cistrome | 61942 |
| H3K27ac | -558.1 | -557.6 | 256.2 | 390.6 | 22363 | Cistrome | 101739 |
| H3K27ac | -496.9 | -504.2 | 326.4 | 388.6 | 39573 | Cistrome | 83018 |
| H3K27ac | -661.2 | -667.2 | 298.8 | 380.4 | 36874 | Cistrome | 62110 |
| H3K27ac | -671.0 | -674.9 | 247.1 | 378.7 | 23506 | Cistrome | 61940 |
| H3K27ac | -631.7 | -641.3 | 289.7 | 376.5 | 35218 | Cistrome | 62112 |
| H3K27ac | -466.9 | -479.3 | 340.2 | 384.7 | 45825 | Cistrome | 85293 |
| H3K27ac | -703.9 | -693.4 | 200.8 | 379.0 | 4400 | Cistrome | 85867 |
| H3K27ac | -558.9 | -532.3 | 240.3 | 379.2 | 24442 | Cistrome | 61939 |
| H3K27ac | -689.6 | -695.2 | 300.6 | 376.0 | 32883 | Cistrome | 62109 |

| | | | | | | | |
|---|---|---|---|---|---|---|---|
| H3K27ac | -499.7 | -509.6 | 336.3 | 380.7 | 38279 | Cistrome | 85283 |
| H3K27ac | -390.9 | -398.3 | 271.3 | 379.8 | 44200 | Cistrome | 82335 |
| H3K27ac | -723.4 | -702.9 | 334.6 | 367.6 | 25172 | Cistrome | 66757 |
| H3K27ac | -753.2 | -747.0 | 254.0 | 365.2 | 11136 | Cistrome | 66393 |
| H3K27ac | -600.3 | -582.4 | 265.1 | 365.2 | 20517 | Cistrome | 42163 |
| H3K27ac | -373.2 | -377.1 | 285.0 | 372.4 | 50279 | Cistrome | 82334 |
| H3K27ac | -637.2 | -656.4 | 279.4 | 363.4 | 30630 | Cistrome | 62111 |
| H3K27ac | -530.6 | -536.9 | 343.6 | 371.6 | 5418 | Cistrome | 51146 |
| H3K27ac | -859.8 | -819.2 | 302.9 | 359.8 | 15455 | Cistrome | 81343 |
| H3K27ac | -660.4 | -652.0 | 297.5 | 360.4 | 16665 | Cistrome | 81342 |
| H3K27ac | -499.7 | -496.7 | 275.2 | 365.7 | 34708 | Cistrome | 84767 |
| H3K27ac | -438.4 | -434.9 | 266.0 | 362.3 | 42466 | Cistrome | 88909 |
| H3K27ac | -542.9 | -531.0 | 300.4 | 359.6 | 22842 | Cistrome | 81344 |
| H3K27ac | -787.2 | -797.3 | 285.0 | 356.7 | 13243 | Cistrome | 81345 |
| H3K27ac | -592.3 | -577.3 | 269.1 | 351.2 | 21201 | Cistrome | 66394 |
| H3K27ac | -615.0 | -632.6 | 294.5 | 359.9 | 35090 | Cistrome | 49529 |
| H3K27ac | -756.6 | -740.3 | 251.8 | 354.5 | 7375 | Cistrome | 84766 |
| H3K27ac | -494.7 | -481.8 | 241.7 | 351.2 | 24024 | Cistrome | 61941 |
| H3K27ac | -555.1 | -558.2 | 257.2 | 345.0 | 21349 | Cistrome | 66395 |
| H3K27ac | -485.1 | -472.3 | 265.8 | 354.3 | 16412 | Cistrome | 83017 |
| H3K27ac | -763.7 | -728.0 | 273.3 | 349.3 | 12964 | Cistrome | 81347 |
| H3K27ac | -526.3 | -560.1 | 292.7 | 346.5 | 22907 | Cistrome | 57096 |
| H3K27ac | -499.5 | -511.5 | 277.3 | 347.4 | 20922 | Cistrome | 87277 |
| H3K27ac | -510.7 | -504.5 | 296.6 | 346.3 | 33159 | Cistrome | 42164 |
| H3K27ac | -570.8 | -566.9 | 274.4 | 351.0 | 12793 | Cistrome | 87276 |
| H3K27ac | -736.3 | -729.9 | 259.3 | 336.8 | 13255 | Cistrome | 85454 |
| H3K27ac | -624.3 | -621.5 | 260.6 | 335.7 | 13528 | Cistrome | 81346 |
| H3K27ac | -716.3 | -763.0 | 291.3 | 335.8 | 23025 | Cistrome | 49530 |
| H3K27ac | -494.1 | -501.5 | 278.0 | 336.6 | 25411 | Cistrome | 85455 |
| H3K27ac | -351.9 | -361.4 | 230.4 | 319.6 | 6904 | Cistrome | 87275 |
| H3K27ac | -611.2 | -621.6 | 258.6 | 282.1 | 7309 | Cistrome | 87281 |
| H3K27ac | -112.6 | -80.9 | -3.4 | -21.7 | 876 | Cistrome | 67551 |
| H3K27me1 | 10.4 | 10.2 | -9.7 | 6.2 | 3757 | Cistrome | 69463 |
| H3K27me1 | -3.3 | -10.7 | -2.4 | 0.1 | 431 | Cistrome | 69460 |
| H3K27me1 | -2.6 | -1.2 | -1.9 | 0.0 | 574 | Cistrome | 69466 |
| H3K27me2 | 3.0 | 1.5 | -0.1 | 0.0 | 1246 | Cistrome | 69459 |
| H3K27me3 | -779.3 | -818.1 | 730.4 | 484.4 | 100000 | Roadm. Epi. | E075-H3K27me3.narrowPeak.bed.gz |
| H3K27me3 | -580.5 | -607.8 | 507.2 | 313.7 | 65497 | Roadm. Epi. | E076-H3K27me3.narrowPeak.bed.gz |
| H3K27me3 | -395.1 | -452.7 | 57.1 | 227.7 | 2705 | Cistrome | 51162 |
| H3K27me3 | -405.3 | -410.2 | 367.6 | 272.8 | 100000 | Roadm. Epi. | E075-H3K27me3.broadPeak.bed.gz |
| H3K27me3 | -485.6 | -509.7 | 469.0 | 229.6 | 100000 | Roadm. Epi. | E106-H3K27me3.narrowPeak.bed.gz |
| H3K27me3 | -281.1 | -293.5 | 281.0 | 174.7 | 100000 | Roadm. Epi. | E076-H3K27me3.broadPeak.bed.gz |
| H3K27me3 | -232.9 | -234.8 | 194.5 | 66.9 | 100000 | Roadm. Epi. | E106-H3K27me3.broadPeak.bed.gz |
| H3K27me3 | -65.5 | -57.5 | 42.5 | 30.4 | 699 | Cistrome | 51147 |
| H3K27me3 | -159.1 | -155.8 | 103.1 | -168.7 | 199001 | Ensemble | homo_sapiens.GRCh38.HCT116.H3K27me3.ccat_histone.peaks.20190329.mergedd1000.bed.gz |
| H3K27me3 | -279.5 | -289.5 | -31.4 | -341.4 | 100000 | ENCODE | ENCFF806AYM.bed.gz |
| H3K36me3 | -499.5 | -519.2 | 208.7 | 837.8 | 100000 | Roadm. Epi. | E075-H3K36me3.narrowPeak.bed.gz |
| H3K36me3 | -397.7 | -414.4 | 97.1 | 771.3 | 100000 | Roadm. Epi. | E106-H3K36me3.narrowPeak.bed.gz |

| | | | | | | | |
|---|---|---|---|---|---|---|---|
| H3K36me3 | -329.0 | -341.3 | 122.7 | 740.9 | 100000 | Roadm. Epi. | E076-H3K36me3.narrowPeak.bed.gz |
| H3K36me3 | -261.3 | -269.3 | 48.6 | 717.1 | 72341 | ENCODE | ENCFF029GQD.bed.gz |
| H3K36me3 | -186.6 | -188.5 | 44.8 | 530.2 | 43916 | Ensemble | homo_sapiens.GRCh38.HCT116.H3K36me3.ccat_histone.peaks.20190329.mergedd1000.bed.gz |
| H3K36me3 | -202.0 | -207.3 | 182.7 | 491.3 | 100000 | Roadm. Epi. | E076-H3K36me3.broadPeak.bed.gz |
| H3K36me3 | -246.2 | -251.3 | 191.1 | 492.8 | 100000 | Roadm. Epi. | E075-H3K36me3.broadPeak.bed.gz |
| H3K36me3 | -194.0 | -198.7 | 77.7 | 462.1 | 100000 | Roadm. Epi. | E106-H3K36me3.broadPeak.bed.gz |
| H3K36me3 | -35.1 | -44.2 | 2.7 | 137.8 | 563 | Cistrome | 51149, 51164 |
| H3K4me1 | -1067.6 | -1119.8 | 573.3 | 718.4 | 100000 | Roadm. Epi. | E075-H3K4me1.narrowPeak.bed.gz |
| H3K4me1 | -457.7 | -477.3 | 438.2 | 712.2 | 100000 | ENCODE | ENCFF986BGX.bed.gz |
| H3K4me1 | -446.2 | -456.4 | 331.5 | 629.8 | 99735 | Ensemble | homo_sapiens.GRCh38.HCT116.H3K4me1.ccat_histone.peaks.20190329.bed.gz |
| H3K4me1 | -418.0 | -426.3 | 445.5 | 590.5 | 100000 | Cistrome | 83177 |
| H3K4me1 | -588.2 | -613.8 | 442.3 | 578.7 | 100000 | Cistrome | 70068 |
| H3K4me1 | -442.6 | -463.1 | 333.8 | 575.7 | 82838 | ENCODE | ENCFF963BLP.bed.gz |
| H3K4me1 | -421.0 | -435.7 | 417.8 | 553.9 | 100000 | Cistrome | 101556 |
| H3K4me1 | -458.5 | -479.1 | 401.2 | 550.3 | 97772 | Cistrome | 74630 |
| H3K4me1 | -399.7 | -412.6 | 431.8 | 553.2 | 100000 | Cistrome | 85458 |
| H3K4me1 | -494.2 | -504.1 | 470.2 | 543.8 | 97947 | Cistrome | 51161 |
| H3K4me1 | -597.6 | -621.3 | 433.2 | 538.4 | 87601 | Cistrome | 82750 |
| H3K4me1 | -543.4 | -548.6 | 428.5 | 571.1 | 100000 | Roadm. Epi. | E075-H3K4me1.broadPeak.bed.gz |
| H3K4me1 | -464.1 | -479.0 | 424.4 | 519.2 | 100000 | Cistrome | 82751 |
| H3K4me1 | -445.5 | -467.6 | 428.7 | 514.0 | 100000 | Cistrome | 82748 |
| H3K4me1 | -570.0 | -600.9 | 430.1 | 525.1 | 86089 | Cistrome | 70076 |
| H3K4me1 | -611.9 | -639.4 | 353.5 | 516.5 | 100000 | Roadm. Epi. | E106-H3K4me1.narrowPeak.bed.gz |
| H3K4me1 | -451.4 | -468.9 | 414.6 | 518.3 | 88273 | Cistrome | 70075 |
| H3K4me1 | -449.8 | -467.4 | 393.0 | 504.4 | 91410 | Cistrome | 70067 |
| H3K4me1 | -360.2 | -371.4 | 400.7 | 496.1 | 100000 | Cistrome | 88015 |
| H3K4me1 | -344.7 | -358.5 | 403.3 | 488.6 | 100000 | Cistrome | 88811 |
| H3K4me1 | -465.9 | -485.4 | 383.7 | 494.5 | 78930 | Cistrome | 9246 |
| H3K4me1 | -432.9 | -448.9 | 321.2 | 485.1 | 100000 | Roadm. Epi. | E106-H3K4me1.broadPeak.bed.gz |
| H3K4me1 | -409.1 | -418.8 | 389.0 | 476.7 | 86738 | Cistrome | 84634 |
| H3K4me1 | -334.8 | -367.7 | 270.2 | 482.3 | 79205 | Cistrome | 57095 |
| H3K4me1 | -557.7 | -576.3 | 454.8 | 474.9 | 48818 | Cistrome | 51145 |
| H3K4me1 | -917.8 | -919.0 | 467.5 | 442.3 | 37547 | Cistrome | 42909 |
| H3K4me1 | -509.3 | -516.7 | 365.3 | 435.0 | 41288 | Cistrome | 101557 |
| H3K4me1 | -673.3 | -714.7 | 431.7 | 422.8 | 29827 | Cistrome | 42908 |
| H3K4me1 | -351.9 | -335.7 | 395.7 | 418.6 | 4238 | Cistrome | 72104 |
| H3K4me1 | -407.6 | -423.6 | 268.8 | 391.8 | 100000 | Roadm. Epi. | E076-H3K4me1.narrowPeak.bed.gz |
| H3K4me1 | -464.4 | -526.0 | 398.3 | 406.6 | 4839 | Cistrome | 72106 |
| H3K4me1 | -469.0 | -485.7 | 370.5 | 402.0 | 47917 | Cistrome | 88979 |
| H3K4me1 | -622.3 | -647.3 | 403.7 | 380.1 | 35174 | Cistrome | 86455 |
| H3K4me1 | -264.0 | -271.0 | 270.3 | 375.7 | 100000 | Roadm. Epi. | E076-H3K4me1.broadPeak.bed.gz |
| H3K4me1 | -282.1 | -291.7 | 265.9 | 350.0 | 34853 | Cistrome | 101558 |
| H3K4me1 | -31.1 | -45.8 | 87.4 | 86.6 | 567 | Cistrome | 84587 |
| H3K4me1 | -63.1 | -55.1 | 16.5 | 54.3 | 422 | Cistrome | 45270 |
| H3K4me2 | -449.7 | -459.2 | 403.0 | 536.0 | 69442 | ENCODE | ENCFF915XUY.bed.gz |

| | | | | | | | |
|---|---|---|---|---|---|---|---|
| H3K4me2 | -417.2 | -426.3 | 389.9 | 464.5 | 79557 | Cistrome | 101374 |
| H3K4me2 | -784.9 | -809.3 | 396.2 | 459.7 | 45297 | Cistrome | 82749 |
| H3K4me2 | -478.0 | -484.2 | 380.5 | 423.2 | 62083 | Cistrome | 101375 |
| H3K4me2 | -913.1 | -928.0 | 427.5 | 417.0 | 40745 | Cistrome | 42911 |
| H3K4me2 | -461.0 | -462.9 | 400.8 | 411.7 | 64397 | Cistrome | 42910 |
| H3K4me2 | -530.7 | -539.2 | 364.8 | 408.0 | 52922 | Cistrome | 70073 |
| H3K4me2 | -462.1 | -469.6 | 370.4 | 404.8 | 69348 | Cistrome | 83378 |
| H3K4me2 | -792.5 | -820.5 | 345.9 | 401.4 | 39615 | Cistrome | 70066 |
| H3K4me2 | -640.8 | -652.7 | 361.4 | 396.1 | 47492 | Cistrome | 83353 |
| H3K4me2 | -638.5 | -667.8 | 367.7 | 393.7 | 42419 | Cistrome | 70074 |
| H3K4me2 | -634.2 | -650.1 | 338.4 | 387.1 | 40335 | Cistrome | 70065 |
| H3K4me2 | -866.8 | -880.0 | 395.5 | 375.0 | 23301 | Cistrome | 83307 |
| H3K4me2 | -955.8 | -968.7 | 384.0 | 368.5 | 29310 | Cistrome | 70079 |
| H3K4me2 | -897.0 | -929.1 | 405.9 | 365.7 | 17134 | Cistrome | 88737 |
| H3K4me2 | -1277.3 | -1242.0 | 355.7 | 357.1 | 21206 | Cistrome | 70081 |
| H3K4me2 | -1708.3 | -1711.7 | 452.3 | 329.8 | 7869 | Cistrome | 83376 |
| H3K4me3 | -762.4 | -792.4 | 481.1 | 509.2 | 84678 | Roadm. Epi. | E075-H3K4me3.narrowPeak.bed.gz |
| H3K4me3 | -607.3 | -628.3 | 384.9 | 437.0 | 31924 | ENCODE | ENCFF023MGT.bed.gz |
| H3K4me3 | -861.7 | -900.1 | 406.5 | 422.7 | 26743 | ENCODE | ENCFF575AUU.bed.gz |
| H3K4me3 | -454.8 | -463.1 | 368.7 | 443.1 | 100000 | Roadm. Epi. | E075-H3K4me3.broadPeak.bed.gz |
| H3K4me3 | -1000.1 | -1107.1 | 392.5 | 403.8 | 17658 | Ensemble | homo_sapiens.GRCh38.HCT116.H3K4me3.SWEmbl_R0005.peaks.20190329.bed.gz |
| H3K4me3 | -929.4 | -946.7 | 396.7 | 384.1 | 26517 | Cistrome | 51144 |
| H3K4me3 | -793.3 | -813.5 | 397.4 | 379.9 | 34446 | Cistrome | 45271 |
| H3K4me3 | -877.5 | -892.1 | 391.2 | 372.2 | 31205 | Cistrome | 92369 |
| H3K4me3 | -476.9 | -485.1 | 365.1 | 373.8 | 48602 | Cistrome | 85292 |
| H3K4me3 | -663.8 | -686.3 | 378.1 | 377.2 | 33338 | Cistrome | 102153 |
| H3K4me3 | -854.8 | -831.0 | 381.7 | 369.9 | 29334 | Cistrome | 42912 |
| H3K4me3 | -590.4 | -600.5 | 361.0 | 374.5 | 31737 | Cistrome | 102154 |
| H3K4me3 | -890.3 | -906.2 | 397.2 | 366.7 | 30486 | Cistrome | 42161 |
| H3K4me3 | -951.8 | -974.8 | 390.3 | 368.9 | 28373 | Cistrome | 89284 |
| H3K4me3 | -876.5 | -903.5 | 387.6 | 367.9 | 29898 | Cistrome | 92368 |
| H3K4me3 | -976.1 | -997.0 | 386.6 | 368.1 | 27614 | Cistrome | 89285 |
| H3K4me3 | -1105.8 | -1139.3 | 386.2 | 367.7 | 25156 | Cistrome | 42913 |
| H3K4me3 | -841.8 | -868.2 | 391.8 | 371.5 | 26578 | Cistrome | 81264 |
| H3K4me3 | -966.0 | -1123.7 | 246.6 | 370.4 | 2983 | Cistrome | 81266 |
| H3K4me3 | -840.6 | -854.4 | 378.9 | 366.2 | 24080 | Cistrome | 88713 |
| H3K4me3 | -1124.9 | -1201.8 | 376.7 | 362.6 | 19739 | Cistrome | 88725 |
| H3K4me3 | -1027.5 | -1086.1 | 378.2 | 358.3 | 20453 | Cistrome | 88698 |
| H3K4me3 | -987.5 | -1024.7 | 388.2 | 352.6 | 27726 | Cistrome | 42160 |
| H3K4me3 | -1004.3 | -1066.4 | 397.1 | 353.8 | 27346 | Cistrome | 81633 |
| H3K4me3 | -611.6 | -640.5 | 385.6 | 368.8 | 39820 | Cistrome | 51160 |
| H3K4me3 | -998.4 | -1042.3 | 396.8 | 351.1 | 26055 | Cistrome | 56017 |
| H3K4me3 | -1169.5 | -1234.0 | 391.8 | 353.4 | 22478 | Cistrome | 57094 |
| H3K4me3 | -1059.0 | -1076.0 | 391.5 | 353.9 | 25236 | Cistrome | 81265 |
| H3K4me3 | -1014.3 | -1060.9 | 415.8 | 360.4 | 27993 | Cistrome | 42162 |
| H3K4me3 | -959.6 | -1009.1 | 388.8 | 350.7 | 25988 | Cistrome | 56016 |
| H3K4me3 | -768.8 | -779.9 | 374.0 | 352.9 | 22224 | Cistrome | 88697 |
| H3K4me3 | -1188.4 | -1247.6 | 334.9 | 350.5 | 16524 | Cistrome | 70064 |
| H3K4me3 | -1053.6 | -1081.8 | 392.2 | 343.0 | 24912 | Cistrome | 81625 |
| H3K4me3 | -876.9 | -915.9 | 398.2 | 345.6 | 25654 | Cistrome | 56019 |
| H3K4me3 | -1073.8 | -1174.4 | 357.5 | 347.5 | 18426 | Cistrome | 70071 |
| H3K4me3 | -1174.0 | -1239.4 | 334.2 | 341.3 | 16204 | Cistrome | 70063 |
| H3K4me3 | -1078.5 | -1140.5 | 366.0 | 344.8 | 17739 | Cistrome | 85447 |
| H3K4me3 | -1054.1 | -1148.6 | 376.7 | 344.9 | 19186 | Cistrome | 85451 |
| H3K4me3 | -907.4 | -889.7 | 327.6 | 336.0 | 19549 | Cistrome | 38319 |

| | | | | | | | |
|---|---|---|---|---|---|---|---|
| H3K4me3 | -907.4 | -889.7 | 327.6 | 336.0 | 19549 | Cistrome | 8440 |
| H3K4me3 | -1176.4 | -1234.7 | 363.8 | 338.6 | 19113 | Cistrome | 81635 |
| H3K4me3 | -1143.4 | -1180.0 | 355.2 | 342.4 | 19410 | Cistrome | 70072 |
| H3K4me3 | -1176.5 | -1219.3 | 357.4 | 341.9 | 16561 | Cistrome | 85449 |
| H3K4me3 | -530.8 | -546.0 | 359.6 | 356.5 | 72243 | Roadm. Epi. | E076-H3K4me3.narrowPeak.bed.gz |
| H3K4me3 | -1234.7 | -1254.3 | 340.7 | 333.2 | 18978 | Cistrome | 72103 |
| H3K4me3 | -1248.3 | -1268.7 | 346.3 | 336.6 | 15522 | Cistrome | 85446 |
| H3K4me3 | -1196.1 | -1226.9 | 350.6 | 334.5 | 16408 | Cistrome | 85452 |
| H3K4me3 | -1137.1 | -1223.9 | 361.2 | 334.6 | 17932 | Cistrome | 85453 |
| H3K4me3 | -1053.0 | -1067.9 | 335.9 | 329.0 | 15627 | Cistrome | 72105 |
| H3K4me3 | -1229.8 | -1222.5 | 342.0 | 333.5 | 15943 | Cistrome | 85448 |
| H3K4me3 | -1249.0 | -1267.8 | 354.5 | 332.3 | 16383 | Cistrome | 85450 |
| H3K4me3 | -975.9 | -1000.3 | 379.7 | 324.4 | 23519 | Cistrome | 56018 |
| H3K4me3 | -863.5 | -860.9 | 346.1 | 322.3 | 21480 | Cistrome | 38321 |
| H3K4me3 | -660.4 | -688.0 | 317.2 | 317.4 | 17967 | Cistrome | 49532 |
| H3K4me3 | -409.0 | -425.8 | 245.4 | 280.3 | 68861 | Roadm. Epi. | E106-H3K4me3.narrowPeak.bed.gz |
| H3K4me3 | -271.5 | -277.3 | 237.3 | 285.9 | 100000 | Roadm. Epi. | E076-H3K4me3.broadPeak.bed.gz |
| H3K4me3 | -147.2 | -149.2 | 132.5 | 223.6 | 100000 | Roadm. Epi. | E106-H3K4me3.broadPeak.bed.gz |
| H3K79me2 | -852.1 | -903.2 | 259.1 | 440.9 | 10541 | Cistrome | 101708 |
| H3K79me2 | -630.1 | -700.2 | 225.8 | 418.7 | 8655 | Cistrome | 101707 |
| H3K79me2 | -46.9 | -45.9 | 25.2 | 245.6 | 2368 | Cistrome | 83603 |
| H3K79me2 | 0.0 | 0.0 | 160.0 | 195.0 | 20855 | Cistrome | 88455 |
| H3K9ac | -781.0 | -791.5 | 539.8 | 674.8 | 100000 | Roadm. Epi. | E075-H3K9ac.narrowPeak.bed.gz |
| H3K9ac | -416.4 | -424.9 | 372.4 | 540.1 | 100000 | Roadm. Epi. | E075-H3K9ac.broadPeak.bed.gz |
| H3K9ac | -596.9 | -613.3 | 349.0 | 510.0 | 36097 | ENCODE | ENCFF724JXS.bed.gz |
| H3K9ac | -546.1 | -544.9 | 367.3 | 417.9 | 49287 | Cistrome | 87280 |
| H3K9ac | -501.4 | -516.8 | 349.9 | 426.2 | 60918 | Roadm. Epi. | E076-H3K9ac.narrowPeak.bed.gz |
| H3K9ac | -722.5 | -733.0 | 353.8 | 408.8 | 32466 | Cistrome | 100642 |
| H3K9ac | -566.9 | -567.6 | 362.4 | 393.6 | 44150 | Cistrome | 87279 |
| H3K9ac | -687.9 | -687.5 | 334.2 | 392.5 | 29925 | Cistrome | 100641 |
| H3K9ac | -515.3 | -520.4 | 356.6 | 391.3 | 40603 | Cistrome | 87278 |
| H3K9ac | -568.7 | -581.4 | 369.2 | 376.7 | 40039 | Cistrome | 87273 |
| H3K9ac | -510.3 | -511.1 | 328.9 | 337.4 | 24383 | Cistrome | 82746 |
| H3K9ac | -535.5 | -529.2 | 313.0 | 335.6 | 18541 | Cistrome | 87272 |
| H3K9ac | -156.2 | -152.2 | 199.1 | 354.7 | 100000 | Roadm. Epi. | E076-H3K9ac.broadPeak.bed.gz |
| H3K9ac | -621.7 | -643.6 | 315.6 | 318.0 | 20226 | Cistrome | 82747 |
| H3K9ac | -835.8 | -857.0 | 337.7 | 298.0 | 13135 | Cistrome | 82745 |
| H3K9ac | -649.1 | -677.5 | 278.0 | 284.8 | 5854 | Cistrome | 82744 |
| H3K9me2 | -646.1 | -707.2 | 522.2 | 332.6 | 16338 | Cistrome | 85108 |
| H3K9me2 | -0.6 | -0.3 | 0.6 | 0.0 | 1493 | Cistrome | 54504 |
| H3K9me2 | 290.7 | 334.9 | -1.0 | -3.7 | 4735 | ENCODE | ENCFF045ELD.bed.gz |
| H3K9me2 | -107.3 | -96.9 | -97.4 | -132.3 | 612 | Cistrome | 93700 |
| H3K9me2 | -109.6 | -84.8 | -108.3 | -200.0 | 726 | Cistrome | 93701 |
| H3K9me3 | -11.0 | -14.9 | -43.1 | -25.0 | 471 | Cistrome | 51148 |
| H3K9me3 | -32.1 | -26.8 | -57.7 | -55.6 | 715 | Cistrome | 51163 |
| H3K9me3 | -17.1 | -14.8 | 4.5 | -45.7 | 100000 | Roadm. Epi. | E075-H3K9me3.broadPeak.bed.gz |
| H3K9me3 | 2.2 | 8.5 | -0.3 | -103.1 | 100000 | Roadm. Epi. | E076-H3K9me3.broadPeak.bed.gz |
| H3K9me3 | -50.2 | -20.6 | -17.9 | -122.4 | 64003 | Roadm. Epi. | E076-H3K9me3.narrowPeak.bed.gz |
| H3K9me3 | -169.0 | -180.4 | -4.8 | -185.6 | 100000 | Roadm. Epi. | E075-H3K9me3.narrowPeak.bed.gz |
| H3K9me3 | 85.2 | 95.0 | -146.1 | -248.3 | 100000 | Roadm. Epi. | E106-H3K9me3.broadPeak.bed.gz |

| | | | | | | | |
|---|---|---|---|---|---|---|---|
| H3K9me3 | -31.2 | -38.6 | -233.9 | -329.2 | 100000 | Roadm. Epi. | E106-H3K9me3.narrowPeak.bed.gz |
| H3K9me3 | 198.1 | 208.8 | -407.1 | -420.4 | 116955 | Ensemble | homo_sapiens.GRCh38.HCT116.H3K9me3.ccat_histone.peaks.20190329.mergedd1000.bed.gz |
| H3K9me3 | 312.5 | 327.6 | -664.1 | -645.0 | 100000 | ENCODE | ENCFF580MMU.bed.gz |
| H4K16ac | -595.3 | -607.4 | 457.0 | 503.2 | 2290 | Cistrome | 83887 |
| H4K16ac | -842.0 | -847.7 | 383.7 | 382.0 | 25148 | Cistrome | 83526 |
| H4K20me1 | -27.0 | -34.5 | 0.0 | 481.4 | 20508 | ENCODE | ENCFF937WLG.bed.gz |
| H4K20me3 | -219.9 | -227.2 | 237.6 | 222.4 | 39942 | Cistrome | 88457 |
| **polymerase II - active transcription** | | | | | | | |
| POLR2A | -809.5 | -684.6 | 143.4 | 568.1 | 3029 | Cistrome | 82412 |
| POLR2A | -515.1 | -535.9 | 276.6 | 507.2 | 66413 | Cistrome | 34398 |
| POLR2A | -760.5 | -775.3 | 389.4 | 463.8 | 31718 | Cistrome | 82333 |
| POLR2A | -542.0 | -549.1 | 392.0 | 452.9 | 55528 | Cistrome | 46201 |
| POLR2A | -951.7 | -857.4 | 94.8 | 469.0 | 4543 | Cistrome | 55679 |
| POLR2A | -452.0 | -388.3 | 34.8 | 461.3 | 1700 | Cistrome | 86621 |
| POLR2A | -924.7 | -929.8 | 366.9 | 443.1 | 24151 | Cistrome | 92091 |
| POLR2A | -652.8 | -656.2 | 400.9 | 438.1 | 40923 | Cistrome | 39917 |
| POLR2A | -819.1 | -807.4 | 351.8 | 439.6 | 23335 | Cistrome | 88903 |
| POLR2A | -644.3 | -649.4 | 395.9 | 438.9 | 41125 | Cistrome | 39921 |
| POLR2A | -963.2 | -962.3 | 360.7 | 436.8 | 23301 | Cistrome | 89219 |
| POLR2A | -940.5 | -950.6 | 399.7 | 434.6 | 24689 | Cistrome | 93206 |
| POLR2A | -1014.6 | -1051.9 | 362.7 | 429.4 | 20360 | Cistrome | 89218 |
| POLR2A | -903.8 | -909.0 | 350.1 | 429.6 | 20196 | Cistrome | 83958 |
| POLR2A | -841.8 | -849.6 | 380.7 | 419.2 | 25213 | Cistrome | 88093 |
| POLR2A | -874.5 | -864.5 | 338.6 | 420.7 | 19821 | Cistrome | 83956 |
| POLR2A | -857.3 | -845.8 | 348.6 | 415.3 | 20421 | Cistrome | 83953 |
| POLR2A | -919.5 | -918.3 | 380.1 | 414.1 | 21544 | Cistrome | 93205 |
| POLR2A | -843.9 | -922.3 | 429.4 | 417.8 | 45426 | Cistrome | 57088 |
| POLR2A | -601.9 | -573.6 | 26.7 | 417.0 | 2749 | Cistrome | 55677 |
| POLR2A | -936.3 | -945.2 | 330.0 | 409.7 | 17288 | Cistrome | 83950 |
| POLR2A | -949.2 | -1010.8 | 395.7 | 402.0 | 27176 | Cistrome | 5661 |
| POLR2A | -793.7 | -794.4 | 350.6 | 404.8 | 21269 | Cistrome | 83959 |
| POLR2A | -742.6 | -731.6 | 341.6 | 404.3 | 23176 | Cistrome | 88100 |
| POLR2A | -743.5 | -732.8 | 352.1 | 401.4 | 24764 | Cistrome | 82337 |
| POLR2A | -724.8 | -701.9 | 341.4 | 399.5 | 23854 | Cistrome | 88099 |
| POLR2A | -911.1 | -945.4 | 373.6 | 394.0 | 20284 | Cistrome | 83342 |
| POLR2A | -1097.7 | -1135.9 | 341.8 | 389.1 | 16764 | ENCODE | ENCFF271RGE.bed.gz |
| POLR2A | -865.7 | -869.2 | 345.1 | 391.7 | 18859 | Cistrome | 83957 |
| POLR2A | -1180.4 | -1255.8 | 339.8 | 387.4 | 15512 | ENCODE | ENCFF786BQV.bed.gz |
| POLR2A | -710.6 | -701.0 | 340.5 | 396.0 | 22757 | Cistrome | 83951 |
| POLR2A | -1014.6 | -1007.6 | 371.0 | 387.3 | 18540 | Cistrome | 88094 |
| POLR2A | -835.7 | -820.7 | 364.4 | 387.7 | 21013 | Cistrome | 88096 |
| POLR2A | -670.8 | -647.1 | 329.2 | 385.3 | 22512 | Cistrome | 88091 |
| POLR2A | -608.6 | -617.7 | 352.9 | 389.1 | 22283 | Cistrome | 86928 |
| POLR2A | -999.5 | -1042.7 | 364.9 | 374.5 | 17656 | Cistrome | 45715 |
| POLR2A | -853.3 | -851.4 | 375.3 | 372.0 | 27623 | Cistrome | 68779 |
| POLR2A | -736.6 | -719.4 | 313.2 | 379.1 | 18860 | Cistrome | 83955 |
| POLR2A | -909.4 | -892.4 | 378.8 | 375.7 | 19702 | Cistrome | 49535 |
| POLR2A | -654.6 | -663.4 | 370.6 | 375.4 | 30595 | Cistrome | 88095 |
| POLR2A | -1188.1 | -1177.4 | 338.6 | 371.9 | 9895 | Cistrome | 88098 |
| POLR2A | -758.9 | -742.9 | 362.6 | 368.4 | 26413 | Cistrome | 68785 |
| POLR2A | -583.6 | -584.6 | 391.0 | 377.5 | 25215 | Cistrome | 86927 |
| POLR2A | -768.8 | -742.1 | 336.8 | 373.1 | 19072 | Cistrome | 84236 |
| POLR2A | -967.8 | -1107.6 | 275.6 | 371.3 | 5224 | Cistrome | 82395 |
| POLR2A | -1539.8 | -1491.4 | 341.1 | 356.7 | 7560 | Cistrome | 92088 |
| POLR2A | -837.5 | -882.5 | 347.9 | 365.0 | 21087 | Cistrome | 55671 |
| POLR2A | -765.7 | -777.9 | 354.3 | 367.4 | 22292 | Cistrome | 82336 |

944

| | | | | | | | |
|---|---|---|---|---|---|---|---|
| POLR2A | -891.2 | -904.0 | 383.9 | 360.5 | 26872 | Cistrome | 1152 |
| POLR2A | -996.3 | -1034.1 | 349.9 | 363.9 | 15181 | Cistrome | 83344 |
| POLR2A | -957.7 | -979.2 | 358.0 | 359.2 | 17619 | Cistrome | 83343 |
| POLR2A | -907.3 | -869.2 | 357.8 | 357.3 | 23166 | Cistrome | 68784 |
| POLR2A | -1013.0 | -1026.5 | 338.0 | 362.3 | 11472 | Cistrome | 84237 |
| POLR2A | -1165.9 | -1226.7 | 356.5 | 355.0 | 14701 | Cistrome | 55662 |
| POLR2A | -881.5 | -862.9 | 369.8 | 355.3 | 19125 | Cistrome | 57089 |
| POLR2A | -1232.7 | -1249.3 | 352.9 | 351.3 | 14697 | Cistrome | 55672 |
| POLR2A | -1252.6 | -1323.7 | 323.8 | 345.3 | 9002 | Cistrome | 55663 |
| POLR2A | -1012.4 | -1065.7 | 351.1 | 356.1 | 14058 | Cistrome | 83349 |
| POLR2A | -706.6 | -774.6 | 119.4 | 353.7 | 2841 | Cistrome | 82396 |
| POLR2A | -560.4 | -577.3 | 386.9 | 361.4 | 24672 | Cistrome | 86929 |
| POLR2A | -890.8 | -919.3 | 343.6 | 345.9 | 19957 | Cistrome | 68778 |
| POLR2A | -1254.4 | -1243.7 | 314.6 | 340.0 | 10710 | Cistrome | 55673 |
| POLR2A | -670.3 | -685.3 | 347.7 | 349.6 | 24837 | Cistrome | 84239 |
| POLR2A | -1085.5 | -1124.3 | 331.0 | 343.1 | 14541 | Cistrome | 55675 |
| POLR2A | -1020.0 | -1018.4 | 303.2 | 339.3 | 10930 | Cistrome | 55674 |
| POLR2A | -1108.2 | -1112.5 | 276.7 | 342.3 | 5158 | Cistrome | 82394 |
| POLR2A | -1113.9 | -1170.5 | 271.3 | 341.8 | 6486 | Cistrome | 83350 |
| POLR2A | -1118.6 | -1173.1 | 314.8 | 323.0 | 9147 | Cistrome | 49533 |
| POLR2A | -1182.7 | -1215.6 | 323.8 | 311.9 | 9599 | Cistrome | 49534 |
| POLR2A | -735.2 | -742.2 | 346.2 | 321.7 | 21629 | Cistrome | 89206 |
| POLR2A | -740.6 | -780.3 | 307.3 | 303.9 | 21388 | Cistrome | 1151 |
| POLR2A | -149.5 | -115.9 | 9.6 | 275.3 | 587 | Cistrome | 86619 |
| POLR2A | -242.7 | -267.5 | 39.3 | 189.0 | 1241 | Cistrome | 86354 |
| POLR2A | -133.5 | -135.0 | 34.6 | 127.5 | 429 | Cistrome | 82332 |
| POLR2A | -324.0 | -331.7 | 51.9 | 105.3 | 1387 | Cistrome | 55670 |
| POLR2A | -117.4 | -131.7 | 27.2 | 79.9 | 746 | Cistrome | 92642 |
| POLR2A | -256.4 | -209.0 | 48.4 | 75.1 | 705 | Cistrome | 88092 |
| POLR2A | -143.3 | -129.3 | 33.0 | 83.9 | 681 | Cistrome | 92641 |
| POLR2A | -257.3 | -211.4 | 0.4 | 90.6 | 967 | Cistrome | 55678 |
| POLR2A | -148.5 | -150.0 | 32.1 | 42.5 | 455 | Cistrome | 76192 |
| POLR2A | -109.1 | -91.9 | 14.7 | 16.2 | 485 | Cistrome | 76190 |
| POLR2A | -130.0 | -132.6 | -0.2 | 11.8 | 525 | Cistrome | 55680 |
| POLR2A-5Sp | -720.0 | -699.2 | 347.4 | 478.3 | 40045 | ENCODE | ENCFF910KOG.bed.gz |
| POLR2A-5Sp | -759.6 | -741.0 | 345.8 | 474.6 | 36786 | ENCODE | ENCFF934KIL.bed.gz |
| **DNA accessibility** | | | | | | | |
| ATAC-seq | -462.3 | -474.3 | 398.5 | 384.5 | 61825 | Cistrome | 105339 |
| ATAC-seq | -486.0 | -488.5 | 397.5 | 381.0 | 47323 | Cistrome | 105341 |
| ATAC-seq | -441.0 | -443.5 | 384.2 | 368.7 | 51038 | Cistrome | 105340 |
| ATAC-seq | -488.2 | -493.3 | 377.2 | 358.0 | 43777 | Cistrome | 105338 |
| ATAC-seq | -513.5 | -488.8 | 305.3 | 337.7 | 4594 | Cistrome | 105336 |
| ATAC-seq | -380.7 | -386.0 | 331.3 | 262.2 | 52706 | Cistrome | 66174 |
| ATAC-seq | -347.1 | -351.8 | 284.6 | 187.1 | 38536 | Cistrome | 78866 |
| ATAC-seq | -367.6 | -376.4 | 288.2 | 181.2 | 27142 | Cistrome | 81122 |
| ATAC-seq | -315.4 | -321.2 | 273.2 | 165.5 | 36453 | Cistrome | 81121 |
| ATAC-seq | -298.1 | -308.0 | 266.2 | 151.5 | 40498 | Cistrome | 78867 |
| ATAC-seq | -355.2 | -363.0 | 267.8 | 144.5 | 38154 | Cistrome | 80026 |
| ATAC-seq | -384.5 | -394.7 | 275.3 | 137.0 | 34053 | Cistrome | 80027 |
| ATAC-seq | -346.3 | -352.9 | 264.0 | 135.3 | 34750 | Cistrome | 80024 |
| ATAC-seq | -151.3 | -143.2 | 67.0 | 80.9 | 1432 | Cistrome | 105335 |
| ATAC-seq | -110.6 | -103.7 | 56.4 | 61.4 | 1148 | Cistrome | 105337 |
| ATAC-seq | -90.8 | -93.1 | 18.8 | 18.6 | 844 | Cistrome | 105334 |
| ATAC-seq | -485.2 | -483.4 | 279.5 | 4.5 | 7797 | Cistrome | 80025 |
| DNase | -353.6 | -361.7 | 356.9 | 390.2 | 80101 | Cistrome | 44956 |
| DNase | -347.8 | -356.7 | 359.0 | 384.3 | 78833 | Cistrome | 42166 |
| DNAse | -344.8 | -352.8 | 352.4 | 384.0 | 76277 | Cistrome | 42165 |
| DNase | -383.0 | -388.3 | 356.2 | 372.7 | 67217 | Cistrome | 44955 |
| DNase | -395.6 | -406.8 | 384.5 | 375.1 | 59718 | Cistrome | 42167 |

| | | | | | | | |
|---|---|---|---|---|---|---|---|
| DNAse | -441.3 | -448.1 | 369.6 | 353.1 | 56316 | Cistrome | 42168 |
| DNAseI | -449.9 | -452.3 | 339.4 | 376.3 | 50504 | Ensemble | |
| **Transcription factors** | | | | | | | |
| AFF4 | -143.7 | -147.9 | 14.5 | 65.3 | 1126 | Cistrome | 55639 |
| AFF4 | -30.0 | -25.0 | 0.0 | 50.1 | 483 | Cistrome | 39916 |
| AFF4 | -147.3 | -174.6 | 4.0 | 22.5 | 1331 | Cistrome | 55641 |
| AFF4 | -44.9 | -76.3 | 12.3 | 5.8 | 3112 | Cistrome | 1140 |
| AFF4 | -83.9 | -78.3 | 1.5 | 5.8 | 658 | Cistrome | 55640 |
| AFF4 | -29.8 | -45.9 | 4.4 | 3.4 | 969 | Cistrome | 1141 |
| ARID1A | -588.2 | -614.6 | 430.2 | 455.0 | 46436 | Cistrome | 88976 |
| ARID1A | -603.8 | -616.6 | 404.6 | 410.5 | 21805 | Cistrome | 88981 |
| ARID1A | -35.6 | -34.5 | -0.1 | -3.2 | 1872 | Cistrome | 85741 |
| ARID1A | -46.1 | -41.7 | -23.8 | -52.1 | 768 | Cistrome | 85740 |
| ATF3 | -490.4 | -491.7 | 371.1 | 390.7 | 49476 | Cistrome | 46217 |
| ATF3 | -513.6 | -509.0 | 304.0 | 366.3 | 25474 | Ensemble | homo_sapiens.GRCh38.HCT116.ATF3.SWEmbl_R0005.peaks.20190329.bed.gz |
| ATF3 | -332.9 | -341.7 | 309.8 | 357.3 | 60225 | Cistrome | 68464 |
| ATF3 | -357.8 | -359.5 | 278.0 | 306.7 | 25930 | Cistrome | 68460 |
| ATF3 | -190.4 | -196.7 | 21.0 | 44.0 | 827 | Cistrome | 68463 |
| BANP | -18.4 | -18.9 | 54.2 | 3.3 | 4666 | Cistrome | 71951 |
| BANP | -25.3 | -22.9 | 40.7 | 1.3 | 6183 | Cistrome | 71950 |
| BRD4 | -613.6 | -583.6 | 268.0 | 339.4 | 7401 | Cistrome | 68777 |
| BRD4 | -578.6 | -561.6 | 296.1 | 331.6 | 10531 | Cistrome | 68783 |
| BRD4 | -760.6 | -694.2 | 309.1 | 267.3 | 3725 | Cistrome | 67539 |
| BRD4 | -387.5 | -420.2 | 145.9 | 245.9 | 3745 | Cistrome | 68782 |
| BRD4 | -86.2 | -79.3 | 8.1 | 61.3 | 975 | Cistrome | 68776 |
| BRDU | -138.2 | -126.6 | -94.3 | -124.7 | 734 | Cistrome | 91681 |
| CBX3 | -356.8 | -356.0 | 394.0 | 445.4 | 97142 | Cistrome | 46209 |
| CBX3 | -279.7 | -293.0 | 10.1 | 378.7 | 16531 | Cistrome | 9228 |
| CBX3 | -290.2 | -316.2 | 3.1 | 369.2 | 41056 | Cistrome | 9230 |
| CBX3 | -30.3 | -34.9 | 78.9 | 23.2 | 1278 | Cistrome | 9237 |
| CBX3 | -12.9 | -7.5 | 2.3 | 1.7 | 1491 | Cistrome | 9231 |
| CBX3 | 0.0 | 0.0 | 0.3 | 0.6 | 2594 | Cistrome | 9232 |
| CBX3 | -19.8 | -27.4 | 5.0 | 0.6 | 574 | Cistrome | 9236 |
| CDK9 | -850.4 | -937.0 | 408.6 | 264.1 | 12576 | Cistrome | 57093 |
| CEBPB | -391.6 | -396.3 | 359.2 | 400.2 | 66557 | Cistrome | 46206 |
| CEBPB | -418.3 | -424.2 | 289.8 | 375.1 | 30413 | Ensemble | homo_sapiens.GRCh38.HCT116.CEBPB.SWEmbl_R0005.peaks.20190329.bed.gz |
| CENPA_liftOver | -352.5 | -362.7 | -666.9 | -657.0 | 1056 | | CENPA_GSM1105684_2fold_enriched_merged_hg38.bed.gz |
| CENPA-HuRef* | -432.3 | -442.1 | -889.4 | -866.3 | 1598 | Cistrome | 40153 |
| CNOT3 | -775.1 | -786.6 | 475.4 | 320.1 | 47784 | Cistrome | 76964 |
| CTCF | -312.7 | -327.0 | 332.9 | 330.5 | 54768 | Cistrome | 46218 |
| CTCF | -328.8 | -345.7 | 326.9 | 312.4 | 54059 | Cistrome | 85285 |
| CTCF | -288.7 | -304.2 | 320.6 | 302.2 | 49581 | ENCODE | ENCFF850PKJ.bed.gz |
| CTCF | -321.1 | -339.5 | 316.6 | 300.8 | 41280 | Cistrome | 85284 |
| CTCF | -250.6 | -263.6 | 307.5 | 300.2 | 58981 | ENCODE | ENCFF518MQA.bed.gz |
| CTCF | -286.0 | -297.8 | 324.1 | 303.6 | 36079 | Cistrome | 42151 |
| CTCF | -278.8 | -292.7 | 319.9 | 300.5 | 52155 | ENCODE | ENCFF171SNH.bed.gz |
| CTCF | -293.2 | -308.2 | 316.9 | 300.2 | 29057 | Cistrome | 72779 |
| CTCF | -330.7 | -332.7 | 314.9 | 300.0 | 23805 | Cistrome | 85286 |
| CTCF | -249.6 | -259.3 | 314.7 | 292.3 | 51750 | Cistrome | 42152 |
| CTCF | -294.1 | -308.1 | 307.9 | 294.1 | 42363 | Cistrome | 45717 |
| CTCF | -294.1 | -307.8 | 307.7 | 294.0 | 42385 | Cistrome | 42148 |
| CTCF | -255.2 | -273.0 | 323.5 | 293.0 | 30116 | ENCODE | ENCFF056ESE.bed.gz |
| CTCF | -269.1 | -284.5 | 303.6 | 285.8 | 52042 | Cistrome | 42149 |
| CTCF | -268.9 | -284.1 | 303.5 | 285.7 | 52021 | Cistrome | 45716 |

| | | | | | | | |
|---|---|---|---|---|---|---|---|
| CTCF | -273.9 | -287.0 | 316.6 | 283.8 | 45381 | Cistrome | 42154 |
| CTCF | -242.8 | -255.4 | 306.0 | 283.0 | 45369 | Cistrome | 42150 |
| CTCF | -239.7 | -250.6 | 315.9 | 284.4 | 35571 | ENCODE | ENCFF549PGC.bed.gz |
| CTCF | -269.5 | -285.7 | 317.5 | 283.0 | 49084 | Cistrome | 42153 |
| CTCF | -274.6 | -291.0 | 277.8 | 264.4 | 50595 | Enseble | homo_sapiens.GRCh38.HCT116.CTCF.SWEmbl_R0005.peaks.20190329.bed.gz |
| EGR1 | -584.6 | -587.7 | 403.0 | 409.9 | 47145 | Cistrome | 46214 |
| Egr1 | -800.6 | -780.5 | 341.8 | 393.0 | 17084 | Ensemble | homo_sapiens.GRCh38.HCT116.Egr1.SWEmbl_R0005.peaks.20190329.bed.gz |
| ELF1 | -554.5 | -563.6 | 393.0 | 400.8 | 45158 | Cistrome | 46202 |
| ELF1 | -679.7 | -685.1 | 323.6 | 385.4 | 24133 | Ensemble | homo_sapiens.GRCh38.HCT116.ELF1.SWEmbl_R0005.peaks.20190329.bed.gz |
| ELL2 | -599.4 | -606.4 | 230.5 | 356.5 | 4198 | Cistrome | 55645 |
| ELL2 | -261.2 | -250.9 | 61.6 | 139.3 | 1977 | Cistrome | 55647 |
| ELL2 | -157.2 | -170.0 | 46.3 | 89.6 | 1387 | Cistrome | 55646 |
| ELL2 | -152.7 | -191.3 | 39.3 | 69.0 | 1098 | Cistrome | 55648 |
| ELL2 | -133.0 | -153.0 | 28.9 | 26.5 | 1683 | Cistrome | 1143 |
| ELL2 | -30.5 | -23.5 | 1.1 | 8.2 | 1036 | Cistrome | 1144 |
| EP300 | -259.9 | -250.9 | 250.5 | 273.8 | 19798 | Cistrome | 42907 |
| EZH2 | -69.4 | -60.1 | 0.1 | -1.8 | 672 | Cistrome | 101554 |
| EZH2 | -25.1 | -22.1 | 2.8 | -5.1 | 660 | ENCODE | ENCFF806ESQ.bed.gz |
| EZH2 | -18.9 | -9.4 | 0.9 | -7.1 | 867 | ENCODE | ENCFF926EZW.bed.gz |
| EZH2 | -94.9 | -87.3 | -21.5 | -53.6 | 616 | Cistrome | 101555 |
| EZH2 | -265.8 | -315.2 | -11.0 | -240.6 | 5222 | Ensemble | homo_sapiens.GRCh38.HCT116.EZH2.SWEmbl_R0005.peaks.20190329.bed.gz |
| EZH2-T487p | -71.9 | -84.8 | -36.8 | -91.4 | 560 | Cistrome | 100246 |
| EZH2-T487p | -4.6 | -4.1 | 7.5 | -0.5 | 425 | ENCODE | ENCFF325ZGB.bed.gz |
| FOSL1 | -388.8 | -397.4 | 366.3 | 401.9 | 70141 | Cistrome | 46212 |
| FOSL1 | -376.0 | -376.1 | 279.6 | 352.2 | 31315 | Ensemble | homo_sapiens.GRCh38.HCT116.FOSL1.SWEmbl_R0005.peaks.20190329.bed.gz |
| HEXIM1 | -593.4 | -631.4 | 460.5 | 325.7 | 55397 | Cistrome | 57092 |
| HNF4A | -214.7 | -193.7 | 227.0 | 348.6 | 3800 | Cistrome | 54666 |
| HNF4A | -390.9 | -391.6 | 275.2 | 340.9 | 25783 | Cistrome | 54673 |
| HNF4A | -311.9 | -269.2 | 294.5 | 336.6 | 4471 | Cistrome | 54665 |
| HNF4A | -396.1 | -416.5 | 272.3 | 331.0 | 15595 | Cistrome | 54674 |
| HNF4A | -132.9 | -109.2 | 137.3 | 241.2 | 2286 | Cistrome | 54660 |
| HSF1 | -115.8 | -119.3 | 2.6 | 9.0 | 851 | Cistrome | 51108 |
| HSF1 | -128.9 | -109.9 | 2.6 | 6.9 | 772 | Cistrome | 51110 |
| HSF1 | -80.6 | -117.7 | -0.4 | 0.1 | 486 | Cistrome | 51107 |
| ICE1 | -473.8 | -469.4 | 356.2 | 365.1 | 41898 | Cistrome | 39920 |
| ICE1 | -443.4 | -451.1 | 349.3 | 356.7 | 39819 | Cistrome | 39914 |
| ICE1 | -59.9 | -49.5 | 10.1 | 50.1 | 548 | Cistrome | 39919 |
| ICE2 | -123.2 | -92.9 | 173.9 | 204.1 | 2316 | Cistrome | 39924 |
| JUND | -420.8 | -427.6 | 377.0 | 412.3 | 69908 | Cistrome | 46213 |
| Jund | -440.3 | -437.6 | 291.0 | 375.5 | 31993 | Ensemble | homo_sapiens.GRCh38.HCT116.Jund.SWEmbl_R0005.peaks.20190329.bed.gz |
| JUND | -260.8 | -254.5 | 222.5 | 275.0 | 23965 | ENCODE | ENCFF998KDQ.bed.gz |
| JUND | -261.3 | -261.6 | 221.2 | 269.5 | 21907 | ENCODE | ENCFF333VCK.bed.gz |
| KAT2B | -153.6 | -109.7 | -21.5 | -57.4 | 728 | Cistrome | 85025 |
| KDM3B | -412.9 | -401.7 | 313.6 | 310.7 | 15346 | Cistrome | 82127 |
| KDM3B | -289.4 | -262.0 | 337.5 | 312.9 | 5043 | Cistrome | 82126 |
| KDM5A | -1055.0 | -1298.9 | 146.8 | 363.5 | 4675 | Cistrome | 92366 |
| KDM5A | -1036.4 | -1241.3 | 173.6 | 358.0 | 5429 | Cistrome | 92367 |

| | | | | | | | |
|---|---|---|---|---|---|---|---|
| KMT2B | -260.0 | -255.2 | 262.6 | 299.3 | 21353 | Cistrome | 42906 |
| LARP7 | -530.1 | -568.0 | 422.3 | 255.8 | 39326 | Cistrome | 57091 |
| LEO1 | -331.4 | -244.1 | 16.0 | 242.7 | 1500 | Cistrome | 55661 |
| MAX | -504.0 | -512.3 | 417.9 | 446.1 | 74266 | Cistrome | 46216 |
| Max | -719.8 | -735.1 | 380.4 | 438.6 | 33616 | Ensemble | homo_sapiens.GRCh38.HCT116.Max.SWEmbl_R0005.peaks.20190329.bed.gz |
| MCM2 | -322.6 | -334.4 | 242.2 | 148.3 | 65110 | Cistrome | 83886 |
| MECP2 | -458.7 | -465.9 | 316.3 | 322.7 | 34007 | Cistrome | 34399 |
| MECP2 | -310.3 | -342.2 | 255.3 | 239.1 | 8828 | Cistrome | 34400 |
| MTA2 | -59.9 | -47.3 | -37.4 | -77.1 | 474 | Cistrome | 87604 |
| MYC | -1065.3 | -1051.0 | 358.0 | 361.9 | 13446 | Cistrome | 70809 |
| MYC | -1008.2 | -1035.8 | 371.7 | 343.1 | 15844 | Cistrome | 70807 |
| MYC | -888.2 | -879.5 | 404.6 | 328.1 | 17904 | Cistrome | 70806 |
| MYC | -1150.6 | -1141.9 | 381.5 | 318.3 | 8287 | Cistrome | 70805 |
| MYC | -966.3 | -940.6 | 426.7 | 301.3 | 11599 | Cistrome | 70810 |
| MYC | -501.7 | -538.9 | 357.7 | 301.3 | 48637 | Cistrome | 70808 |
| MYC | -407.9 | -412.4 | 163.6 | 107.5 | 1306 | Cistrome | 76186 |
| NIPBL | -578.5 | -575.8 | 357.0 | 355.4 | 29832 | Cistrome | 84953 |
| NIPBL | -570.7 | -562.9 | 346.5 | 349.8 | 28604 | Cistrome | 88450 |
| NR0B2 | -133.1 | -118.3 | 4.6 | 33.5 | 603 | Cistrome | 55660 |
| PAF1 | -713.0 | -671.5 | 279.0 | 411.6 | 9761 | Cistrome | 88906 |
| PAF1 | -716.2 | -735.7 | 365.6 | 389.7 | 26835 | Cistrome | 55667 |
| PAF1 | -738.2 | -733.7 | 334.2 | 365.9 | 20824 | Cistrome | 84915 |
| PAF1 | -1236.5 | -1214.6 | 182.7 | 347.1 | 5762 | Cistrome | 55666 |
| PAF1 | -1061.6 | -1066.0 | 339.7 | 348.1 | 15947 | Cistrome | 55668 |
| PAF1 | -1017.9 | -1019.0 | 328.0 | 341.5 | 16156 | Cistrome | 55664 |
| PAF1 | -1261.5 | -1233.2 | 311.9 | 335.7 | 10352 | Cistrome | 55669 |
| PAF1 | -1180.3 | -1262.1 | 278.7 | 327.8 | 7895 | Cistrome | 55665 |
| PHIP | -487.5 | -491.6 | 400.5 | 386.7 | 41242 | Cistrome | 85463 |
| PHIP | -500.7 | -507.5 | 380.4 | 355.8 | 36060 | Cistrome | 85459 |
| PHIP | -591.6 | -612.5 | 396.1 | 344.7 | 15366 | Cistrome | 85462 |
| PHIP | -489.6 | -585.1 | 391.1 | 326.7 | 3676 | Cistrome | 87886 |
| PHIP | -911.1 | -977.0 | 388.1 | 326.1 | 7263 | Cistrome | 87884 |
| PHIP | -849.7 | -885.9 | 386.2 | 317.4 | 4636 | Cistrome | 85464 |
| PHIP | -191.3 | -176.5 | 11.3 | 67.8 | 891 | Cistrome | 85465 |
| RAD21 | -333.5 | -343.7 | 365.2 | 380.4 | 91714 | Cistrome | 46207 |
| RAD21 | -709.2 | -760.2 | 430.4 | 384.7 | 67731 | Cistrome | 71320 |
| RAD21 | -669.5 | -713.1 | 439.5 | 373.6 | 98855 | Cistrome | 71322 |
| RAD21 | -439.8 | -379.1 | 380.1 | 365.7 | 6718 | Cistrome | 85290 |
| RAD21 | -644.1 | -698.2 | 419.1 | 348.2 | 90949 | Cistrome | 71321 |
| Rad21 | -314.6 | -326.4 | 316.7 | 329.1 | 49942 | Ensemble | homo_sapiens.GRCh38.HCT116.Rad21.SWEmbl_R0005.peaks.20190329.bed.gz |
| RAD21 | -615.9 | -664.5 | 384.9 | 307.1 | 84470 | Cistrome | 71319 |
| RAD21 | -302.7 | -313.5 | 306.2 | 295.6 | 36976 | Cistrome | 81366 |
| REST | -500.8 | -511.1 | 313.2 | 323.1 | 14210 | Cistrome | 46203 |
| REST | -126.0 | -132.9 | 124.9 | 55.6 | 3483 | Ensemble | homo_sapiens.GRCh38.HCT116.REST.SWEmbl_R0005.peaks.20190329.bed.gz |
| SIN3A | -710.8 | -704.0 | 337.4 | 413.0 | 27435 | Ensemble | homo_sapiens.GRCh38.HCT116.SIN3A.SWEmbl_R0005.peaks.20190329.bed.gz |
| SIN3A | -637.4 | -636.8 | 370.8 | 380.2 | 33224 | Cistrome | 46208 |
| SIRT1 | -684.0 | -706.3 | 334.6 | 252.3 | 39329 | Cistrome | 83885 |
| SKP2 | -636.7 | -641.8 | 348.0 | 277.2 | 100000 | Cistrome | 93781 |
| SMARCA4 | -152.2 | -114.3 | 107.0 | 242.1 | 7586 | Cistrome | 72102 |
| SMARCA4 | -22.5 | -19.3 | 0.1 | 9.2 | 767 | Cistrome | 72100 |
| SMARCC1 | -87.6 | -74.9 | 49.7 | 149.3 | 6118 | Cistrome | 72101 |

| | | | | | | | |
|---|---|---|---|---|---|---|---|
| SMARCC1 | -2.2 | -0.8 | 0.4 | 13.9 | 1081 | Cistrome | 72099 |
| SMC1A | -454.4 | -433.1 | 331.4 | 324.4 | 7426 | Cistrome | 85288 |
| SMC1A | -273.8 | -286.0 | 305.1 | 285.3 | 43636 | Cistrome | 85289 |
| SP1 | -1040.6 | -1141.5 | 597.6 | 500.9 | 82625 | Cistrome | 5395 |
| SP1 | -441.7 | -443.3 | 391.5 | 427.3 | 72471 | Cistrome | 46215 |
| SP1 | -525.7 | -524.2 | 324.3 | 401.0 | 34242 | Ensemble | homo_sapiens.GRCh38.HCT116.SP1.SWEmbl_R0005.peaks.20190329.bed.gz |
| SP1 | -1147.5 | -1179.0 | 420.6 | 387.6 | 21322 | Cistrome | 70813 |
| SP1 | -1078.5 | -1143.4 | 452.5 | 386.6 | 24694 | Cistrome | 70811 |
| SP1 | -1255.1 | -1387.0 | 438.2 | 382.0 | 19784 | Cistrome | 70812 |
| SP1 | -433.7 | -462.8 | 415.5 | 376.0 | 100000 | Cistrome | 70814 |
| SP1 | -557.1 | -581.9 | 421.5 | 304.1 | 30938 | Cistrome | 70815 |
| SP1 | -455.9 | -494.8 | 346.9 | 297.7 | 46179 | Cistrome | 70816 |
| SRF | -463.2 | -464.5 | 379.4 | 402.6 | 57748 | Cistrome | 46211 |
| Srf | -581.9 | -563.4 | 321.5 | 399.9 | 28421 | Ensemble | homo_sapiens.GRCh38.HCT116.Srf.SWEmbl_R0005.peaks.20190329.bed.gz |
| TCF4 | -30.6 | -16.5 | 47.3 | 55.8 | 3003 | Cistrome | 54661 |
| TCF4 | -26.4 | -32.5 | 33.4 | 43.5 | 1508 | Cistrome | 54669 |
| TCF4 | -27.3 | -17.1 | 34.7 | 36.1 | 1898 | Cistrome | 54664 |
| TCF4 | -17.4 | -12.8 | 27.0 | 34.8 | 1990 | Cistrome | 54667 |
| TCF4 | -35.6 | -23.6 | 23.4 | 28.0 | 1825 | Cistrome | 54663 |
| TCF4 | -15.3 | -18.3 | 14.4 | 19.9 | 1043 | Cistrome | 54670 |
| TCF4 | -17.3 | -10.2 | 10.2 | 16.0 | 872 | Cistrome | 54662 |
| TCF7L1 | -597.7 | -645.1 | 420.3 | 347.0 | 47979 | Cistrome | 70804 |
| TCF7L1 | -362.2 | -372.5 | 326.1 | 269.1 | 11321 | Cistrome | 70803 |
| TCF7L1 | -219.6 | -218.8 | 241.5 | 267.7 | 8191 | Cistrome | 70802 |
| TCF7L2 | -428.4 | -417.7 | 357.1 | 340.0 | 45234 | Cistrome | 45714 |
| TCF7L2 | -472.8 | -460.2 | 328.2 | 323.2 | 20454 | ENCODE | ENCFF199EHQ.bed.gz |
| TCF7L2 | -461.7 | -450.0 | 328.0 | 314.1 | 15431 | ENCODE | ENCFF736EVD.bed.gz |
| TCF7L2 | -347.6 | -280.3 | 290.4 | 288.7 | 4946 | Ensemble | homo_sapiens.GRCh38.HCT116.TCF7L2.SWEmbl_R0005.peaks.20190329.bed.gz |
| TEAD4 | -449.3 | -450.6 | 340.1 | 360.2 | 39773 | Cistrome | 46210 |
| TEAD4 | -423.1 | -412.1 | 251.5 | 301.6 | 11796 | Ensemble | homo_sapiens.GRCh38.HCT116.TEAD4.SWEmbl_R0005.peaks.20190329.bed.gz |
| TET2 | -691.8 | -748.7 | 160.5 | 267.0 | 3570 | Cistrome | 50981 |
| TOP1 | -339.6 | -310.6 | 309.6 | 266.6 | 11177 | Cistrome | 68774 |
| TOP1 | -183.9 | -183.2 | 264.0 | 236.9 | 7423 | Cistrome | 68781 |
| TOP1 | -168.5 | -183.5 | 302.1 | 168.6 | 4395 | Cistrome | 68775 |
| TOP1 | -255.0 | -246.6 | 321.9 | 123.7 | 3755 | Cistrome | 68780 |
| TP53 | -78.2 | -78.6 | 214.1 | 236.6 | 9393 | Cistrome | 82544 |
| TP53 | -21.2 | -23.3 | 45.2 | 47.4 | 3353 | Cistrome | 50345 |
| TP53 | -33.6 | -30.0 | 22.2 | 18.6 | 1004 | Cistrome | 92110 |
| TP53 | -14.4 | -15.4 | 29.7 | 12.5 | 781 | Cistrome | 53285 |
| TP53 | -17.9 | -17.6 | 30.4 | 12.5 | 717 | Cistrome | 53286 |
| TP53 | -24.1 | -24.5 | 13.8 | 13.1 | 1368 | Cistrome | 82545 |
| TP53 | -14.0 | -17.8 | 32.7 | 5.9 | 1900 | Cistrome | 68462 |
| TP53 | -15.8 | -19.2 | 2.2 | 0.2 | 807 | Cistrome | 50344 |
| TP53 | -5.0 | -7.5 | 5.9 | 0.0 | 880 | Cistrome | 68461 |
| TRIM28 | -799.4 | -878.3 | 408.5 | 258.4 | 24209 | Cistrome | 57090 |
| USF1 | -474.1 | -476.9 | 384.6 | 397.2 | 57530 | Cistrome | 46205 |
| USF1 | -703.1 | -689.1 | 332.1 | 387.6 | 9985 | Ensemble | homo_sapiens.GRCh38.HCT116.USF1.SWEmbl_R0005.peaks.20190329.bed.gz |
| USP49 | -134.5 | -131.3 | 108.1 | 75.3 | 2124 | Cistrome | 39345 |
| USP49 | -99.3 | -109.3 | 16.8 | 5.8 | 1350 | Cistrome | 39347 |
